# Engineering the mechanosensitivity of single DNA molecules via high-throughput microfluidic force spectroscopy

**DOI:** 10.64898/2026.02.24.707783

**Authors:** Matthew P. DeJong, Yujia Bian, Jennifer E. Ortiz-Cárdenas, Bianca Figueroa, Ananya Pant, Eduardo Posadas-Barrera, Lillian Brixi, Magnus S. Bauer, Alexander R. Dunn, Polly M. Fordyce

**Affiliations:** Department of Chemical Engineering, Stanford University, Stanford, CA, USA; Department of Genetics, Stanford University, Stanford, CA, USA; Stanford Microfluidics Foundry, Stanford University, Stanford, CA, USA; Stanford Cardiovascular Institute, Stanford University School of Medicine, Stanford, CA, USA; Department of Bioengineering, Stanford University, Stanford, CA, USA; Sarafan ChEM-H Institute, Stanford University, Stanford, CA, USA; Chan Zuckerberg Biohub, San Francisco, CA, USA

## Abstract

Single-molecule force spectroscopy (SMFS) is a powerful tool to measure how biomolecules respond to mechanical force, but limited sequence throughput constrains its potential. Here, we present a single-molecule, multiplexed, microfluidic force spectroscopy (SM^3^FS) assay that uses parallelized microfluidics to measure up to 80 sequence variants per experiment. Using SM^3^FS, we stretched, overstretched, or unzipped 241 different DNA structures and generated 131,847 single-molecule traces from a quarter-million observations. High sequence throughput allowed us to identify DNA structures that are kinetically stable yet mechanically fragile (F_rupture_ < 3 pN at 0.5 pN s^-1^), revealing how mechanosensitivity can arise as an intrinsic property of multivalent systems. By enabling systematic sequence-function mapping under force, SM^3^FS opens a path to high-throughput nonequilibrium studies of biomolecules.

## Introduction

Understanding and engineering mechanical biomolecules would substantially benefit from techniques that systematically generate many variants and measure their behavior under mechanical load.^1,2^ High-throughput biomolecular engineering tools have revolutionized our ability to understand how biomolecular sequence encodes function through folded structure,^3,4^ stability,^5,6^ and interaction landscapes.^7–9^ Nearly all these methods make measurements in the absence of force, yet processes like antibody maturation,^10,11^ T-cell signaling,^12,13^ bacterial pathogenesis,^14,15^ cell migration,^16^ sensing (touch, hearing, proprioception),^17^ homeostasis (blood pressure, tissue maintenance),^18^ and embryonic morphogenesis^19^ rely on molecular force sensing and occur far from thermodynamic equilibrium.

While fluctuation theorems can calculate equilibrium properties from nonequilibrium information,^20–22^ these theorems cannot calculate the reverse because nonequilibrium properties are pathway-dependent. For example, the rupture force of DNA duplexes changes from ∼12 pN to >30 pN when pulled in unzipping versus shearing geometries, despite sharing the same equilibrium binding constant^23^; changing the pulling direction selects a new reaction pathway and consequently dramatically changes nonequilibrium properties. Thus, existing high-throughput tools that measure properties at equilibrium are fundamentally limited in their ability to investigate the nonequilibrium processes that drive life.

Single-molecule force spectroscopy (SMFS) applies calibrated forces to molecules and measures their resulting motions (e.g. unfolding, stretching, dissociation) and can therefore quantify nonequilibrium properties such as transition state distances, force sensitivities, and dynamics.^24,25^ The quantitative understanding of DNA duplex mechanics resulting from SMFS measurements^26,27^ led to the design of DNA-based sensors for imaging and engineering cellular force transduction,^28–31^ as well as DNA-based catch bonds that strengthen with force.^32,33^ However, because nonequilibrium properties are challenging to predict, understanding quickly diminishes for molecules more complex than a standard DNA duplex.

Recent technological advances using microfluidic force spectroscopy,^34,35^ acoustic force spectroscopy,^36^ magnetic tweezers,^37,38^ and centrifugal force spectroscopy^39,40^ have increased *measurement* throughput: the number of single molecules that can be measured in an experiment (**Supplementary Table 1**). Nevertheless, these technologies remain constrained in *sequence* throughput, measuring one sequence variant per experiment. Conversely, recent cell-based assays expand *sequence* throughput, the number of individual biomolecule sequences characterized in a single experiment, by linking sequence to cellular mechanotransduction.^41–45^ However, existing cell-based approaches do not directly measure the properties of single molecules and therefore cannot quantify force-dependent kinetics required to understand the molecular mechanisms that underlie force sensing. Thus, there is a need for scalable, direct *in vitro* approaches to understand and engineer the force-dependent properties of biomolecules.^1^

Here, we developed SM^3^FS, a tool for high-throughput, multiplexed force spectroscopy assays. SM^3^FS uses microfluidic force spectroscopy^34,35^ to apply tension forces ranging from < 0.25 to 100 pN to single molecules tethered between a surface and beads. Wide-field imaging and optimized surface chemistry make it possible to track up to 18,000 beads in parallel and resolve molecular transitions down to 2 nm. SM^3^FS employs three multiplexing strategies (spatial, mechanical, and sequential) to scale SMFS assays and measure up to 80 DNA duplex variants per experiment. Using 10 microfluidic devices, we stretched, overstretched, or unzipped 241 variants of DNA duplexes, measuring a total of 131,847 single-molecule traces from 249,148 beads. These measurements of DNA sequence variants revealed how mechanosensitivity can arise as an intrinsic nonequilibrium property of multivalent systems. These and other measurements demonstrate how the throughput of SM^3^FS can enable mechanistic insights that are not readily obtained using conventional single-molecule techniques.

## Results

### Microfluidic force spectroscopy is a simple, precise, and scalable SMFS assay

SM^3^FS builds upon previously developed bead-based microfluidic force spectroscopy assays^34,35,46–48^ in which beads are tethered to surfaces via a single molecule and subjected to calibrated drag forces from fluid flow (**Figs. 1A,B**). Here, we pattern the flow chambers with a monomeric anti-digoxigenin antibody fragment (Fab) that binds a DNA duplex labeled with digoxigenin and biotin on opposite 5’ ends such that a single DNA duplex can tether streptavidin-coated polystyrene beads to the device surface (**Fig. 1A**). A programmable pressure controller controls fluid flow to the channels, thereby applying drag forces on the bead (**Fig. 1B**). The resulting tension on the tether (*F_T_*) depends on this drag force (*F_D_*) and the angle formed between the device surface and the tether (*α*) (**Eqn. 1 Fig. 1C**):

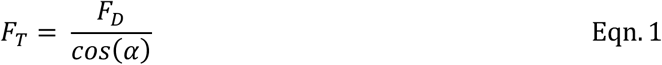

**Figure 1.**
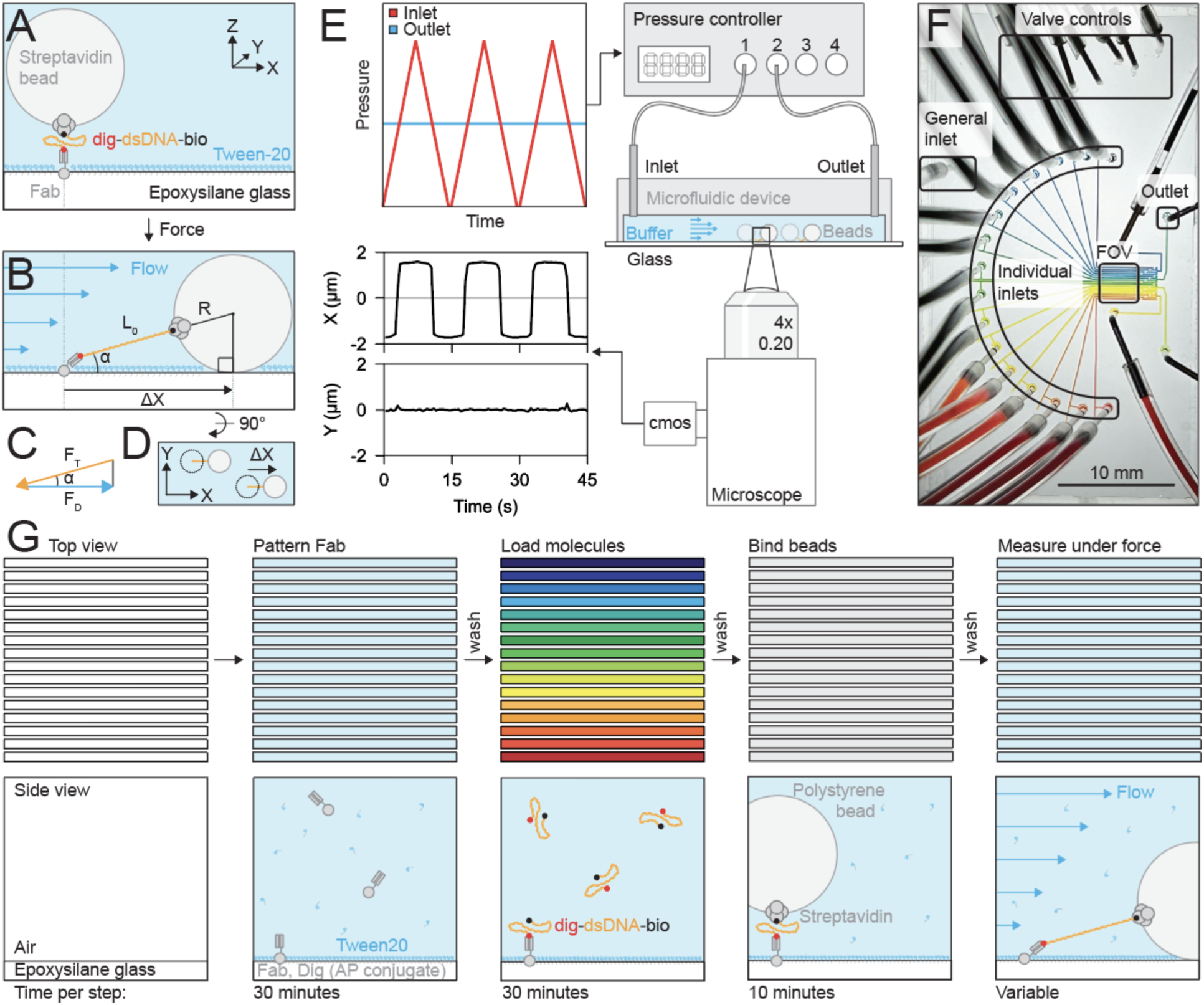
Multiplexed microfluidics for parallelized surface preparation, force application, and measurements. **A.** An anti-digoxigenin Fab immobilizes dsDNA that is labeled with digoxigenin and biotin on opposite 5’ ends. The dsDNA tether recruits a streptavidin-coated bead. **B.** Flow exerts drag on the bead and tension on the tether. X-position displacements (Δ*X*) are measured in response to force. **C.** The molecular geometry (*⍺*) of the bead and tether determines the relationship between tension and drag force. **D.** X-Y particle tracking simultaneously measures many molecules under force. **E.** Microfluidic force spectroscopy varies pressure-driven flow to exert force on tethered beads and measures their response by tracking X-Y position. **F.** Image of a multichannel microfluidic device for SM^3^FS indicating positions of a general inlet, valve controls, individual channel inlets, a shared image area, and outlets. **G.** Top and side view schematics of SM^3^FS workflow (see text).

As flow within microfluidic channels is laminar (maximum Re < 0.5), the drag force on the bead follows Stokes’ law and is proportional to the flow velocity (*v*) and bead radius (*R*) (*F*_*D*_ ∝ *R* · *v*). The bead experiences shear flow at the channel wall, such that for a given wall shear rate, *γ_w_*, the flow velocity at the center of a bead on the surface also depends on the bead radius (*v* = *γ_w_* · *R*).^49^ Both flow rate and shear rate also increase linearly with flow pressure: *v* ∝ *γ_w_* ∝ *P*. Combining terms, the drag force on the tethered bead scales linearly with the flow pressure and with the square of the bead radius: *F*_*D*_ ∝ *R* · *v* ∝ *γ_w_* · *R*^2^ ∝ *P* · *R*^2^, resulting in a lower bead-to-bead error in force than methods with forces that scale with bead volume (∼*R^3^*) such as acoustic-,^36^ magnetic-,^37^ and centrifugal-force based assays^39^ (**Supplementary Table 1**).

Tracking the position of the bead over time allows the measurement of single-molecule stretching, unfolding, and unbinding events (**Fig. 1D**). Here, we use a standard inverted microscope to monitor displacements of micron-sized beads in response to flow forces (**Fig. 1E**). To assess the spatial resolution of our experimental setup, we tracked immobilized 1-μm fluorescent beads. These measurements yielded X-Y resolutions of 20 and 2 nm for 4x and 100x objectives, respectively (**Supplementary Fig. 1**). Unlike other image-based approaches that require Z-position measurements for particle tracking, X-Y particle tracking enables widefield imaging and the quantification of force-dependent responses for over 10,000 single molecules in one field of view, more than other bead-based SMFS platforms reported to date (**Supplementary Table 1**).

### A multichannel device spatially multiplexes microfluidic force spectroscopy assays

To expand the sequence throughput of microfluidic force spectroscopy assays, we engineered a valved multichannel device that positions 16 parallel channels (150 µm × 3,330 µm) within a 3.33 × 3.33 mm field of view (**Fig. 1F, Supplementary Fig. 2**). A shared inlet tree connects all 16 channels and allows parallel introduction of reagents required for surface functionalization, tethering beads to immobilized molecules, and applying drag forces (**Figs. 1F,G**). Channel-specific valves allow introduction of reagents to a single channel at a time and prevent cross-contamination between channels (**Supplementary Figs. 2,3**).

To begin assays, we bond valved multichannel SM^3^FS devices to epoxysilane-coated glass slides (**Fig. 1G**). Next, we fill channels with anti-digoxigenin Fab suspended in a Tween-20 surfactant solution to reduce nonspecific sticking and control the surface density of Fab molecules. After washing channels with a Tween-20 solution to further passivate the channel surfaces, we load molecular tethers comprised of biotin- and digoxigenin-labeled dsDNA. After incubating to allow labeled DNA duplexes to attach, we introduce streptavidin-coated beads. Once patterning is complete, we use pressure-driven flow to apply drag force (*F_D_*) to the beads and tension force (*F_T_*) to their tethers (**Fig. 1C**).

### Multiplexed assays identify optimal surface functionalization conditions for high-throughput single-molecule measurements

Efficient measurement of force-dependent displacements for many molecules requires minimizing non-specific sticking while maximizing the number of singly tethered beads. Typical SMFS experiments use sparse attachment densities to enrich for primarily single-molecule measurements, a factor that limits experimental throughput. To gain insight into how the surface density of randomly placed DNA attachment points determines the number of single-molecule tethers, we performed Monte Carlo simulations in which we randomly placed 4-kbp DNA and 1-µm beads in a 20 × 20 µm grid, assumed binding between any colocalized tethers and beads, and counted the number of beads bound to one or multiple DNA tethers (**Supplementary Fig. 4**). In the limit of few surface-attached DNA molecules, essentially all beads had a single DNA tether. As the number of DNA molecules increased, the number of beads with a single DNA tether initially increased before decreasing to zero as beads with multiple DNA tethers predominated. Thus, if single-tether beads can be experimentally distinguished from multiple-tether beads, moderate surface attachment densities should substantially increase single-molecule measurement throughput.

To test this hypothesis, we experimentally varied the ratio of Fab to Tween-20 passivating surfactant and quantified single-molecule attachment. Specifically, we coated channels with varying v/v percentages of Fab (15 concentrations, 0% to 100% v/v Fab solution in PBS with 0.1% Tween-20), added digoxigenin- and biotin-labeled 4,000-bp DNA tethers (or buffer alone as a negative control), bound 1-μm beads, and then oscillated the direction of flow while monitoring bead displacements in both X and Y (**Figs. 2A-C**). In the absence of DNA, non-specific sticking was minimal, with low numbers of beads observed at even the highest Fab concentrations. In the presence of DNA, the number of detected beads dramatically increased, confirming specific attachment via DNA tethers (**Fig. 2A**).

**Figure 2.**
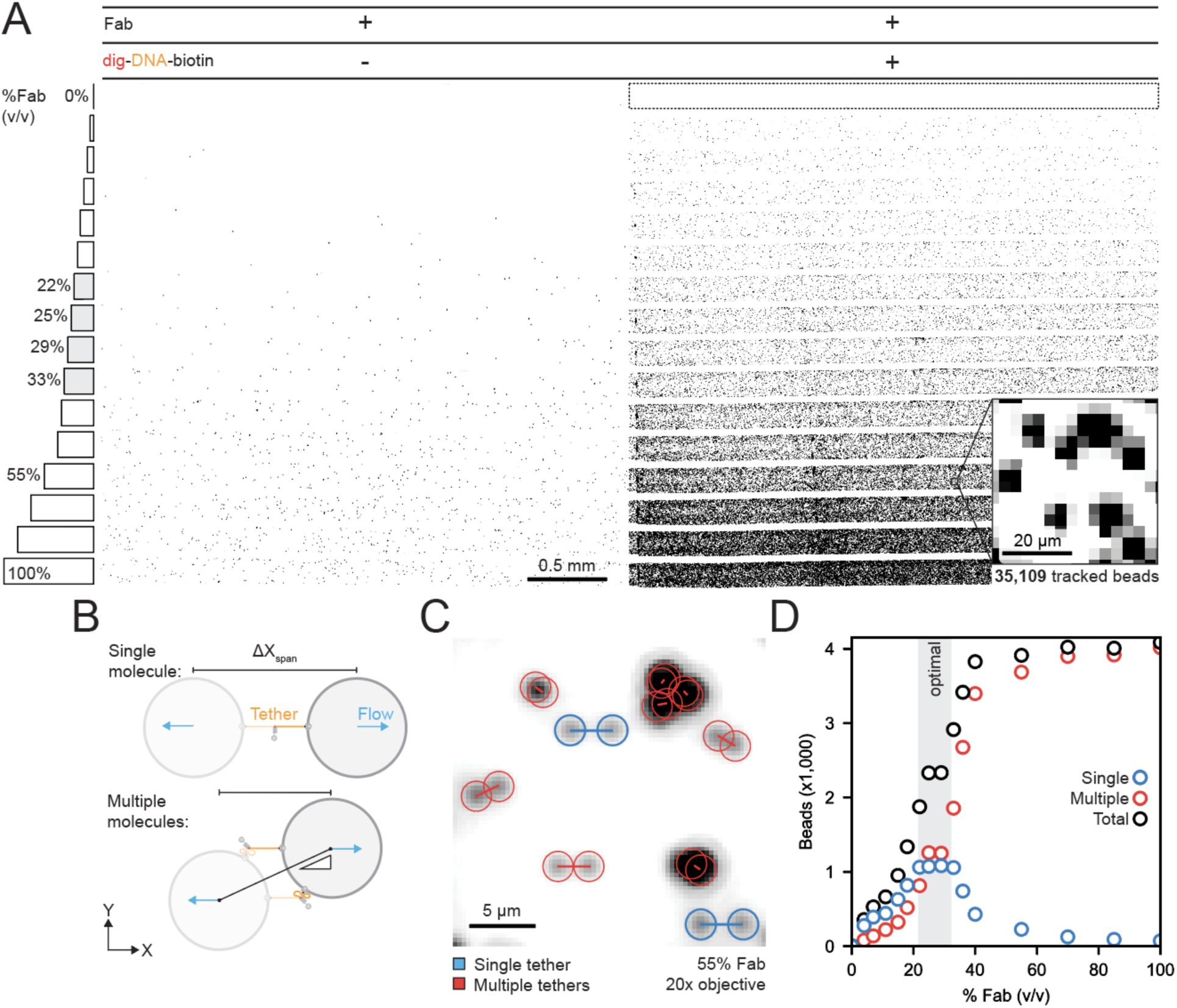
Optimizing surface preparation for single-molecule tethers. **A.** Inverted fluorescence images of green fluorescent 1-µm beads within SM3FS devices without (*left*) and with (*right*) dsDNA tethers; each channel was coated with a different % anti-dig Fab. A box indicates the negative control channel; shaded Fab % labels indicate the range of densities typically used in subsequent experiments. The expanded inset shows the region highlighted in 2C. **B.** Schematic illustrating how length and angle of bead displacements in a flow reversal assay differentiates beads bound by single- or multi-molecule tethers. **C.** Representative summed projection image from a flow reversal assay for the region highlighted in 2A illustrating movements for beads bound by single- or multi-molecule tethers (blue and red, respectively). **D**. Number of beads attached to single (*blue*) vs. multiple (*red*) tethers as a function of relative Fab volume fractions; *grey* shaded bar indicates relative Fab volume fractions of 22 to 33% that yield an optimal density of single-molecule tethers; measurements identified 7,962 total single DNA tethers.

Beads tethered by a single molecule should transit a maximum expected displacement of 3,580 nm parallel to the direction of flow, corresponding to the full extension of the 4-kbp dsDNA. Beads tethered by multiple molecules should transit a shorter distance and rotate between attachment points with a perpendicular component to their displacements (**Fig. 2B**). Observed trajectories were consistent with these expectations (**Fig. 2C, Supplementary Fig. 5**), making it possible to clearly identify singly tethered beads. At low Fab concentrations, all beads exhibited trajectories consistent with single tethers. As Fab concentrations increased, the absolute number of singly tethered beads increased and then decreased, consistent with Monte Carlo predictions (**Fig. 2D**, **Supplementary Fig. 5**). Overall, Fab concentrations between 22 and 33% yielded approximately equal numbers of singly and multiply tethered beads (∼1,100 per channel) and maximized the density of single molecules, making it possible to measure ∼18,000 single-molecule tethers in a 16-channel device.

### Parallelized dsDNA stretching measurements demonstrate spatial multiplexing and relate applied pressure to tensions below 25 pN

Applying calibrated tension via bead-based assays requires accurately relating applied pressures to exerted tension. Physiologically relevant tensions range from < 5 pN (corresponding to the activation force of Notch, the force generated by myosin II, and the forces experienced by key proteins within cellular adhesion complexes)^50–54^ to > 50 pN, corresponding to the peak forces experienced by integrins.^28,55^ We leveraged the well-characterized nanomechanical properties of dsDNA to calibrate pressure to drag and tension forces at low forces by measuring force-extension curves, which require low 10s of pN of tension,^47,56^ for 15 dsDNA duplexes of varying lengths (500-7500 bp; **Supplementary Fig. 6**).

To obtain force-extension curves for all 15 dsDNA duplexes in parallel, we introduced each construct (and a buffer-only negative control) to separate channels within the device and then added 1-μm beads to form tethers. The negative control channel bound only one bead, indicating minimal cross contamination (**Fig. 3A**, bottom lane); the 7500 bp construct did not amplify successfully and was excluded from further analysis (**Supplementary Fig. 6**). Upon applying pressure-driven oscillating flows, we simultaneously tracked 22,268 beads. Inspection of individual bead trajectories established that 10,984 were attached via single-molecule tethers (**Figs. 3A,B; Supplementary Fig. 7**). The distances traversed by each singly tethered bead increased across channels, consistent with expected increasing contour lengths (**Fig. 3C, Supplementary Fig. 7**). For dsDNA duplexes with faint alternate bands visible by gel electrophoresis, distance distributions revealed rare populations (typically < 2 mass %) with shorter contour lengths, highlighting the sensitivity of SM^3^FS for detecting and quantifying molecular heterogeneity (**Supplementary Fig. 8**).

**Figure 3.**
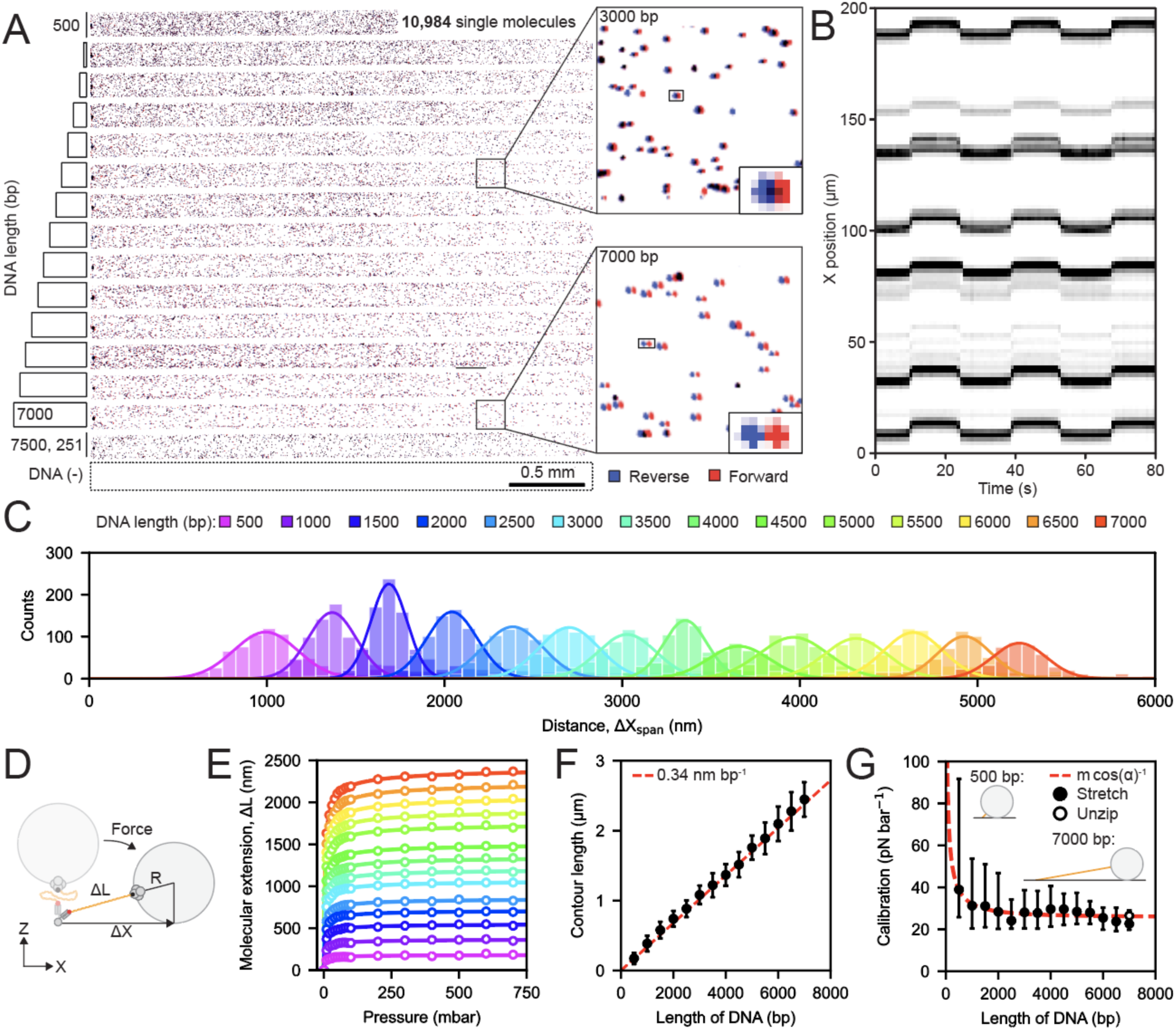
Parallelized DNA force-response measurements at scale. **A.** Inverted fluorescence images of green-fluorescent 1-µm beads within an SM^3^FS device patterned with dsDNA tethers ranging from 500 to 7000 bp. Bead positions under forward and reverse flow are shown in *red* and *blue*; a box delineates the negative control channel and panels at right show zoomed-in representative images. A line shows the region highlighted in 3B. **B.** Kymograph of *X* position over time for 12 representative beads attached via 6,000-bp tethers from the region indicated in 3A. **C.** Histograms showing the distribution of bead displacements during a flow reversal assay for each tether length; histograms show information for 10,984 single-molecule tethers. **D.** Schematic illustrating bead displacement (*ΔX*) and molecular extension (*ΔL*) upon application of flow force. **E.** Measured DNA extension as a function of applied pressure for each dsDNA tether length tethered to 1-µm Dynabeads. Markers indicate the mean extension calculated from a Gaussian fit; solid lines show WLC fit to these data. **F.** Mean contour lengths returned from the WLC fit as a function of DNA tether length (in bp). Error bars denote standard deviations; the red dashed line indicates the expected contour length. **G.** The tension-pressure calibration as a function of DNA tether length (in bp). The calibration represents the slope, *a*, of the linear tension-pressure relationship: *F_T_* (pN) = *a* * *P* (mbar). The red-dashed line was the tension-pressure relationship from the force balance, *a = m*·cos(*⍺*)^-1^, where *m* is the drag force-pressure calibration (**Supplementary Note 1**) and *⍺* is determined via the geometric model shown in (Fig. 1B). Error bars indicate the interquartile range of the calibration. The calibration constant *m* was calculated from a least-squares fit. Open circle indicates independent measurement of unzipping for a 15-bp duplex attached to a 7-kbp tether. Inset schematic shows a 1-micron bead attached to a 500- or 7000-bp tether, drawn to scale.

Next, we incrementally increased flow pressure, quantified bead displacements across all channels at each pressure, and calculated molecular extensions (*ΔL*) from bead displacement (*ΔX*) for each dsDNA tether assuming a simple geometric model 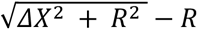; **Fig. 3D, Supplementary Fig. 9**). Single-molecule extension traces were well-fit with a worm-like-chain (WLC) model^57^ (**Fig. 3E, Supplementary Figs. 10-12**) with fitted contour lengths increasing linearly with the size of the DNA tether, as expected (**Fig. 3F**). The ratio of the fitted and theoretical persistence lengths yielded tension-to-pressure calibrations across molecular geometries and the expected, linear pressure-to-drag force relationship (**Fig. 3G, Supplementary Note 1**).^58,59^

To confirm the accuracy of our pressure-force calibration, we performed an additional experiment in which we tethered beads to surfaces via dsDNA constructs containing a 15-bp DNA duplex^23^ known to unzip at 10.3 pN (**Supplementary Fig. 13**). One strand of the 15-bp duplex was displayed at the end of a 7-kbp tether and the other was attached to the bead such that unzipping of the duplex led to irreversible bead loss. Beads ruptured at an average pressure of 340 ± 30 mbar, corresponding to an estimated tension of 9 ± 1 pN when using the force/pressure calibration derived above (**Fig. 3G, Supplementary Fig. 13**). This additional calibration confirmed that applied tensions can be accurately determined from applied pressures in a low force regime ranging 0.25 to 25 pN.

### Increasing bead size extends the range of applied tensions to over 100 pN

To assess the capability of our device to measure force responses beyond 25 pN, we relied on the fact that B-form dsDNA undergoes a cooperative transition to an overstretched, 70% longer conformation (S-form dsDNA) at ∼65 pN (**Fig. 4A**).^60^ As drag force scales with R^2^, we increased the bead diameter from 1 to 3 μm to access order-of-magnitude larger forces. Using 3-μm beads, the 65 pN overstretching transition was clearly visible as a secondary displacement beyond the maximum expected displacement for B-form DNA (**Figs. 4B,C; Supplementary Fig. 14**). Dig/anti-dig ruptures at lower forces than the DNA B-to-S transition^35^ such that most (84%) single molecules ruptured from the surface before overstretching; however, the remaining 603 (of 3,742) single-molecule tethers underwent a detectable B-to-S transition (**Fig. 4D, Supplementary Figs. 15-19**). Across tether lengths, tethers transitioned in length at a drag force corresponding to ∼65 pN of tension force according to the geometric model (**Eqn. 1** and **Supplementary Fig. 20**). Measured drag forces exceeded 100 pN for some tethers, demonstrating that SM^3^FS can access tension forces beyond 100 pN.

**Figure 4.**
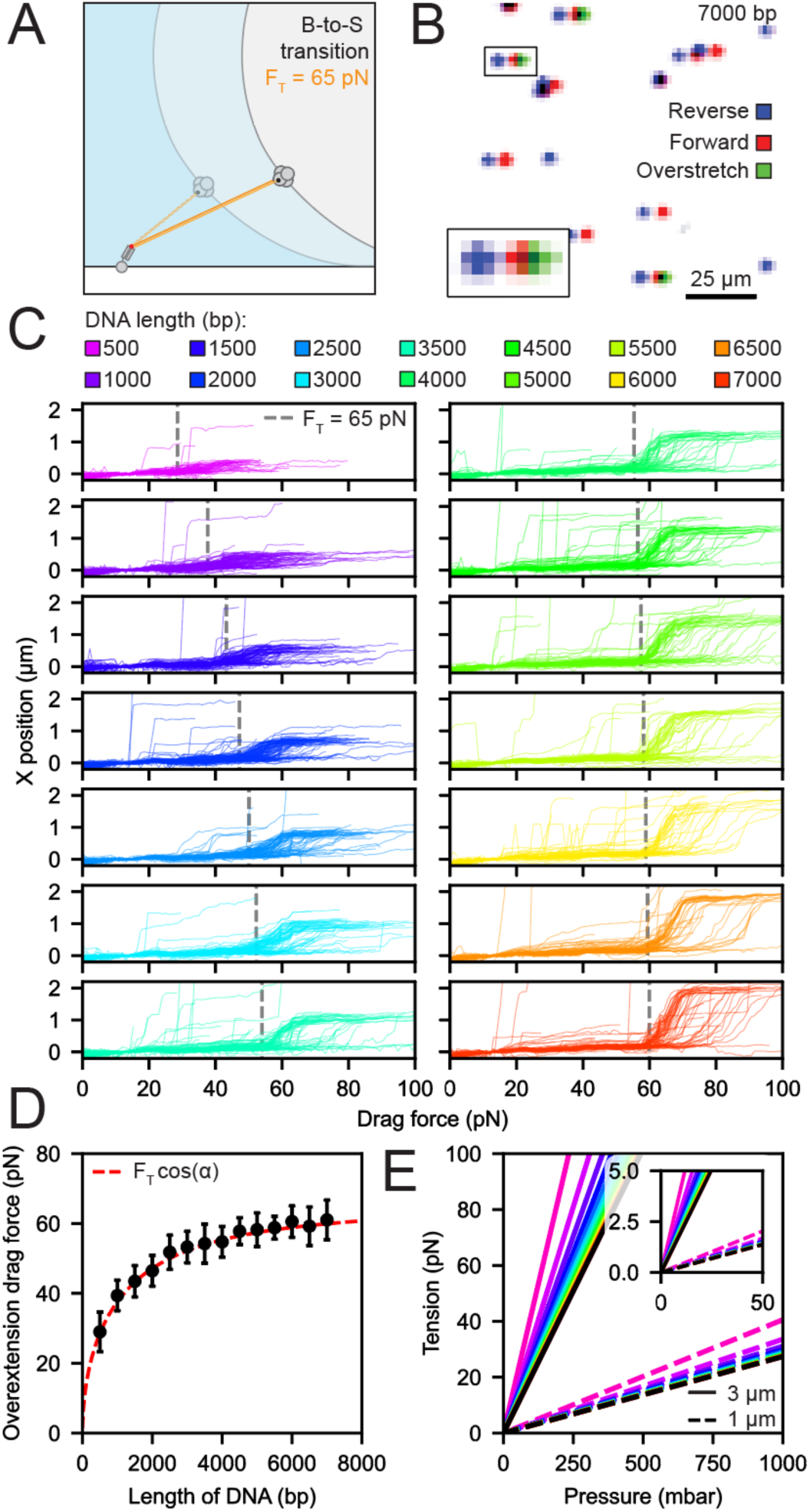
Precise force-response measurements past 100 pN. **A.** Schematic illustrating the expected DNA overextension and associated bead displacement for applied tension >65 pN. **B.** Bright-field image overlays showing representative bead positions under reverse flow (*blue*), initial forward extension (*red*), and overextension (*green*) for 7,000-bp tethers. **C.** Measured *X* position as a function of drag force for 3,742 single-molecule tethers of varying lengths. Grey dashed lines indicate when the expected tension corresponds to 65 pN. **D.** Calculated drag forces at the overextension transition based on a sudden molecular extension beyond the contour length; markers denote the mean, error bars denote standard deviations derived from a Gaussian fit. The dashed red line indicates predictions from the geometric model, where F_T_ is the overextension tension of 65 pN. **E.** Calibrated pressure-tension relationships for 1- and 3-micron beads. Dashed and solid lines indicate calibrations for 1- and 3-micron beads, respectively. Lines are colored by tether length as in 4C. Black lines denote the drag force-pressure relationship in the limit of long tethers (**Supplementary note 2**).

Together, these measurements of DNA stretching and overstretching confirm that: (1) the relationship between applied pressure and drag force is linear across a wide range of pressures, and (2) a simple geometric model is sufficient to predict molecular tension forces resulting from applied pressure. While torque should, in theory, rotate the bead and hence alter the force balance between drag force and tension^61^, the with-torque model decreased agreement with experimental observations across all tested molecular geometries (**Supplementary Fig. 21**), possibly because a balancing torque from static friction opposes hydrodynamic torque (**Supplementary Note 2, Supplementary Figs. 22-25**). Together, these DNA stretching and overstretching calibration experiments establish the ability to apply forces to molecular tethers ranging from 0.25 to 100 pN using 1- and 3-micron beads and pressures accessible with a commercially available pressure control system (**Fig. 4E**).

### Mechanical and sequential multiplexing expands throughput

Profiling >16 different sequences requires measurements across microfluidic devices that may have slightly different calibrations between pressure and applied force due to small differences in tubing length or channel dimensions that alter overall resistance. To increase measurement accuracy across devices, we included an internal standard in each channel. Specifically, we employed a mechanical multiplexing approach in which we patterned each channel with: (1) a constant 15-bp ‘fiducial’ duplex (previously shown to unzip and rupture at 10.3 ± 0.5 pN at a ramp rate of 11 pN s^-1^)^23^, and (2) a ‘variable’ duplex of interest, each tethered with linkers of different lengths (3,000 and 7,000 bp) such that they could be unambiguously identified by the distance traversed upon application of flow (**Figs. 3C** and **5A**). As with prior unzipping experiments (**Supplementary Fig. 13**), duplex unzipping led to irreversible bead loss, providing the opportunity to sequentially multiplex measurements by iteratively hybridizing new sequences (**Fig. 5A**).

**Figure 5.**
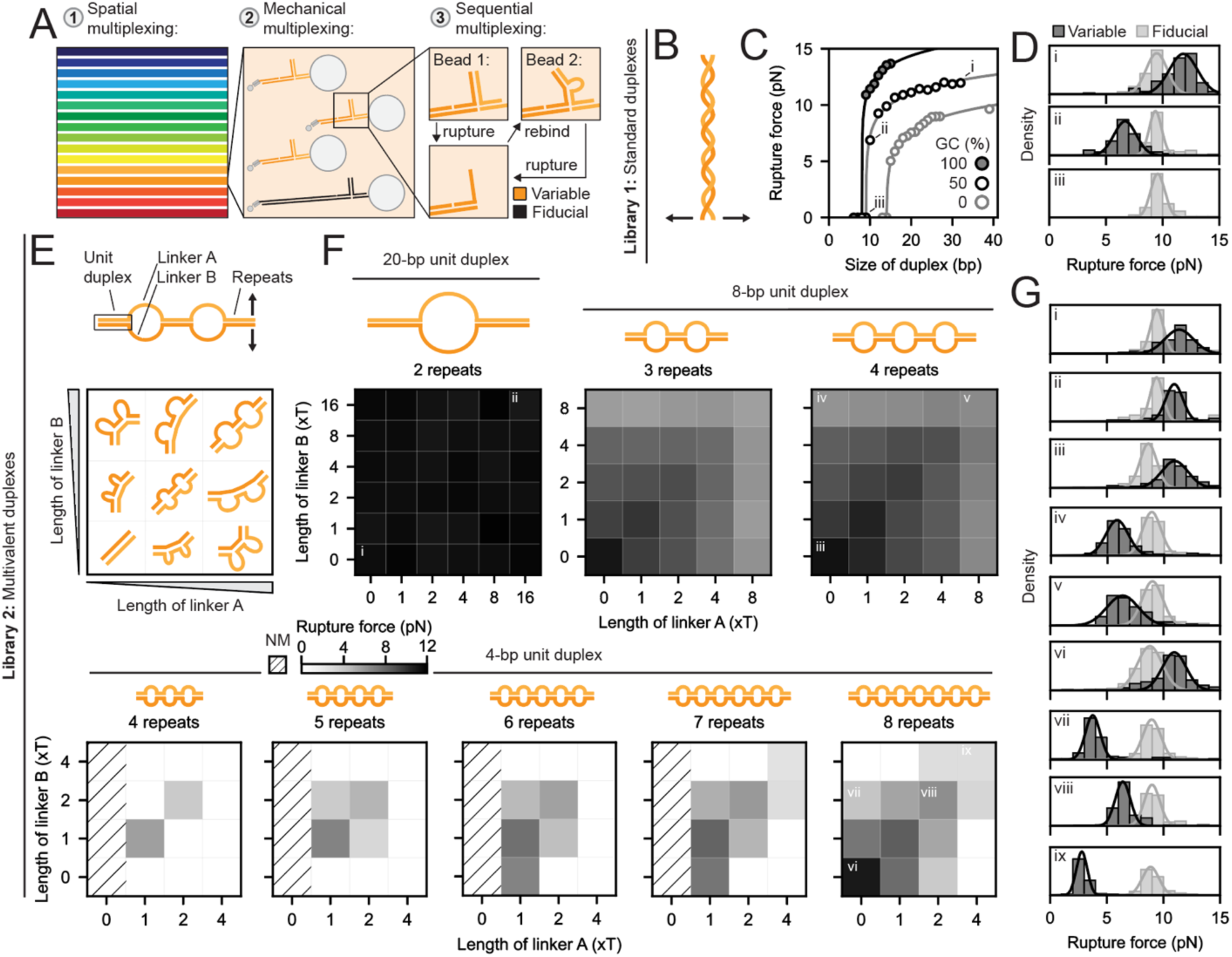
Engineering DNA duplexes to decouple mechanical strength from thermodynamic stability. **A.** Schematic illustrating unzipping assays employing spatial multiplexing (*i.e.* patterning a different surface-attached ‘tether’ sequence in each channel), mechanical multiplexing (*i.e.* including ‘fiducial’ sequences with known rupture forces in each channel on long tethers that can be uniquely identified by extension length), and sequential multiplexing (*i.e.* iteratively unzipping and hybridizing different sequences to surface-attached ‘tether’ strands). **B.** Standard DNA duplex pulled in an unzipping geometry. **C.** Measured unzipping forces for standard DNA duplexes of varying lengths and GC contents. The rupture force distributions for points *i*-*iii* are highlighted in 5D. The trend lines are fit to an empirical logarithmic model (see Methods). The force ramp rate was 0.5 pN s^-1^. **D.** Rupture force distributions for the standard duplexes highlighted in 5C. Light and dark grey indicate fiducial and variable DNA duplexes, respectively. **E.** Multivalent DNA duplexes consist of repeats of unit duplexes linked by poly-T linkers A and B. The linker size is held constant for a given strand but can differ for A and B, introducing potential strain. **F.** Rupture force matrices for multivalent duplexes. A schematic for each matrix illustrates the multivalent duplex with the largest, equal-length linkers. Dark grey indicates larger unzipping forces. White indicates no measurable rupture events. Duplexes with dashed boxes were not measured. The rupture force distributions for points *i*-*ix* are highlighted in 5G. The force ramp rate was 0.5 pN s^-1^. **G.** Unzipping force histograms for multivalent duplexes highlighted in 5F. Light and dark grey indicate fiducial and variable DNA duplexes, respectively.

### A mechanically encoded fiducial enables efficient ratiometric rupture force measurements

To assess the mechanical and sequential multiplexing strategy, we iteratively quantified rupture forces for the same 15-bp fiducial duplex on both 3-kbp and 7-kbp tethers. In a single field of view, we measured 18,378 single-molecule duplex unzipping events from 31,533 total beads, averaging 1,148 single molecules per channel (**Supplementary Figs. 26, 27**); three iterative rounds of hybridization and unzipping yielded 47,188 single-molecule events from 75,435 total tracked beads in 30 minutes. Beads typically bound in the same location upon loading and hybridization, establishing the ability to repeatedly measure rupture from a single immobilized DNA strand (**Supplementary Fig. 26**). Measured contour lengths revealed two distinct populations of molecules corresponding to the 3- and 7-kbp tethers (**Supplementary Fig. 27**). At a ramp rate of ∼0.5 pN s^-1^ (corresponding to a pressure ramp rate of 1 bar min^-1^, **Fig. 3G**), the 15-bp fiducial duplex unzipped at 9.3 ± 0.6 pN (S.D.; N = 1,347 molecules), in agreement with expectation for the slower ramp rate (**Supplementary Figs. 28, 29**). The time to rupture varied across channels and devices, reflecting small differences in pressure-to-force relationships (3% and 5% CV between channels and devices, respectively, **Supplementary Figs. 27, 30**). Normalizing the rupture force of the duplex attached to the 3-kbp tether by the same duplex on the 7-kbp tether reduced the per channel variance to 0.8%, enabling precise ratiometric unzipping measurements (**Supplementary Fig. 27**).

### High-throughput force spectroscopy of DNA sequence variant libraries identifies multivalent constructs that unzip at low forces

Many important force-responsive cellular processes experience small, ∼3 pN, molecular forces.^50–54^ However, published DNA tension sensors^28^ based on irreversible duplex unzipping rupture at ∼ 10 pN and therefore do not access this biologically relevant, low-force regime. Inspired by the multivalent architecture of highly sensitive, naturally occurring force sensors such as Notch and chromatin, we hypothesized that a sticker–spacer architecture in which short, weakly interacting DNA duplexes are connected by flexible linkers could achieve avidity-driven stability while still unzipping at low forces because each individual duplex remains intrinsically weak. To test this idea, we designed and measured the mechanical strengths of two libraries of DNA duplexes.

We first used SM^3^FS to measure the unzipping forces of a library of standard DNA duplexes without linkers (**Fig. 5B**). To quantify the lower limit of mechanical strength of DNA, we quantified unzipping forces for 40 DNA duplexes with 0, 50, or 100% GC content and lengths ranging from 8 to 39 base pairs. As the length of the hybridized DNA duplex increased, the unzipping force increased and approached 12 ± 2, 14 ± 3, and 18 ± 5 pN for long (100 bp) duplexes with 0, 50%, or 100% GC content, respectively, in agreement with past unzipping force measurements of ∼10 and 20 pN for pure A-T and G-C duplexes, respectively (**Figs. 5B-D, Supplementary Figs. 31-33**).^62,63^ Below a critical length of 15, 10, or 9 bp for GC contents of 0%, 50%, or 100%, respectively, duplexes were unstable and did not tether beads to surfaces (**Supplementary Fig. 31-34, Supplemental Note 4**). Duplexes with the minimal length required for tethering unzipped at 5 ± 1 pN, 7 ± 1 pN, and 11 ± 2 pN for 0%, 50%, or 100% GC content, respectively. These measurements establish a new lower limit for forces that can be sensed by standard DNA duplexes.

Next, we quantified sequence-strength relationships for a second library of DNA duplex variants that systematically varied sticker affinity, linker length, and the number of sticker/linker repeats. Multivalent DNA variants were comprised of 2-8 hybridized duplexes of varying length (4, 8, or 20 bp with 50% GC content) separated by varying lengths (0 to 16 nt) of flexible polyT linkers, yielding 186 variants in total (**Fig. 5E**). Iteratively introducing and hybridizing bead-tethered oligonucleotides with different length linkers created dsDNA duplexes with either symmetric bubbles or asymmetric bulges, with asymmetric bulges introducing strain in the multivalent duplex (**Fig. 5E**).

Rupture force matrices for the 20-, 8-, and 4-bp constructs were symmetric, indicating that bead vs. surface immobilization did not impact quantification and that measurements were reproducible (**Fig. 5F, Supplementary Figs. 35-37**). For the largest (20-bp) concatenated unit duplexes, measured rupture forces were similar to their non-concatenated counterparts, with little dependence on linker composition (**Figs. 5F,G i and ii**). As unit duplex length decreased, measured rupture forces increasingly diverged for different numbers of repeats, and linker-induced strain had a greater impact (**Figs. 5F,G iii-ix**). Few tethers were observed for 3 to 6 repeats of the shortest (4-bp) unit duplex, suggesting that these constructs are not thermodynamically stable; no rupture events were detected for 2 repeats of the 8-mer duplex, likely due to formation of a secondary structure element that prevented hybridization **(Supplementary Figs. 38** and **39**). The number of detected tethers increased for 7 repeats of the (4-bp) unit duplex, and tethers were abundant for 8 repeats, suggesting that these multivalent DNA structures were stable (**Supplementary Fig. 37**). Increasing the number of 4-bp repeats only marginally increased rupture force such that a 7 × (4-bp + 4T) duplex ruptured at 2.4 ± 0.4 and an 8 × (4-bp + 4T) duplex ruptured at 2.7 ± 0.5 pN. To our knowledge, these values make unzipping of the (4-bp + 4T) constructs one of the most force-sensitive dissociation processes ever reported.

### Evidence of and mechanistic basis for decoupling mechanical strength from thermodynamic stability

As a successful measurement of a rupture event first requires formation of a thermodynamically stable interaction between the two halves of the duplex structure, the number of observed tethers should provide qualitative information about thermodynamic stability. Consistent with this, the number of observed tethers increased with DNA duplex length (0, 53, and 97 tethers for 8-, 10-, and 32-bp 50% GC standard DNA duplexes (**Supplementary Fig. 32**) and number of repeated units for multivalent duplexes (4, 27, and 105 for 6 ×, 7 ×, and 8 × (4-bp + 4T) repeats (**Supplementary Fig. 37**)). Importantly, many observed rupture events of the 8 × (4-bp + 4T) duplex suggested a thermodynamically stable interaction despite rupturing below 3 pN.

To assess the effect of thermodynamic stability on mechanical strength, we computationally estimated the free energy of hybridization for all measured duplexes.^64^ For short, standard DNA duplexes, hybridization free energies correlated with unzipping force, as expected (shaded region, **Fig. 6A**, **Supplementary Fig. 40**). For multivalent duplexes, the calculated free energies of hybridization indicated that duplexes with matched linker lengths and sufficient repeats were thermodynamically stable, in line with our experiments. Increasing linker lengths progressively decoupled mechanical strength from thermodynamic stability to yield stable dsDNA constructs that unzipped at very low forces (**Fig. 6A**, left) and mismatched linker lengths introduced strain that further reduced mechanical strength (**Fig. 6A**, right). Despite sharing similar free energies of hybridization, the 8 × (4-bp + 4T) duplex ruptured at a ∼60% lower force than the 10-bp standard DNA duplex (**Fig. 6B**). Thus, multivalency provides a potential strategy to decouple mechanical strength and thermodynamic stability.

**Figure 6.**
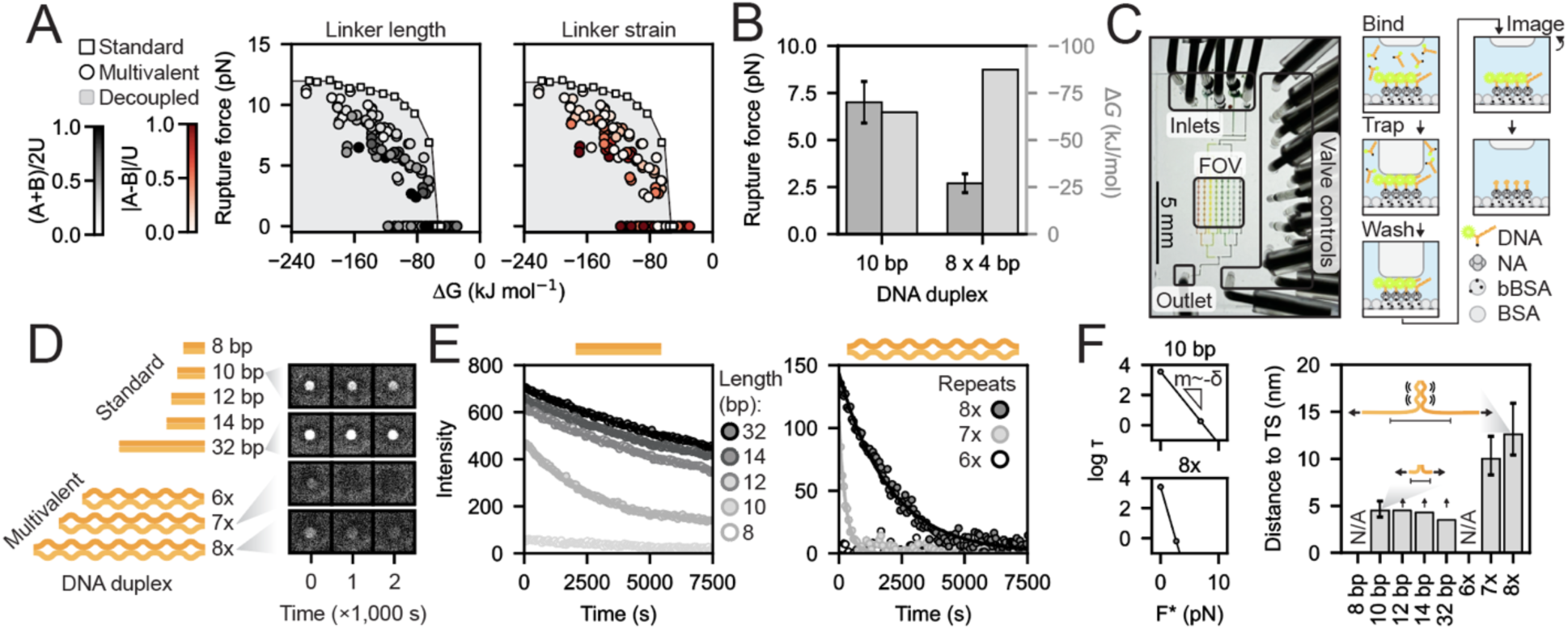
Mechanistic basis for decoupling mechanical strength from thermodynamic stability. **A.** Measured most probable unzipping force vs. free energy of hybridization at 21°C predicted with DuplexFold.^64^ Squares indicate standard duplexes with 50% GC content. Circles indicate all measured multivalent duplexes with 4- or 8-bp unit duplexes colored by length of linkers (A + B) or strain of linkers |A – B| relative to the length of the unit duplex, U, in *grey* (left) and *red* (right). The grey shaded region indicates the region in which mechanical strength is weaker than expected relative to the thermodynamic stability of standard DNA. **B.** Measured most probable rupture unzipping force and predicted free energy of hybridization for a 10-bp standard duplex and the 8 x (4 bp + 4T) multivalent duplex. Error bars represent the standard deviation of measured rupture forces. **C.** Image of microfluidic device (left) and schematic of experimental protocol (right) for miniaturized MITOMI assays. Device includes multiple inlets, valve controls, a shared image area, and an outlet; experimental protocol involves sequentially loading unique duplexes into each row of channels via a binary inlet tree, trapping bound duplexes, and washing away unbound duplexes. The dissociation of surface-immobilized duplexes is imaged with flow to prevent rebinding. DNA is labeled with fluorescein and a biotin for imaging and immobilization, respectively. **D.** Example background-subtracted fluorescent images for 32- and 10-bp duplexes with 50% GC content and the 8× and 7×4-bp repeat construct with 4T linkers. **E.** Dissociation curves (intensity *vs.* time) for fluorescently-labeled standard DNA duplexes with 50% GC content (*left*) and for multivalent 4-bp + 4T duplexes (*right*). Intensity is calculated from average, background-corrected fluorescence intensity from three chambers. **F.** Log bond lifetime *τ* versus dissociation force for the 10-bp and 8 x (4 bp + 4T) duplexes. The zero-force bond lifetime was estimated from the dissociation measurements in 6E. The estimated bond lifetime at the most probable rupture force was numerically estimated from the Bell-Evans model (**Supplementary note 5**). The distances to the transition state *δ* are proportional to the slope *m*. Arrows denote a lower bound imposed by photobleaching in 6E. Error bars denote the associated error propagated from the standard deviation of the measured unzipping force. Cartoons depict transition state models for the 10-bp standard duplex and 8 x (4 bp + 4T) duplex.

As computational free energy predictions may not be accurate for DNA duplexes with large, atypical bulges,^65^ we also measured bulk, zero-force dissociation rates for a subset of DNA duplexes via a new miniaturized assay based on mechanically induced trapping of molecular interactions (miniMITOMI)^66,67^ (**Fig. 6C, Supplementary Fig. 41**). Briefly, we immobilized DNA duplexes modified with fluorescein and biotin on opposite strands on a neutravidin-coated surface and measured the decay of the fluorescent signal under continuous flow to quantify zero-force dissociation kinetics of the two strands (**Figs. 6C,D**). 12, 14, and 32 bp DNA duplexes with 50% GC content remained stably bound (*τ* > 50,000 ± 20,000 s) (**Fig. 6E**). Dissociation of the 10-bp duplex was substantially faster (*τ*_0_ = 3,600 ± 200 s) and thus directly measurable in this assay. The 8-bp duplex was not stable in this assay, consistent with the lack of observed rupture events from the 8-bp condition in the SM^3^FS assay.

For 6×, 7×, and 8 × (4-bp + 4T) multivalent duplex variant constructs, kinetic stability increased with number of unit duplex repeats (**Figs. 6D**, **E**). The 7-repeat construct rapidly dissociated (*τ*_0_ = 280 ± 20 s), while dissociation for the 8-repeat construct was ∼10-fold slower (*τ*_0_ = 2,600 ± 200 s). We note that the 8 × (4-bp + 4T) construct ruptured at 2.7 pN compared to 6.9 pN for a 50% GC 10-bp duplex despite having similar zero-force dissociation rates. These direct measurements of the thermal off rate provide experimental evidence that multivalency can decouple mechanical strength from stability.

The rate of a reaction *k_f_* (here, duplex dissociation) under force *F* depends on the change in extension *δ* that a molecule undergoes in going from the ground to transition state:

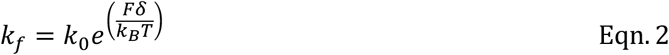

where *k_0_* is the rate in the absence of force, *k_B_* is Boltzmann’s constant, and *T* is temperature.^68^ We used force-dependent (**Fig. 5F**) and zero-load dissociation rates (**Fig. 6E**) to calculate *δ* for various duplex structures (**Supplementary Fig. 42, Supplementary Note 5**) ^69,70^. The distance to the transition state for unzipping the 10-bp, 50% GC duplex was 4.5 ± 1.0 nm, consistent with the literature value of 5.1 ± 0.4 nm, suggesting that unzipping ∼7bp of the duplex leads to irreversible rupture.^26^ In contrast, distances to the transition state for the 8 × and 7 × (4-bp + 4T) duplexes were 13 ± 3 nm and 10 ± 2 nm, respectively (**Fig. 6G**). Accounting for the elasticity of the unzipped multivalent duplexes below 3 pN, these distances suggest that unzipping 3 to 4 unit duplexes is sufficient to drive irreversible rupture (**Fig. 6H, Supplementary Fig. 43, Supplementary Note 6**). Therefore, multivalency independently tunes the height of and distance to the transition state of dissociation, offering a flexible strategy for mechanosensing.

## Discussion

Here, we developed a new microfluidic platform, SM^3^FS, for high-throughput force spectroscopy of sequence variant libraries through advances in single-molecule immobilization and multiplexing. By varying bead size and flow rate, we established that SM^3^FS can apply forces from 0.25 pN to 100 pN, enabling future application to established microfluidic force spectroscopy assays of DNA-associated enzymes,^34,71,72^ cytoskeletal motors,^73^ and protein-protein interactions.^14,46,74^

We leveraged the sequence multiplexing capabilities of SM^3^FS to systematically map sequence-mechanics relationships of a new class of multivalent DNA duplex structures. Specifically, we measured detectable unzipping forces for 155 (of 226) DNA duplexes with rupture forces ranging from over 10 pN to below 3 pN, the pulling force of a single head of myosin II^52^ (**Supplementary Fig. 44**). While our most stable yet sensitive sensor ruptures at 2.7 pN, a fractal multivalent design, among other modifications, could likely lower tension sensing thresholds yet further (**Supplementary Fig. 45, Supplementary Note 7**). These low-tension thresholds should complement DNA-based sensor technologies^28,44,75–77^ that benefit from the tractability and modularity of DNA.

Beyond practical applications, DNA is a tractable biophysical model for understanding force-dependent biomolecular properties. Using DNA as a modular model for multivalency, we found that multivalency was sufficient for decoupling thermodynamic stability from mechanical strength. This architecture was inspired by sensitive mechanosensors in living systems, such as chromatin and Notch^50,78–80^, which also contain interacting domains connected by flexible linkers that drive multivalent interactions. We speculate that multivalency provides sufficient thermodynamic stability to prevent spontaneous activation yet allows responsiveness to small forces. These results challenge the notion that avidity necessarily leads to mechanically robust linkages;^81–83^ instead, our engineering efforts show how mechanosensitivity can arise as an intrinsic nonequilibrium property of multivalency.

In future work, technical modifications could further optimize SM^3^FS for a variety of different applications. Here, we employed a 4x objective and 4 MP camera with a 6.5-µm pixel size, which maximized the number of single molecules that could be measured but limited spatial and temporal resolution to 20 nm and 10 ms, respectively. Imaging with a 100x objective reduced FOV but allowed SM^3^FS to detect 2-nm X-Y displacements of bead position (**Supplementary Fig. 1**), theoretically enabling detection of structural transitions like the 8-nm step of kinesin^84^ or the 25-nm extension of titin Ig domain unfolding.^85^ Coupling SM^3^FS with recently developed ‘macroscope’ configurations^86,87^ capable of ultra-wide FOV imaging at high resolution (34 mm × 34 mm, 3.5-μm pixel size) could further increase sequence and measurement throughput.

The vastness of sequence space requires force spectroscopy technologies that can assay many sequences per experiment. Advances in sequence throughput hold promise to explore the mechanobiome, map sequence-mechanics landscapes, and engineer the dynamic properties of biomolecules far from equilibrium. SM^3^FS expands the scope of single-molecule mechanobiology studies and enables the systematic sequence–mechanics mapping necessary for a new era of single-molecule biophysics.

## Author contributions

M.P.D. developed the SM^3^FS experimental assay. M.P.D. and Y.J.B. measured DNA duplex unzipping forces with the SM^3^FS assay. M.P.D. and Y.J.B developed physical models for tension in microfluidic force microscopy. M.P.D. performed the Monte Carlo simulations of molecule immobilization density. M.P.D., J.O.C., B.F., A.P., and E.P.B. designed the SM^3^FS device. J.O.C., B.F., A.P., and E.P.B. fabricated microfluidic SM^3^FS molds. M.P.D., J.O.C., B.F., A.P., and E.P.B. fabricated SM^3^FS devices. M.P.D. and L.K.P. measured DNA duplex dissociation with the k-MITOMI assay. L.B. designed and fabricated miniMITOMI molds and devices. M.S.B. provided early experimental guidance. M.P.D., A.R.D, and P.M.F. conceived the study. M.P.D. prepared the manuscript, and A.R.D. and P.M.F. supervised the project and assisted in manuscript preparation.

## Acknowledgements

M.P.D acknowledges support from a National Science Foundation Graduate Research Fellowship (DGE-1656518) and a Stanford Bio-X Bowes fellowship. The work supported by funding from an NIH Pioneer Award to P.M.F. (1DP1CA290563), the Stanford Bio-X Interdisciplinary Initiatives Seed Grants Program (IIP R12PFAD), and an NIH R35 grant awarded to A.R.D. (GM130332). B.F., A.P., and E.P.B. were supported by an NSF CAREER Award to P.M.F. (2142336) and a generous gift from the Eleftheria Foundation. We are grateful for support from the Fordyce lab particle party team, help with DNA biochemistry from Caroline Horn, thoughtful comments on manuscript from Shawn Costello, Albert Lee, Daria Passow, and Macy Vollbrecht, discussions about multivalency with Shawn Costello and Peter Suzuki. We are also grateful for helpful discussions at the Single Molecule Biophysics (SMB) Conference 2025, Aspen, CO, particularly with Steve Block, Nynke Dekker, Kier Neuman, Wesley Wong, and Michael Woodside.

## Methods

### Sequences

All DNA oligonucleotides were ordered from IDT, and their sequences are provided in Supplementary Materials.

### Device fabrication and photolithography

Molds for the flow and control layers of the multichannel device were designed in AutoCAD (Autodesk, Inc.) and fabricated as described previously^88^. The height of the flow-layer mold was 15 µm, measured with a profilometer (Tencor).

Molds for the flow and control layers of the miniMITOMI device were designed with the open-source software KLayout (https://www.klayout.de/) and fabricated as previously described, with the exception that the entire flow layer was flow round. All mask designs are available as Supplementary Files in a Zenodo online repository.

For both the multichannel and miniMITOMI devices, “push-down” valved polydimethylsiloxane (PDMS) devices were fabricated from the control-layer and flow-layer molds. To fabricate the control layer, a 1:5 ratio of crosslinker to base of RTV 615 (R.S. Hughes) PDMS was mixed for 3 minutes at 3,000 rpm in an ARM-310 mixer (THINKY). The PDMS mixture was poured over the control-layer mold, degassed under house vacuum for 45 minutes, then baked at 80°C for 40 minutes. Once baked, the control-layer inlets were punched using a 20-gauge needle with a drill press (Schmidt).

To fabricate the flow layer, a 1:20 ratio of crosslinker to base of RTV 615 PDMS was mixed for 3 minutes at 3,000 RPM in the ARM-310 mixer then spin-coated over the flow-layer mold. The PDMS was spun over the molds at 500 RPM for 10 seconds at an acceleration of 133 followed by 1,825 RPM for 75 seconds at an acceleration of 266 (Laurel Technologies Corporation, Model WS-650Mz-23NPPB). The PDMS-coated molds were placed on a flat surface at room temperature (21°C) for 5 minutes to relax the spun PDMS. The flow-layer was baked at 80°C for 40 minutes. Individual control-layers were aligned to the flow layer then baked an additional 50 minutes at 80°C. The bonded two-layer device were cut from the flow-layer mold, and the flow-layer holes were punched. The two-layer multichannel devices were placed on epoxysilane-coated glass slides (Arrayit, SME2) and baked at 90°C for 5 hours.

### Video acquisition and processing

We imaged devices using a Nikon Ti-S or Ti-2 microscope equipped with a CMOS camera (Oxford Instruments, Andor Zyla 4.2), a solid-state light source (Lumencor, SOLA SE Light engine), and an eGFP filter set (Chroma, 49002). Full-field imaging was performed with a 4x objective (Nikon, Plan Apo 4x, NA 0.20), and higher resolution imaging was performed with either a 10x, 20x, or 100x objectives (Nikon). We imaged 1-micron green-fluorescent streptavidin beads (Bangs labs, CDFG004) in the eGFP channel. We imaged 1-micron magnetic streptavidin Dynabeads (ThermoFisher, 65001) and 3-micron polystyrene streptavidin beads (Bangs labs, CP01005) in brightfield, using an overhead light for illumination. All imaging used 1x1 binning and exposure times ranging from 50 to 500 ms. The TrackMate^89^ plugin in FIJI extracted bead traces from the video.

### Generation of a dsDNA library for multiplexed force spectroscopy

A set of 15 biotin- and digoxigenin-modified bifunctional dsDNA duplexes were PCR amplified using Q5 polymerase (NEB, M0492S) from M13mp18 phage DNA template (NEB, N4040S). The primers were labeled with a single 5’ biotin and 5’ digoxigenin modification (IDT). The final set of DNA fragments ranged from 500 to 7,000 bp in increments of 500 bp. Post-amplification gel electrophoresis showed a single strong band of the expected length, with most constructs also showing faint off-target bands (**Supplementary Fig. 6**). Attempted amplification of a 7,500 bp construct yielded multiple bands with a dominant band at 251 bp (**Supplementary Fig. 6**). The amplified DNA was PCR purified (Zymo Research, D4004) and eluted in a 1x borate buffer (LabChem, LC117001) with 1 mM MgCl_2_ (Sigma-Aldrich, M8226), hereafter referred to as borate-magnesium buffer, to a final concentration of 50 ng uL^-1^. Oligo sequences are provided in the supplementary materials.

### General SM^3^FS experiment protocol

Fresh 0.1% Tween-20 (Fisher Bioreagents, BP337-100) was prepared in 1x PBS pH 7.4 (Gibco, 10010023) for each experiment. Polyclonal anti-digoxigenin Fab-AP (Roche, 11093274910) was used to immobilize digoxigenin-modified DNA fragments on the surface of the microfluidic device. All reagents were loaded into devices by loading into Tygon tubing (US Plastic, 56515) and inserted into the PDMS device with a small steel tube (New England Small Tube, 1310-02) unless otherwise noted. All SM^3^FS experiments were performed at 21°C.

A switchable, open-source pneumatic manifold^90^ at 25 PSI opened and closed microfluidic valves by pressurizing the control layer above the flow layer. Control lines were first dead-end filled at 5 PSI with DI water prior to increasing the control-layer pressure source to 25 PSI for closing valves during device operation. Once the control layer was filled, all valves except for outlet valves A-D were closed, and a Fab and 0.1% Tween-20 in PBS solution was plugged into the general inlet (annotated inlets, outlets, and valves in **Supplementary Fig. 2**). Knotted Tygon tubing plugged outlets A-D during surface functionalization steps helped dead-end fill the flow-layer during the Fab patterning step. Alternatively, the Fab solution was introduced through the four outlets, and the backflow, lane-specific, and image valves were pressurized, yielding a more consistent patterning density. An MFCS (Fluigent) or Push Pull (Fluigent) pressure controller pressurized the Fab solution to 750 mbar, the general inlet then backflow valves were sequentially opened, and Fab solution filled up to the image valve. To initiate patterning, the image valve was depressurized, and the Fab solution filled the image region and remainder of the device. Once the Fab solution filled the flow layer (less than one minute) a 0.1% Tween-20 PBS solution replaced the Fab solution in the general inlet, was pressurized to 550 mbar, and washed away unbound Fab. Once Fab was washed away, Fab patterning was complete, and all valves were pressurized, and the Tween-20 solution remained pressurized and plugged into the general inlet.

We loaded individual DNA samples at 1 ng µL^-1^ in borate-magnesium buffer into Tygon tubing, and then plugged them into lane-specific inlets. Each tube was pressurized to 6 PSI, and lane valves were depressurized while keeping the image and backflow valves closed. Trapped air from the lane-specific inlet hole punches were dead-end filled in each channel. Once all bubbles were filled, the image valve was depressurized, and DNA simultaneously flowed from each channel-specific inlet, through the imaging area, to the outlets. After flowing DNA for 30 seconds, the image and lane valves were pressurized to trap DNA solution in each channel. We incubated the DNA in the channels for 15 minutes to bind labeled DNA to the channel surface. After incubation, Tween-20 solution from the general inlet flushed away the unbound DNA.

Streptavidin-coated polystyrene beads were diluted in 0.1% Tween-20 in PBS solution, and then loaded onto the device through the general inlet. Beads were typically bound at 50 mbar of flow pressure. After tethers bound the beads, Tween-20 solution flushed away unbound beads. A pressure function, often a linear pressure ramp or triangle wave cycle, was programmed using Oxygen software that controlled the pressure controller. Ramp experiments typically started with reverse flow (for example, -50 mbar was applied with a Push Pull flow controller), then linearly ramped flow pressure through 0 mbar to the desired setpoint. Ramping through 0 mbar identified single-molecule tethers through measuring the angle and magnitude of bead displacements.

## Multiplexed surface functionalization

### Monte Carlo simulation of varying densities of surface functionalization

DNA molecules were randomly patterned in a 20 x 20 grid. Single versus multiple tether events were analyzed by clustering DNA tether positions and quantifying the number of DNA tethers per bead. DNA anchor-point coordinates were sorted by x-position to facilitate efficient searching for nearest neighbors. For each DNA molecule, nearby tethers within a sway radius defined by the 4-kbp tether length and 1-micron bead diameter. If a neighboring tether was within the sway distance, the bead was attached to both tethers (**Supplementary Fig. 4**). The sway radius of the multi-tether bead was updated to be the intersection of the sway regions of all DNA molecules tethered to the bead. Numbers of beads bound to single versus multiple tethers were quantified as a function of number of randomly placed DNA molecules.

### Surface functionalization and flow-reversal measurement

To pattern a unique surface functionalization in each channel, we followed the general SM^3^FS protocol; however, the surface functionalization and DNA patterning steps were reversed. After the control lines were dead-end filled, the general, backflow, and lane valves were pressurized. We plugged solutions with varying Fab:Tween-20 ratios into the lane-specific inlets, and the surface functionalization solutions were pressurized to 750 mbar. To begin surface patterning, we depressurized the lane valves, and the Fab solutions filled their respective channels. We then plugged in 0.1% Tween-20 solution into the general inlet, depressurized the general inlet valve, and dead-end filled the inlet tree of the device from the general inlet. Once the device was full of solution, the outlet plugs were removed, and Tween-20 solution removed unbound Fab. After washing the device, we introduced 1-micron streptavidin-coated fluorescent green polystyrene beads (Bangs labs, CFDG004) to the device, incubated to allow binding, flushed with Tween-20 solution, then imaged the remaining beads that bound nonspecifically to the immobilized Fab. Next, we added 4-kbp biotin- and digoxigenin-labeled dsDNA at 1 ng µL^-1^ in borate-magnesium buffer to the general inlet, flowed DNA solution across the entire device, and allowed DNA to bind for 30 minutes. Streptavidin beads were reintroduced to the device and bound to the immobilized DNA, unbound beads were removed, and beads were imaged under a triangle pressure wave of -400 mbar to 400 mbar with a wave period of 30 seconds to identify beads that were tethered to single molecules. Beads were imaged with a 500 ms exposure and single second frame rate in the eGFP channel.

## Spatial multiplexing of dsDNA

### Flow-reversal measurement

To pattern a unique DNA duplex in each channel, we first functionalized all channels with 10% Fab v/v with 0.1% Tween20 solution. We next loaded channels with biotin- and digoxigenin-modified dsDNA of increasing lengths by 500-bp increments. We reserved a negative control channel that was only loaded with borate-magnesium buffer, allowing quantification of cross-channel contamination. The flow-reversal measurement followed the same protocol as the multiplexed surface functionalization measurement protocol: we introduced 1-micron streptavidin-coated fluorescent green polystyrene beads (Bangs labs, CFDG004) to the device, incubated to allow binding, flushed with Tween-20 solution, then imaged under a triangle pressure wave of -400 mbar to 400 mbar with a wave period of 30 seconds to identify beads that were tethered to single molecules and measure the length of their linkers. Beads were imaged with a 500 ms exposure and single-second frame rate in the eGFP channel.

### Force-extension measurement of dsDNA

To measure force-extension curves of dsDNA, channels were patterned as in the flow-reversal measurement, except using MyOne streptavidin Dynabeads (Invitrogen, 65001). The MyOne beads were introduced to the device through micromedical LDPE tubing (Scientific Commodities, Inc., BB31695-PE/1) without a small steel tube pin to allow the denser magnetic beads to enter the device. Outlet tubing was not used to reduce channel-to-channel deviations in resistance introduced by different outlet tube heights. Once unbound beads were flushed from the device, the force-extension protocol was imaged in brightfield using a 250 ms exposure and 1 second frame rate. For the first 30 seconds, the flow pressure was set to 0 mbar, and the backflow valve was pressurized to enforce a stringent zero-flow condition to detect the tether attachment point (**Supplementary Fig. 9**). Next, flow pressure was increased in increments of 10 mbar every 30 seconds then by increments of 100 mbar once the flow pressure reached 100 mbar to stretch the dsDNA tether.

### Quantification of dsDNA stretching

For each bead attached to single-molecule tether, bead displacement was estimated as the median bead position for the 30-second pressure interval. The attachment point of the DNA tether was determined as the median bead position at 0 mbar and with a closed backflow valve enforcing zero-flow condition prior to the pressure-step protocol. Single-molecule traces were drift corrected by tracking a stuck particle over the course of the 10-minute measurement. The drift correction was ∼0.5 nm s^-1^ for the first 300 seconds then 0.025 nm s^-1^ afterwards.

### Analysis of dsDNA stretching

The molecular extension, *ΔL*, at each pressure was estimated from bead displacement, *ΔX*, and bead radius, *R*, using a simple geometric model (**Supplementary Fig. 9**):

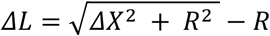

Pressure-extension data from 0 to 100 mbar were fit to a worm-like chain model to get a fit contour and persistence length for each single-molecule tether. Median persistence lengths were used to calibrate tension as a function of flow pressure (**Supplementary Note 1**). The expected influence of molecular geometry on the tension-pressure calibration was calculated from the force balance (Eqn. 1):

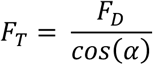

Where the tether-surface angle, *⍺*, was calculated using the geometric model (**Supplementary Note 2**).

### Overextension measurement

To measure overextension of dsDNA, channels were patterned with a 5% Fab solution. Larger, 3-micron streptavidin-coated polystyrene beads (Bangs labs, CP01005) were introduced to attach to DNA tethers, and a linear pressure ramp was applied from -50 to 1,000 mbar in 90 seconds, corresponding to a drag force ramp rate of ∼2.2 pN s^-1^ and a drag force/pressure calibration of ∼190 pN bar^-1^. Beads were imaged in brightfield with a 250 ms exposure and 0.5 second frame rate. Outlet tubing was not used to reduce channel-to-channel deviations in resistance introduced by different outlet tube heights.

### Quantification of dsDNA overstretching

The displacement and tension force of overextension were extracted from single-molecule traces by identifying the range of frames that result in a step-jump in bead position corresponding to the B-to-S transition. A moving window two-frame average was applied to all traces to enhance X-Y spatial resolution, specifically to help unambiguously detect the ∼120 nm expected overextension for the 500-bp tether (**Supplementary Fig. 16**). Drag force was estimated from the overextension transition of the 7-kbp tether, which corresponds to a tension of 65 pN; the corresponding drag force was 92% of the tension, ∼60 pN, as estimated by the geometric model. The drag force of overextension was determined as the point at which the bead displacement passed a 5% threshold of the expected displacement for long tethers beyond their already fully extended state. A more sensitive threshold of 20% identified genuine transitions for tethers shorter than 2,500 bp. The overextension displacement was calculated as the change in x-position of the bead 5 seconds before to 5 seconds after overextension. The drag force and displacement of overextension were fit to a normal distribution to identify the most probable overextension force, displacement, and associated errors.

### Analysis of dsDNA overstretching

The expected displacement from overextension was calculated from the geometric model (**Supplementary Note 2**) in which the end-top-end distance of the tether extended an additional 70% at 65 pN from a fully extended tether of contour length, L_0_:

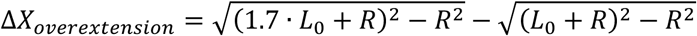

The expected drag force of overextension was calculated from the force balance (Eqn. 1):

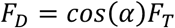

In which the tension force at overextension was set to a constant 65 pN across tether lengths, and the molecular geometry, *⍺*, was calculated using the geometric model (**Supplementary Note 2**).

### Parallelized unzipping measurements of DNA duplexes

To immobilize variable DNA oligonucleotides in channels, oligonucleotides were annealed to dsDNA handles then introduced to the devices in the same spatially multiplexed approach as before. All measured DNA oligonucleotides (IDT) were ordered at 100 µM in 1x IDTE buffer and not purified. Specific care was taken to ensure that mainly full-length duplexes were measured on device. DsDNA handles of two lengths, 3- and 7-kbp, were amplified via PCR with PrimeStar GXL polymerase (Takara, R050B) using a digoxigenin-modified primer and a primer containing an internal abasic site to leave a 5’ overhang for attaching variable DNA oligonucleotides.

Handles were PCR purified and eluted in borate-magnesium buffer. An excess of variable or fiducial oligonucleotides at a final concentration of 1 µM were incubated with the 3- and 7-kbp handles at 1 ng µL^-1^ (∼0.2 nM final concentration) in borate-magnesium buffer at 50°C for 10 minutes. Partially synthesized oligonucleotides likely lacked a complete 5’ generic, complimentary sequence that was required to hybridize to the dsDNA handle. The surface functionalization protocol for the unzipping experiments followed the standard protocol with a 15% v/v solution of Fab solution for patterning channels. After binding the DNA, unbound DNA was washed away with 0.1% Tween-20 solution, leaving primarily full-length immobilized oligonucleotides in the device.

To immobilize the partner oligonucleotide on beads, an excess of variable and fiducial oligonucleotides (0.5 µL of 100 µM oligonucleotide) was incubated with 1 µL of 10 µM generic biotinylated oligonucleotide in 5 µL of PBS at 50°C for 10 minutes. 1 µL of hybridized DNA mixture was then incubated with 2 µL of MyOne Dynabeads (Invitrogen, 65001) in 5 µL of PBS and rotated at room temperature for 15 minutes to allow the beads to bind the biotinylated DNA. DNA immobilization density on bead was selected to minimize nonspecific binding (data not shown). After binding and prior to flowing onto the device, the beads were magnetically pulled down, washed five times with 100 µL of Tween-20 solution to remove unbound variable or fiducial DNA, then suspended in 50 µL Tween-20 solution. Beads were sonicated (VWR, 97043-964) for 30 seconds at 21°C, then 5 µL of fiducial beads were mixed with 15 µL of variable beads prior to loading onto the device. The generic, complimentary sequence on the oligonucleotides were located on the 3’ end to generate a DNA duplex that was pulled in an unzipping geometry. As a result, beads could contain partially synthesized oligonucleotides that were truncated on the 5’ end. The longest oligonucleotide measured was 60 nt, and assuming a per-nucleotide synthesis yield of 99.4%, approximately 70% of oligonucleotides on the bead were expected to be full-length.

Once DNA was immobilized on channels and beads, oligonucleotide-displaying beads were introduced to the imaging area at 25 mbar (corresponding to ∼0.75 pN) to generate hybridized duplex-tethered beads. The oligonucleotide beads were loaded in LDPE tubing (Scientific Commodities, Inc., BB31695-PE/1) without a small steel tube pin. Next, Tween-20 solution gently flushed out unbound beads at 25 mbar for 5 minutes so as to not rupture hybridized duplexes. Since truncated oligonucleotides are less likely to sustain a stable interaction with the channel-immobilized oligonucleotide, primarily full-length DNA duplex interactions remained after washing.

To measure the mechanical strength of the DNA duplexes, the flow pressure of Tween-20 solution was ramped from -50 to 1000 mbar in 60 seconds, corresponding to a tension ramp rate of around 0.5 pN s^-1^. After unzipping the DNA duplex, the bead-associated oligonucleotide and bead were washed away, leaving an unhybridized channel oligonucleotide. A new set of beads displaying a new oligonucleotide sequence could then be introduced to channels.

The 100% GC content duplexes required prehybridizing the DNA duplexes due to the formation of stable secondary structures that prevented hybridization in the device. To prehybridize variable and fiducial DNA duplexes, the dsDNA tether (1 ng µL^-1^, final concentration), “channel” oligonucleotide (5 µM), partner “bead” oligonucleotide (10 µM), and the generic biotinylated oligonucleotide (15 µM) were mixed in borate-magnesium buffer and incubated at 65°C for 10 minutes, 60°C for 15 minutes, and 50°C for 20 minutes. After annealing, 10 µL of fiducial DNA solution was added to each variable duplex solution. The hybridized DNA was then added into individual channels on the SM^3^FS device following the standard protocol. Once the DNA duplexes were immobilized, MyOne streptavidin beads in 0.1% Tween-20 solution were introduced to the device and incubated with the immobilized duplexes. Because unbound oligonucleotides formed stable secondary structures, the required prehybridization step likely biased unzipping measurements toward measuring kinetically stable duplexes that remained bound for the duration of the experiment (**Supplementary Fig. 40**).

## Analysis of DNA duplex unzipping

### Quantification of DNA unzipping

The displacement distance and angle of the bead upon the change of direction of flow indicated the beads attached to single-molecule tethers and the identity of the hybridized duplex (variable or fiducial) based on the distance of the displacement. An observation threshold of 20 single-molecule ruptures was set for reporting an unzipping force (**Supplementary Fig. 34**). The most probable pressure at which beads detached was determined from a Gaussian fit. The rupture pressure of the fiducial duplex indicated the pressure at which the tension on the 7-kbp tether was 9.0 or 9.65 pN, depending on the identity of the fiducial. A 15-bp duplex with 47% GC content was used as the fiducial for calibrating most duplex unzipping measurements, and a 39-bp duplex with 0% GC content was used for a subset of measurements (indicated in **Supplementary Data**) to minimize cross-talk with variable sequences. To apply the fiducial tension calibration to the variable duplex measurement, a tension conversion factor of 106% (0.94^-1^) accounted for the amplification of drag force on the shorter 3-kbp tether that anchored the variable duplex. The median strength of the duplex attachment linkages was >30 pN; therefore, rupture events below 30 pN were likely unzipping of the duplex (**Supplementary Fig. 29**).

### Analysis of standard DNA duplex unzipping (Library 1)

We fit the most probable rupture force, *F*^∗^, to a simple empirical model relative to the duplex size, *L*:

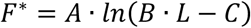

The logarithmic model captured important features of the unzipping force trends across standard DNA duplexes: First, the model followed the general observation that the rupture force increased logarithmically for short DNA duplexes. The log-linear relationship may reflect that the free energy barrier of unzipping increases linearly with the length of a standard duplex. Second, for some minimum length of duplex of length (1 + *C*)*B*^−1^, the duplex was insufficiently stable to yield a measurable rupture force. Below the critical length, the predicted rupture force was less than 0 pN and not physically possible, indicating that the near-zero force thermal dissociation rate was fast relative to the experimental time scale on the order of minutes. Finally, the prefactor *A* scaled the logarithmic function to match the units of force. The fit prefactor was not a strong function of GC-content, ranging from 1.53 to 1.58 pN. The error in extrapolated unzipping forces was propagated from the covariance matrix of the nonlinear fit. While the model captures the relevant physics for unzipping short duplexes, the logarithmic model is unbounded and thus does not generalize to arbitrarily long duplexes.

A well-established physical model for DNA rupture mechanics:^91^

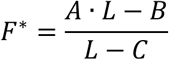

is mathematically bounded for arbitrarily long duplexes and fit the rupture force data, but it did not extrapolate as accurately to the rupture forces of long (100 bp) duplexes compared to the logarithmic model. We expect that including unzipping data across a broader range of duplex lengths, in particular for the 100% GC-content duplexes, could improve the accuracy of prediction for the physically grounded model.

### Bell-Evans model fits from rupture force distributions

Cumulative rupture force distributions were fit to the Bell-Evans model.^24,68,92^ Many pairs of Bell-Evans model parameters are consistent with a given rupture force and ramp rate (**Supplementary Fig. 42**). Therefore, to estimate the distance to the transition state, the zero-force dissociation rate was experimentally determined and used to estimate the distance to the transition state (**Supplementary Note 5**).

### Zero-force dissociation kinetics measurements of DNA duplexes

MiniMITOMI devices were patterned with a biotinylated BSA and neutravidin and blocked with BSA as described in previous MITOMI assays. DNA duplexes were assembled off chip with a fluorescein-modified oligonucleotide that bound the same complimentary sequence as the dsDNA tether and the same biotinylated generic oligo used to immobilize DNA to beads. Biotin oligonucleotide, oligonucleotide A, interacting oligonucleotide B, and fluorescein oligonucleotide mixed in PBS at final concentrations of 5, 10, 12.5 and 15 µM, respectively, then annealed at 50°C for 20 minutes to ensure that a fully assembled DNA duplex that was immobilized and fluorescently labeled. Labeled duplexes were introduced sequentially into the miniMITOMI device. While introducing DNA, pneumatic “button” valves were pressurized to prevent cross-contamination between channels. Once the DNA duplexes were introduced to their assigned lanes, the pneumatic button valves were opened to expose the neutravidin surface to the biotin-labeled duplexes. DNA duplexes were incubated with the neutravidin-coated surface for 15 minutes to allow binding. After binding, the button valves were closed to trap the bound DNA duplex, and PBS washed away unbound DNA. The valves were opened and the array of fluorescent duplexes was imaged in the GFP channel every minute with an exposure time of one second to measure duplex dissociation as the decay of the fluorescein signal. PBS flow pressure was 250 mbar to remove dissociated DNA. Measurements were made at 21°C.

### Quantification of dissociation kinetics

Fluorescence intensity was quantified as the average pixel intensity of the immobilized spot. The background intensity was calculated as the average pixel intensity of the surrounding reaction chamber. The average of three background-subtracted chambers was fit to a single-exponential decay to calculate a bond lifetime, and error was estimated from the covariance matrix of the nonlinear fit.

### Photobleaching correction

The rate of photobleaching, in addition to other sources of sequence-independent loss of fluorescent signal, was estimated as the dissociation rate of the 32-bp duplex under constant flow. The reported bond lifetimes, *τ*, are corrected for photobleaching:

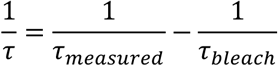

Where *τ*_*bleach*_ = *τ*_32–*bp* *duplex*_. The error of the corrected bond lifetime was propagated from error in the measured dissociation bond lifetime and effective time-constant for photobleaching.

