## Supplementary Material for "Engineering the mechanosensitivity of single DNA molecules via high-throughput microfluidic force spectroscopy"

**Overview:**

Supplementary Note 1: Calibrating flow pressure to drag force via stretching dsDNA
Supplementary Note 2: Physical models of tension
Supplementary Note 3: Properties of bead-size distributions explain non-normally distributed B-to-S transition forces
Supplementary Note 4: The median rupture force of DNA duplexes with fast zero-force off rate and a non-negligible distance to the transition state
Supplementary Note 5. Constrained estimation of the distance to transition state
Supplementary Note 6: Transition state models of the multivalent duplex
Supplementary Note 7: Design principles for lower tension sensing thresholds of DNA
Supplementary Figures 1 – 45
Supplementary Table 1: Parameters of bead-based force spectroscopy assays

**Supplementary Note 1. Calibrating flow pressure to drag force via stretching dsDNA.**

The worm-like chain (WLC) model[1] relating expected DNA length to applied tension is described by the following equation:

$$\frac{F_T P}{k_B T} = \frac{1}{4}\left(1 - \frac{l}{L_0}\right)^{-2} - \frac{1}{4} + \frac{l}{L_0} \qquad Eqn. 1$$

Here, $F_T$ is tension on the tether, $P$ is the persistence length of the tether, $k_B$ is the Boltzmann constant, $T$ is temperature, $l$ is the molecular extension of the tether, and $L_0$ is the contour length of the tether. In $SM^3FS$, we apply flow pressure, $P_{bar}$, and measure the displacement of the bead, ΔX, which maps to a molecular extension, $l$, for a given bead radius. Since we know the relationship between tension and drag forces from Eqn. 1 (from main text),

$$F_T = \frac{F_D}{cos(\alpha)} \qquad Eqn. 2$$

and the drag force increases linearly with flow pressure, $F_D = m{\cdot}P_{bar}$, we may rewrite Eqn. 1 as:

$$F_T = \frac{m \cdot P_{bar}}{cos(\alpha)} \qquad Eqn. 3$$

Here, $m{\cdot}cos(\alpha)^{-1}$ is a linear constant that includes molecular geometry to relate pressure to tension force. Plugging this expression for $F_T$ into Eqn. 1:

$$\frac{P_{bar} \cdot m \cdot P}{cos(\alpha)\, k_B T} = \frac{1}{4}\left(1 - \frac{l}{L_0}\right)^{-2} - \frac{1}{4} + \frac{l}{L_0} \qquad Eqn. 4$$

For a given pressure-extension curve, a nonlinear regression of Eqn. 4 returns a fit contour length, $L_0$, which is the true contour length. The regression also returns a fit persistence length, $P_{fit}$, which we define as a function of the true persistence length, the drag-force/pressure slope, and the molecular geometry:

$$P_{fit} = \frac{m \cdot P}{cos(\alpha)} \qquad Eqn. 5$$

In the case where the molecular tether is very long, α approaches 0; therefore, the pressure-to-drag force calibration curve slope, $m$ with units pN $mbar^{-1}$, is simply the ratio of the fit persistence length and true persistence length:

$$m = \frac{P_{fit}}{P} \qquad Eqn. 6$$

For intermediate tether lengths, $\alpha \neq 0$. Therefore, the ratio of the fit and true persistence length includes the $cos(\alpha)^{-1}$ term:

$$\frac{P_{fit}}{P} = \frac{m}{cos(\alpha)} \qquad Eqn. 7$$

The ratio $m \cdot cos(\alpha)^{-1}$ is the tension-to-pressure calibration curve slope, as derived above. Importantly, the persistence length, *P*, for this form of the WLC model varies as a function of the contour length[2,3]; the empirical relationship between contour length and persistence length is:

$$P(L_0) = P_\infty \left(1 + \frac{aP_\infty}{L_0}\right)^{-1} \qquad Eqn. 8$$

Where $P_\infty$ is the persistence length of an infinitely long DNA tether, $P_\infty$ = 48 nm, and *a* is an empirically fit parameter of 2.75.[2,3]

To assess the effect of molecular geometry (i.e. size of tether versus bead) on the relationship between tension and drag force, a ratio of ratios of the fit and true persistence lengths between a tether of length *x*, and a tether of constant length *C*, cancels out the calibration factor *m*, leaving a ratio of $cos(\alpha)$ terms:

$$\frac{\frac{P_{fit}}{P}\left(L_{0,x}\right)}{\frac{P_{fit}}{P}\left(L_{0,C}\right)} = \frac{cos(\alpha_C)}{cos(\alpha_x)} \qquad Eqn. 9$$

And if we make *C* infinitely long ($C = \infty$) compared to the bead radius, $\alpha_\infty$ approaches zero, and $cos(\alpha_\infty)$ approaches 1:

$$\frac{\frac{P_{fit}}{P}\left(L_{0,x}\right)}{\frac{P_{fit}}{P}\left(L_{0,\infty}\right)} = cos(\alpha_x)^{-1} \qquad Eqn. 10$$

This allows assessment of the impact of molecular geometry on applied tension (**Supplementary note 2, Supplementary Fig. 21**).

**Supplementary Note 2. Physical models for tension.**

We first calculate the Reynolds number at the highest flow pressure on the multichannel device:

$$Re = \frac{\rho v \mathcal{L}}{\mu} \qquad Eqn. 11$$

Here, the fluid density $\rho$ = 1,000 kg m$^{-3}$; the flow velocity $v_{max}$ < 10 mm s$^{-1}$; the characteristic length (here, the hydraulic diameter of a rectangular pipe) $\mathcal{L}$ = 30 µm; and the fluid viscosity $\mu$ = 0.001 Pa s. At the maximum flow rate, Re < 0.3, indicating that flow is laminar. Therefore, the flow velocity, *v*, of our system is proportional to the applied pressure:

$$v(x, y, z, P_{bar}) = k(x, y, z) \cdot P_{bar} \qquad Eqn. 12$$

For Re < 1, the hydrodynamic force exerted on any spherical particle is well approximated by Stokes' law:[4]

$$F_D = 6\pi\mu R v \qquad Eqn. 13$$

Here, $R$ is the radius of the sphere and *v* is the undisturbed velocity of the pipe flow. Combining equation 12 and 13, the hydrodynamic force exerted on a particle at a certain position is proportional to the applied pressure

$$F_D \propto P_{bar} \qquad Eqn. 14$$

Since the bead radius is much smaller than the flow channel dimensions, the beads experience shear flow (**Fig. 1B**). We approximate the flow velocity at the height of the bead radius, R, above the surface in shear flow at a shear rate, $\gamma$:

$$v = \gamma(P_{bar}, \mu, \mathcal{L}, \ldots) \cdot R \qquad Eqn. 15$$

Therefore, we can write the hydrodynamic drag force as:

$$F_D = 6\pi\mu R v = 6\pi\mu\gamma R^2 \qquad Eqn. 16$$

Additional correction factors can be applied for near-wall drag and torque, but calibration using a well-characterized force fiducial (such as the B-to-S transition of DNA) accounts for this correction factor. Here, we assess three models to explain SM$^3$FS: (1) a no-torque model, (2) a model that incorporates torque, and (3) a model that incorporates both torque and friction.

We define a force factor, which is the ratio of tension force (exerted on the molecule) to hydrodynamic drag force and is a function of the molecular geometry defined by the tether length, *L*, and the bead radius, *R*:

$$f_f(L, R) = \frac{F_T(L, R)}{F_D(R)} \qquad Eqn. 17$$

From the force balance, the friction factor is a function of the molecular geometry:

$$f_f(L,R) = \frac{1}{cos\big(\alpha(L,R)\big)} \qquad Eqn.\,18$$

To assess physical models for tension in microfluidic force spectroscopy, we compare a relative force factor of various tether lengths $L$ to the force factor for $L_0$ = 7,000 bp. For example, we measure the pressures that cause the B-to-S transition for these tethers, which correspond to a 65 pN of tension force, and since the pressure is proportional to the velocity and thus the drag force, the ratio of pressures required for B-to-S transition reflects the ratio of force factors:

$$\frac{f_f(L)}{f_f(L_0)} = \frac{\dfrac{65pN}{F_{D,B-to-S}(L)}}{\dfrac{65pN}{F_{D,B-to-S}(L_0)}} = \frac{F_D(L_0)}{F_D(L)} = \frac{P_{bar,L_0}}{P_{bar,L}} \qquad Eqn.\,19$$

Thus, we derive the relative force factor purely from experimental data, specifically the pressure of B-to-S transition for different tethering lengths. By comparing the relative force factor in the experiment, and the relative force factor produced by the model, we can assess the accuracy of our physical models for tension.

**No-torque model (The primary geometric model)**

We first consider the force balance without torque. In this model, the hydrodynamic force on the bead is balanced by the tension force of the tether and the normal force of the surface (**Supplementary Fig. 20**). Here, the drag force is the x-component of the tension force:

$$F_D = F_{T,x} = F_T \cos\alpha \qquad Eqn.\,20$$

Since the model neglects any existing torque, the x-component of the tension force $(\cos\alpha)$ is geometrically calculated assuming that the tension vector passes through the center of the bead:

$$\cos\alpha = \frac{\sqrt{(L+R)^2 - R^2}}{L+R} \qquad Eqn.\,21$$

Rearranging, a dimensionless geometric variable, $K = \frac{L}{R}$, reflects the relative size of the tether to the bead. The tension force is a function of the drag force and molecular geometry:

$$F_T = \frac{6\pi\mu\gamma R^2}{cos\alpha} \qquad Eqn.\,22$$

As a result, the ratio of two tension forces applied to tethers of length L versus $L_0$ is simply a ratio of the molecular geometry terms:

$$\frac{F_T}{F_{T0}} = \frac{cos\alpha_0}{cos\alpha} \qquad Eqn. 23$$

This model aligns well with the measured force factors (**Supplementary Fig. 20**), but it simplifies the free body diagram by excluding torque. Hydrodynamic torque is inherently included in the Poiseuille flow: the fluid velocities at the top and the bottom of the bead are different, creating a tendency to spin. Therefore, we need to include torque within the model and examine how torque would alter the force factor.

**Torque model**

In this model, we include hydrodynamic torque in the system:

$$T = 4\pi\mu\gamma R^3 \qquad Eqn. 24$$

The hydrodynamic torque is balanced by the tension of the tether, which, in this time, would not go over the sphere center to create a reverse torque (**Supplementary Figs. 22** and **23**). The rotated tether creates two variables: $\theta$, which is the angle between the sphere radius and the tether, and $\alpha$, which is the angle between the tether and surface. The torque model has two unknowns, $\theta$ and $\alpha$. Additional equations are required to specify these two variables. One is from torque balance, and the others are from the molecular geometry induced by torque.

The force balance still holds:

$$\begin{cases} F_D = F_T cos\alpha \\ F_D = 6\pi\mu\gamma R^2 \end{cases} \qquad Eqns. 25\ and\ 26$$

Therefore, the ratio of tensions applied to tethers of length L versus $L_0$ remains the same as the no-torque model:

$$\frac{F_T}{F_{T0}} = \frac{cos\alpha_0}{cos\alpha} \qquad Eqn. 27$$

However, torque now rotates the bead and increases the tether angle $\alpha$. To determine the extent of rotation and update $\alpha$, the torque balance yields the following relationship between $\theta$ and $\alpha$:

$$\begin{cases} T_D = F_T R sin\theta \\ T_D = 4\pi\mu\gamma R^3 \end{cases} \qquad Eqns. 28\ and\ 29$$

Solving yields:

$$\frac{sin\theta}{cos\alpha} = \frac{2}{3} \qquad Eqn. 30$$

Through the geometry in the system (**Supplementary Figs. 22A,B**), which includes $\theta$ and $\alpha$, bead radius $R$, the distance between the bead center and the surface attachment point of the tether $C$, and the tether length $L$, we get two additional equations that, along with the above Eqn. 30, fully specify the molecular geometry in the presence of torque.

From triangle LRC (**Supplementary Fig. 22C**):

$$L^2 + R^2 - 2LRcos\theta = C^2 \qquad Eqn. 31$$

From triangle LDX (**Supplementary Fig. 22D**):

$$L^2 + X^2 - 2LXcos\alpha = D^2 \qquad Eqn. 32$$

Several geometric steps transform Eqn. 32 to the following:

$$L^2 + C^2 - R^2 - 2L\sqrt{C^2 - R^2}cos\alpha = 2R^2[1 - sin(\theta + \alpha)] \qquad Eqn. 33$$

Together, the torque balance along with the geometric constraints yield a system of equations that can be numerically solved to yield C, $\theta$ and $\alpha$ for a given L and R:

$$\begin{cases} \frac{sin\theta}{cos\alpha} = \frac{2}{3} \\ L^2 + R^2 - 2LRcos\theta = C^2 \\ L^2 + C^2 - R^2 - 2L\sqrt{C^2 - R^2}cos\alpha = 2R^2[1 - sin(\theta + \alpha)] \end{cases} \qquad Eqns. 34 - 36$$

Notably, the same dimensionless variable $K = \frac{L}{R}$ from the no-torque model arises from the above system of equations 34 to 36. Therefore, like the no-torque model, the molecular geometry is parameterized by $K$. We calculated the bead horizontal movement, *X*, for the with- and without-torque models for 1- and 3-micron beads attached to tethers ranging from 500 to 7,000 bp (**Supplementary Fig. 24**). As expected, torque rotated the bead and increased $\alpha$, but the associated horizontal movements were similar between models, around a 5% relative difference for the tested geometries, but the expected force factors differed between models (**Supplementary Fig. 25**).

Although including torque makes physical sense, the relative tensions predicted by the with-torque model do not match the experimental results (**Supplementary Fig. 21**), suggesting that other factors may influence the molecular geometry and the relationship between hydrodynamic force and tension.

**Friction model**

For the tether to resist horizontal drag force, the tether must also vertically exert force as determined by the molecular geometry (**Supplementary Fig. 20**). Normal force from the channel surface opposes the vertical component of the tension force, and the channel exerts static friction on the bead proportionally to the normal force. As torque rotates the bead, the angle $\alpha$ increases, as does the normal force and associated friction force.

Depending on the magnitude of the hydrodynamic torque, frictional force may or may not be sufficient to balance it. We first examine the scenario where friction can fully counteract the torque, showing that the system reduces to the no-torque model. We also briefly describe the alternative case, where friction fails to balance the torque, leading to behavior analogous to the with-torque model.

Friction acts at the point of contact between the bead and surface, generating a counter-torque that opposes the torque induced by fluid shear. When friction is sufficient to balance the torque, the torque balance condition is:

$$\begin{cases} T_F = F_f R \\ T_D = 4\pi\mu\gamma R^3 \end{cases} \qquad Eqns. 37\ and\ 38$$

Setting the torque from friction, $T_F$, equal to the hydrodynamic torque yields a frictional force, $F_f$, that is sufficient to balance hydrodynamic torque:

$$F_f = 4\pi\mu\gamma R^2 \qquad Eqn. 39$$

In this case, the tether does not generate torque and remains straight, pointing toward the bead's center, like the no-torque model. The horizontal component of the tension force balances both the hydrodynamic drag and friction:

$$\begin{cases} F_D + F_f = F_T \cos\alpha \\ F_D = 6\pi\mu\gamma R^2 \\ F_f = 4\pi\mu\gamma R^2 \end{cases} \qquad Eqns. 40 - 42$$

This gives the total tension force:

$$F_T = \frac{10\pi\mu\gamma R^2}{\cos\alpha} \qquad Eqn. 42$$

Again, $\cos\alpha$ depends on the geometry between the tether and bead:

$$\cos\alpha = \frac{\sqrt{(L+R)^2 - R^2}}{L+R} \qquad Eqn. 41$$

Compared to the no-torque model, the only change is in the constant coefficient ($10\pi\mu$ vs. $6\pi\mu$) Therefore, incorporating friction theoretically increases the tether force by a factor of $\frac{5}{3}$, but maintains the same geometric dependence and linearity.

If the friction force itself could not hold against hydrodynamic torque, an additional torque from the tether would resist hydrodynamic torque. In this case, similar $\theta$ and $\alpha$ variables specify the molecular geometry. The friction force is proportional to the normal force:

$$\begin{cases} F_f = \eta F_N \\ F_N = F_T \sin\alpha \end{cases} \qquad Eqns. 42\ and\ 43$$

Conducting a similar force balance of the horizontal force:

$$\begin{cases} F_T \cos\alpha = F_D + F_f \\ F_D = 6\pi\mu\gamma R^2 \\ F_f = \eta F_T \sin\alpha \end{cases} \qquad Eqns. 44 - 46$$

As $\eta$ approaches zero, the torque balance model is recovered. However, if the friction coefficient is not negligible, we may again set the torque from friction, $T_F$, equal to the hydrodynamic torque and calculate $\eta$ that is necessary to balance the hydrodynamic torque:

$$\eta = \frac{4}{10 \cdot \tan(\alpha)} \qquad Eqn. 47$$

Prior measurements of Tween-20 absorbed on hydrophobic surfaces yielded a large static friction coefficient ~1.[5] Using this friction coefficient, the geometric threshold at which friction force is meaningful is $\tan^{-1}(0.4) = 22°$. Notably, this threshold is independent of bead size but depends on the geometric dimensionless variable Κ. For a 1-micron bead, 22° corresponds to tethers shorter than 2,500 bp having sufficient friction force to balance hydrodynamic torque. For a 3-micron bead, tethers shorter than 7,500 bp have sufficient friction force to balance hydrodynamic torque, consistent with our measurements (**Supplementary Fig. 21**)

In the context of the force balance, for angles lower than 22°, torque can alter the geometry; however, as the length of the tether increases, the drag force approaches the tension force. Thus, correcting for hydrodynamic torque is not necessary, and its impact on the free body diagram becomes vanishingly small. Notably, in this regime of geometries below 22°, the linear pressure-to-force relationship holds (**Supplementary note 2: with-torque model**); a correction can be made relating the bead centroid to molecular extension, but the correction is minor (**Supplementary Fig. 24**).

**Supplementary Note 3. Properties of bead-size distributions explain non-normally distributed B-to-S transition forces.**

A small subset of single-molecule tethers overstretched at higher forces than expected (**Fig. 4C**); these beads likely had smaller diameters, suggesting an overestimation of applied drag force when using median bead parameters. The coefficient of variation of the overextension drag force decreased with tether length following $cos(\alpha)^{-1}$ and asymptotically approached a CV of 7.3%, roughly twice the CV of the bead diameter (CV = 4%, **Supplementary Fig. 19**), as expected for drag force on a bead in shear flow.

**Supplementary Note 4. The median rupture force of DNA duplexes with a fast zero-force off rate and a non-negligible distance to the transition state**

For a molecular interaction that is well approximated by the Bell model[6] (Eqn. 2, from main text) being pulled apart at a constant force ramp rate ($\dot{F}$),[7] in which δ is the distance to the transition state and $k_0$ is the dissociation rate at zero force,[8] the median rupture force ($F_{50\%}$), at which the CDF($F_{50\%}$) = 0.5:

$$CDF(F_{50\%}) = 0.5 = 1 - e^{\left(-\frac{k_0 k_B T}{\dot{F}\delta}\left[e^{\frac{F_{50\%}\delta}{k_B T}} - 1\right]\right)} \qquad Eqn. 48$$

Solving for $F_{50\%}$ yields the median rupture force:

$$F_{50\%} = \frac{k_B T}{\delta} ln\left(\frac{\dot{F}\delta ln(2)}{k_0 k_B T} + 1\right) \qquad Eqn. 49$$

For fast off rates ($k_0 \gg \frac{\dot{F}\delta ln(2)}{k_B T}$) and non-zero distances to the transition state of unzipping, $F_{50\%}$ ~ ln(1) = 0 pN. Plugging Eqn. 49 into the Bell model (Eqn. 2, from main text) yields the dissociation rate at the median rupture force:

$$k(F_{50\%}) = \frac{\dot{F}\delta ln(2)}{k_B T} + k_0 \qquad Eqn. 50$$

Which simplifies to $k(F_{50\%}) \sim k_0$ for fast dissociation rates.

**Supplementary Note 5. Constrained estimation of the distance to transition state**

The most probable rupture force ($F^*$) at a constant force ramp rate ($\dot{F}$) for a molecular interaction that is well approximated by the Bell-Evans model,[6,7] in which $\delta$ is the distance to the transition state and $k_0$ is the dissociation rate at zero force, is:[8]

$$F^* = \frac{k_B T}{\delta} ln\left(\frac{\dot{F}\delta}{k_0 k_B T}\right) \qquad Eqn. 51$$

Thus, the dissociation rate at the most probable rupture force is:

$$k(F^*) = \frac{\dot{F}\delta}{k_B T} \qquad Eqn. 52$$

Plugging into the Bell model yields the following equation:

$$\frac{\dot{F}\delta}{k_B T} = k_0 \; exp\left(\frac{F^*\delta}{k_B T}\right) \qquad Eqn. 53$$

Numerically solving yielded $\delta$ as a function of $k_0$, $F^*$, and $\dot{F}$, providing an estimate of $\delta$ from experimentally derived values for $k_0$ and $F^*$(**Supplementary Fig. 42**). We estimated $k_0$ from the k-MITOMI solution-phase dissociation measurement. Although the solution-phase measurement is likely an overestimate of the true zero-force dissociation rate of the unzipping pathway, the estimate was based on an experimental measurement that was closely related to the unzipping pathway. The measured fluorescence signal decay rates for the 12, 14, and 32 bp standard duplexes were equal and assumed to be the photobleaching rate. Therefore, the dissociation rate of these duplexes was likely slower than the photobleaching rate, and the corresponding distance to the transition state was an underestimate. The parameters $F^*$ and $\dot{F}$ were measured and set from the SM$^3$FS assay.

**Supplementary Note 6. Transition state models of the multivalent duplex**

Given internal mismatches can generate jagged free energy landscapes for duplexes under unzipping force,[9,10] we considered two models for multivalent duplex unzipping. The first model represents a completely unzipping DNA duplex, in which nearly all of the unit duplexes are unzipped in the TS ($\delta_1$), like a standard DNA duplex (**Supplementary Fig. 43**). Alternatively, we considered a second model in which the transition state to unzipping represents unzipping of a single unit duplex ($\delta_2$, **Supplementary Fig. 43**). In the second model, once enough force is applied to unzip the first unit duplex, the remaining duplexes unzip irreversibly. We hypothesized that the nature of the transition state is a function of linker length: once linkers are long enough, the unit duplexes of the multivalent duplex would become mechanically decoupled.

Given such low unzipping forces (< 3 pN), the elasticity of ssDNA may influence the number of predicted unzipped duplexes that contribute to the transition state. The average extension of an ideal elastic oligonucleotide at 2.5 pN is approximately 50% of fully extended ssDNA, yielding an approximate contour length of 0.25 nm per nucleotide, assuming that there is minimal secondary structure formation in the unbound state as predicted by UNAfold at room temperature and 137 mM NaCl.[11,12] The distance to the transition state of the 8 × (4-bp + 4T) construct was calculated to be 12.6 nm, corresponding to the unzipping of approximately three to four unit duplexes. The limiting cases for transition state models 1 and 2 were approximately 30 and 2 nm, respectively. A distance to the transition state of 12.6 nm is consistent with a model in which several, but not all, of the unit duplexes are unzipped in the transition state (**Supplementary Fig. 43**). Once three-to-four unit duplexes are unzipped, the remaining duplexes unzip irreversibly. Assuming symmetry in equilibrium free energy landscape of hybridization, since the unzipping force is sufficient to unzip the remaining unit duplexes, the bound state at the critical rupture force is likely partially unzipped.

**Supplementary Note 7. Design principles for lower tension sensing thresholds of DNA**

We propose three approaches to extend the tension sensing threshold below 3 pN. First, we observed that increasing the size of the linkers decreased the mechanical stability. Further increasing linker length would lower mechanical stability to the $\delta_2$ regime, beyond which the unzipping force is expected to remain constant. Second, reducing the size of the unit duplex to the fundamental limit of a single base pair could lower the tension-sensing threshold. Such a sensor could report on the mechanical strength of a single-base pairing interaction; however, even if this construct could stably hybridize at zero-force, the strength of the single-base pair interaction would impose a limit on tension sensing. Third, a fractal design could extend beyond the tension-sensing limits below single-base pair unit repeat. For the fractal design, the unit duplex itself would be a multivalent duplex (**Supplementary Fig. 45**). The unit multivalent duplex allows finely balancing the destabilization from the flexible linkers with stabilization of the hybridized regions, which in principle, would yield a sensor that ruptures at arbitrarily low tensions.

### Supplementary Figures

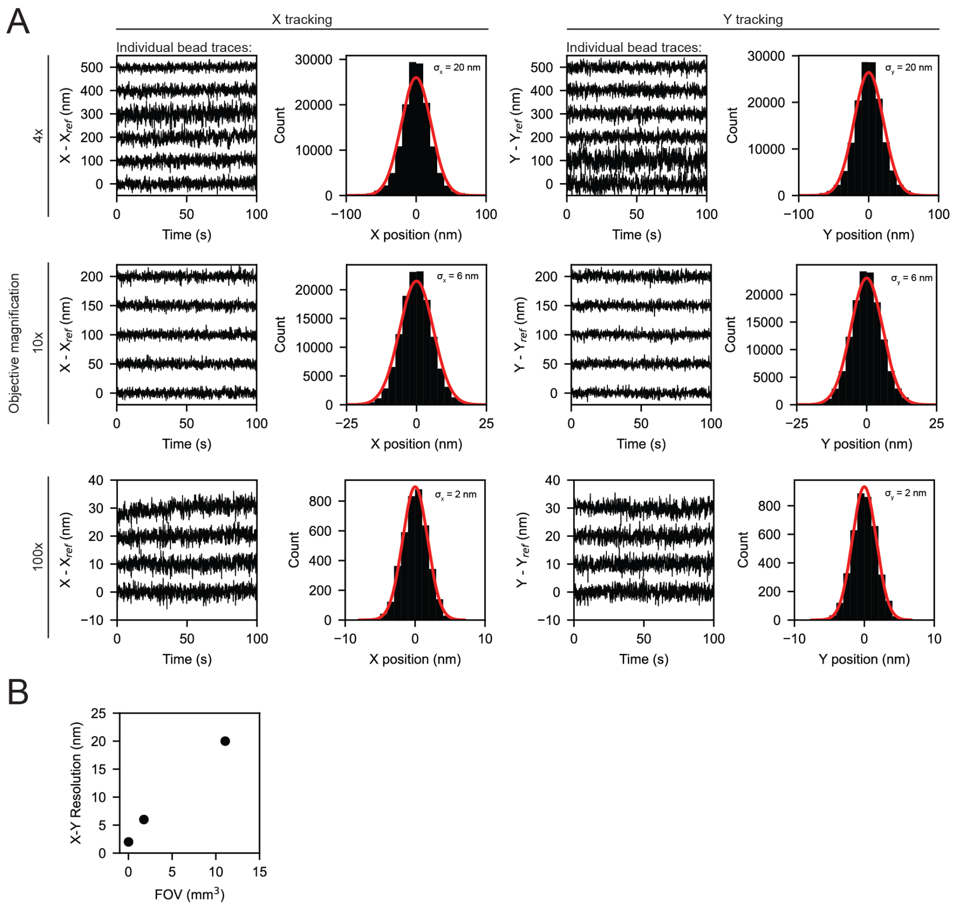


**Figure S1. X-Y position tracking of 1-micron beads quantifies the approximately linear tradeoffs between spatial resolution and FOV. A.** Measured *x* and *y* positions over time for 1-μm green fluorescent polystyrene beads stuck to glass surfaces imaged using 4x (top), 10x (middle), and 100x (bottom) objectives. *x* and *y* positions are given relative to the centroid of an arbitrary bead. Histograms show standard deviations in measured bead position relative to the centroid. Red line indicates Gaussian fit with mean = 0; annotation denotes fitted standard deviation. **B.** Measured resolution (standard deviation returned from Gaussian fits) *vs.* field of view (FOV) for 4x, 10x, and 100x objectives.

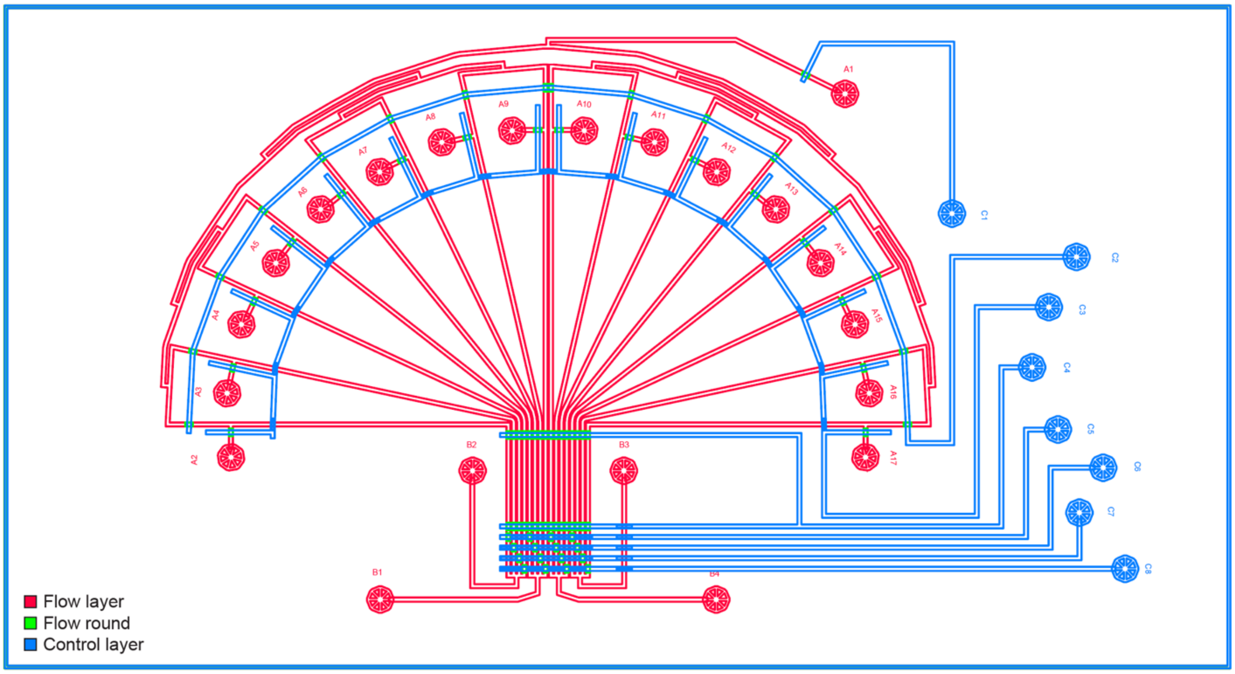


A1: General inlet
A2-A17: Lane-specific inlets

B1-B4: Outlets

C1: General inlet valve
C2: Backflow valve
C3: Lane valve
C4: Image valve
C5-C8: Binary outlet valves

**Figure S2. Schematic of the multichannel device.** Schematic shows different layers of photoresist required to fabricate molding masters for the flow (SU-8 2015, *red*; AZ50 XT, *green*) and control (SU-8 2025) molds. AutoCAD design files are available as Supplementary Files in a Zenodo online repository.

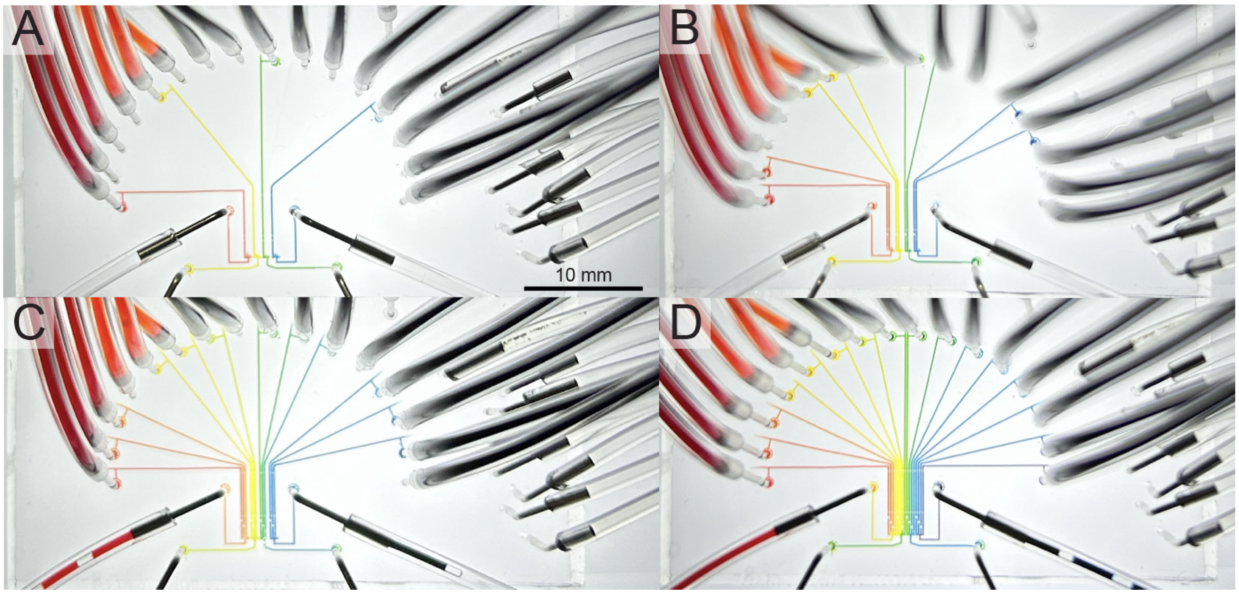


**Figure S3. Valve operation prevents cross contamination. A-D.** Images acquired over time when loading food coloring into channels individually. Even after loading only every other channel (as in A), interdigitated channels without food dye remain clear, confirming that valves at channel outlets prevent cross-contamination.

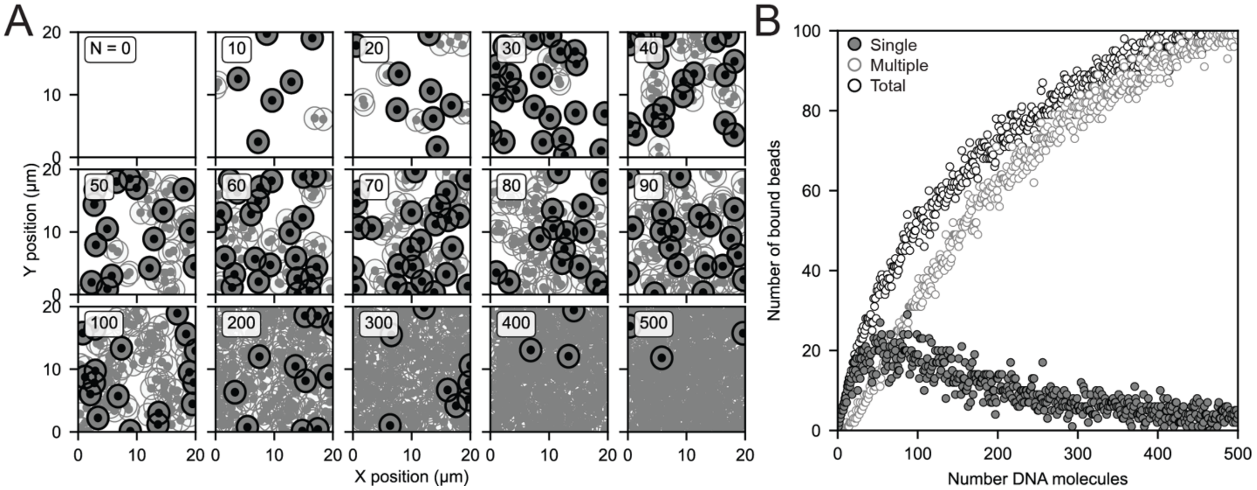


**Figure S4. Monte Carlo simulations provide a statistical model for random surface functionalization. A.** Example results of a single simulation in which we randomly placed *N* 4-kbp DNA molecules and 1-µm beads within a 20 µm x 20 µm 2D surface and counted whether beads overlapped with 1 or more DNA molecules. The center dot represents the attachment point of the tether, and the outer circle represents the constrained region of the tethered bead. Dark circles indicate single tethers and light circles indicate beads attached to more than one tether. **B.** Number of beads bound by one (*grey*, filled) or more than one (light grey, open) DNA molecules and total number of beads (*dark grey*, open) as a function of the number of placed 4-kpb DNA molecules across a total of 500 simulations.

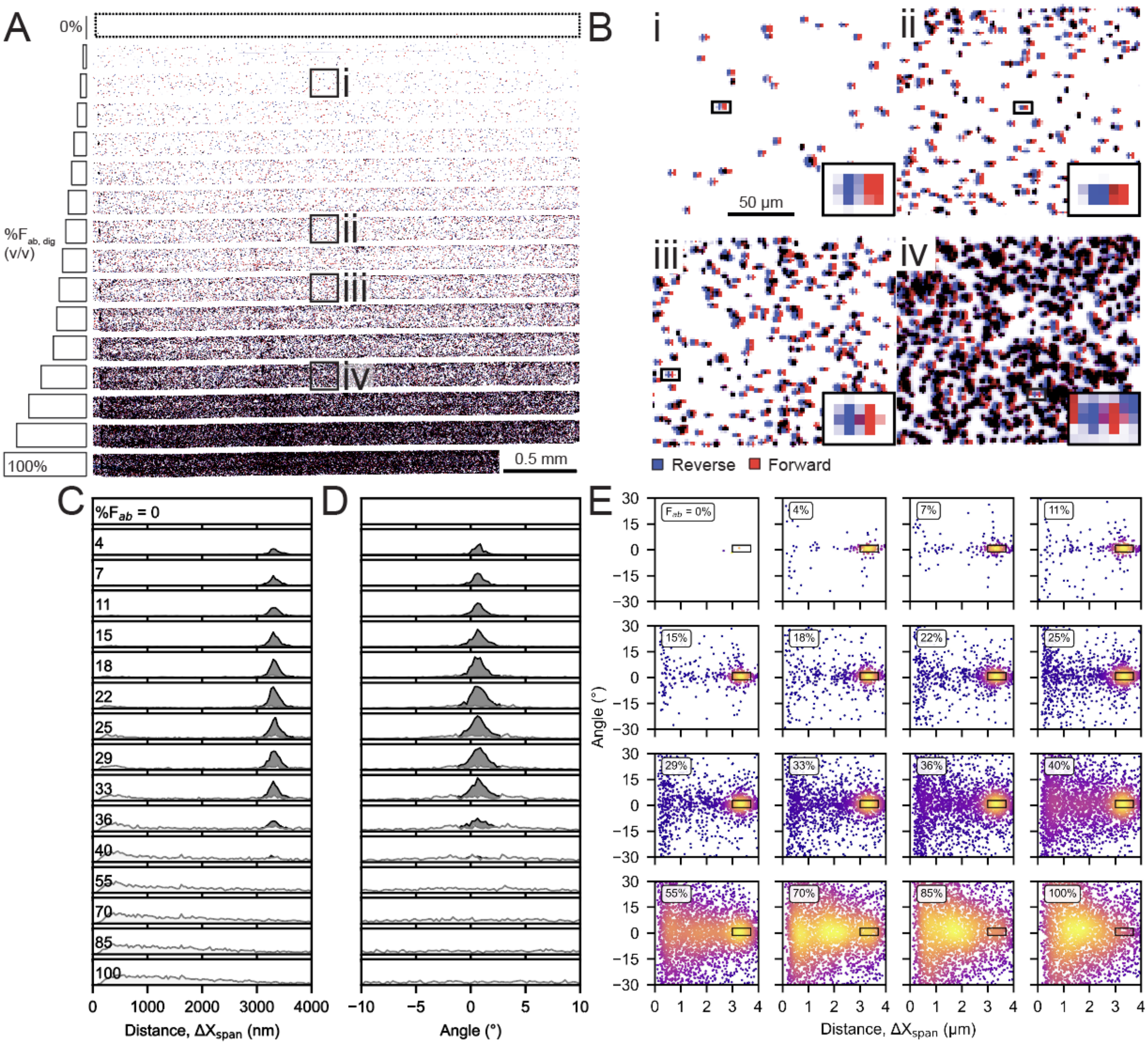


**Figure S5. A flow reversal assay quantifies the abundance of single versus multiple tether events. A.** Inverted fluorescence microscopy images showing beads immobilized by 4-kb DNA tethers across 16 channels patterned with increasing anti-digoxigenin Fab included during the initial patterning step. Labeled boxes (*i*,*ii*, *iii*, and *iv*) indicate the locations of zoomed images shown in (B). **B.** Zoomed-in inverted fluorescence microscopy images of regions indicated in A showing bead positions upon introduction of reverse (*blue*) or forward (*red*) flow. **C.** Histograms showing measured traversal distances of all beads upon flow reversal for each channel. **D.** Histograms showing measured traversal angles of all beads upon flow reversal for each channel. **E.** 2D density plots showing the measured traversal angle vs. the traversal distance for all beads as a function of the amount of anti-digoxigenin Fab within each panel; boxes indicate beads classified as tethered by a single molecule.

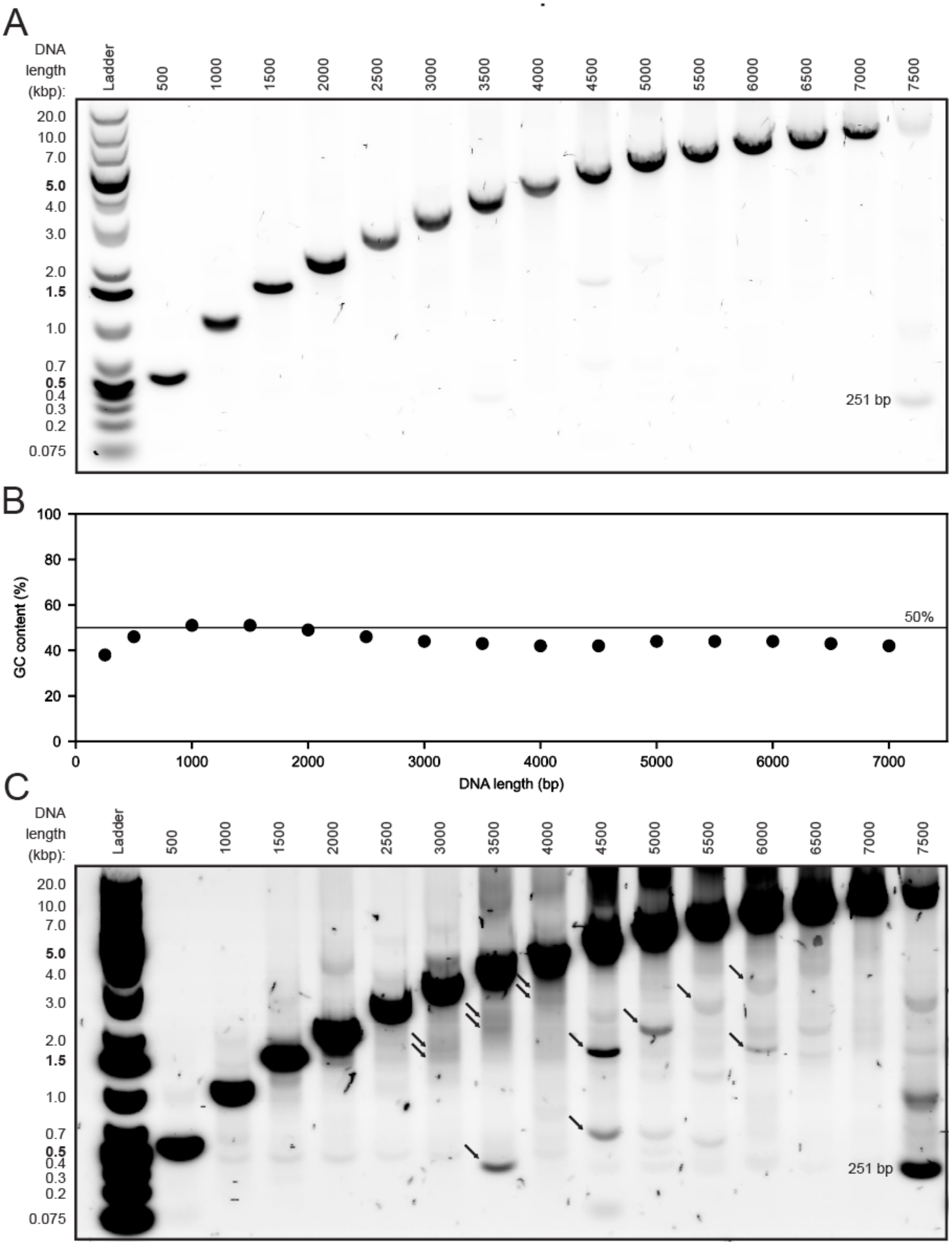


**Figure S6. Digoxigenin- and biotin- modified dsDNA constructs of varying lengths for multiplexed force-extension measurements. A.** DNA electrophoresis gel after PCR amplification of M13mp18 phage DNA using 15 primer pairs designed to amplify dsDNA of varying lengths. The 7500 bp construct did not amplify successfully and was excluded from downstream analysis. **B.** GC content *vs.* amplicon length for amplified dsDNA constructs. **(C)** Identical gel as in (A) but with image contrast enhanced to show the presence of additional, shorter bands for some constructs.

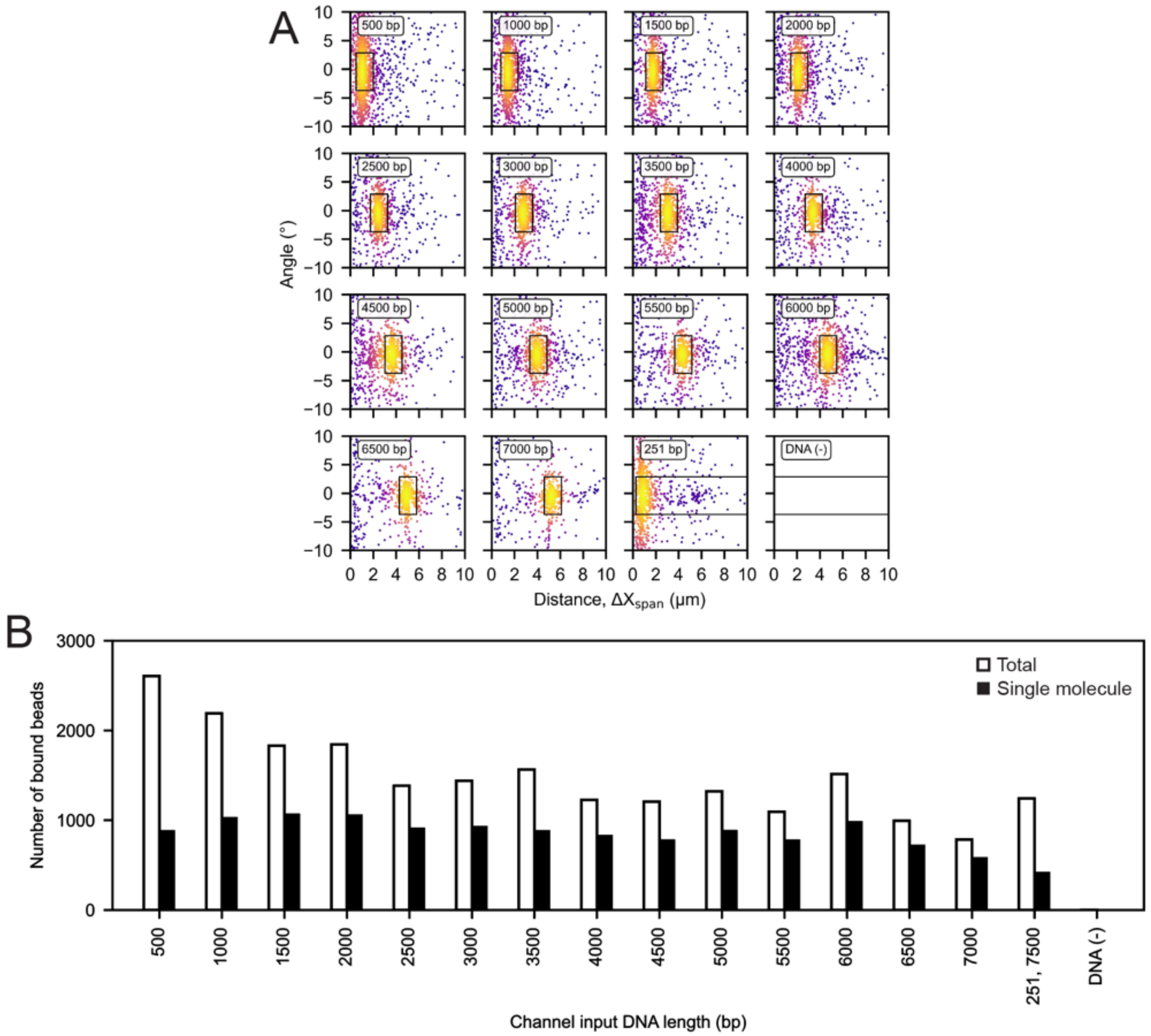


**Figure S7. Quantifying singly- and multiply-tethered beads per channel during force-extension experiments (1-μm diameter beads). A.** Density plots of measured traversal angle *vs.* traversal distance for all beads detected in each channel; boxes indicate beads classified as tethered by a single molecule. **B.** Total (white bars) and single-molecule-tethered (black bars) beads for channels containing dsDNA constructs of the indicated length.

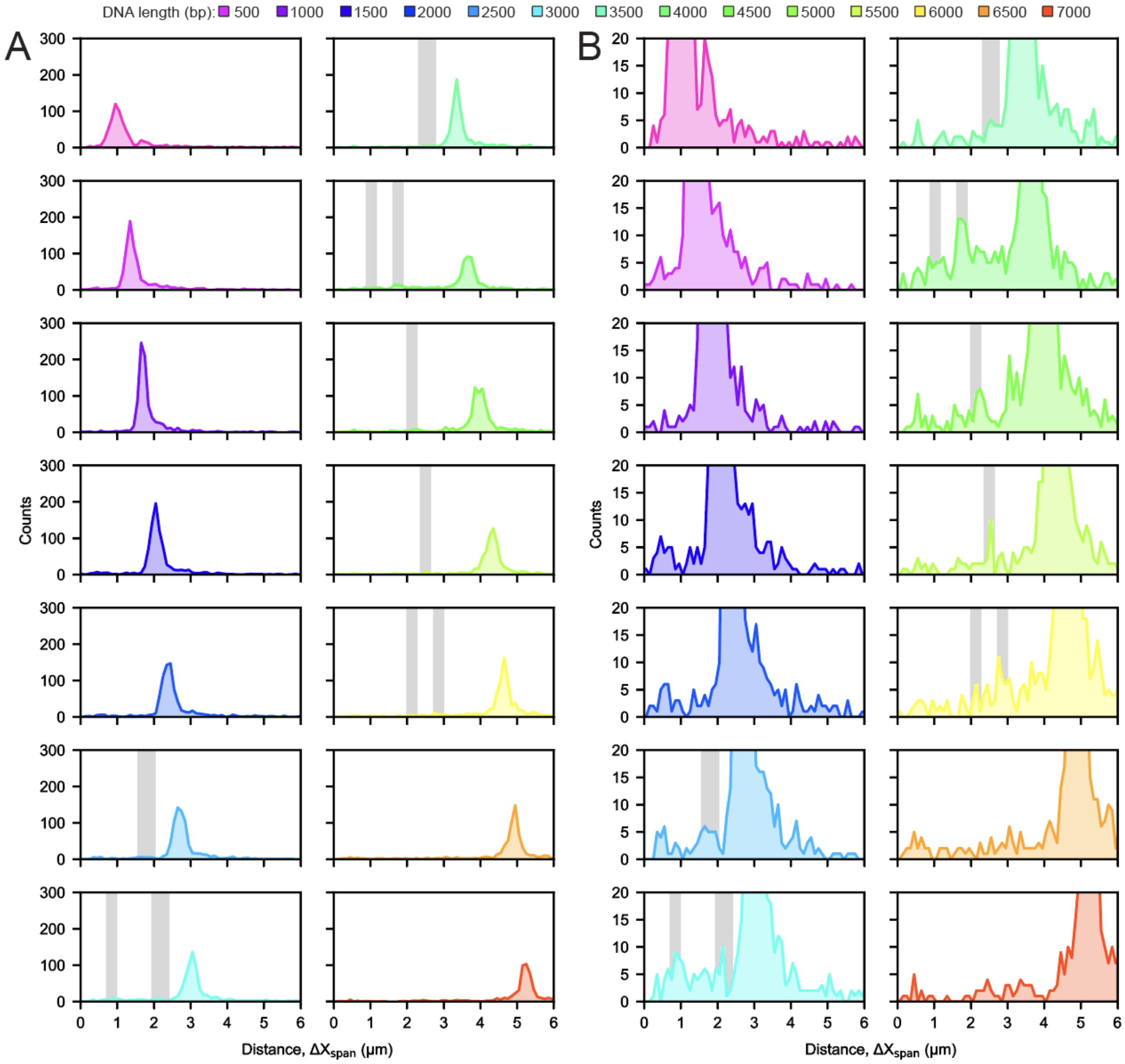


**Figure S8. Multiplexed stretching can detect rare species. A.** Distributions of measured traversal distances for channels containing beads tethered by dsDNA amplicons of different lengths; grey bars denote the expected distances for shorter bands labeled with arrows in Supplementary Figure S6C. **B.** The same distribution shown in (A) with the *y* axis maxima changed to show rare species.

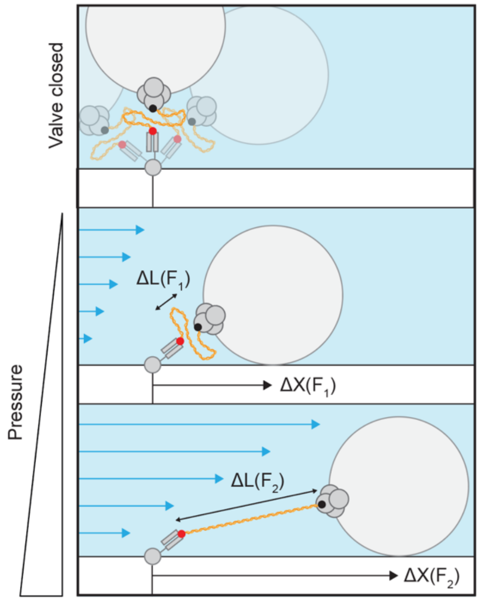


**Supplementary Fig. 9. Schematic of the geometric model relating molecular extension and bead displacements**. With valves closed (top), there is no flow. Tethers constrain beads to diffuse around the attachment point such that fitting the X-Y position to a 2-D Gaussian approximates the tether attachment point on the channel. Upon opening valves and applying pressure to drive flow, the bead is displaced a distance $\Delta X$ and a geometric model relates molecular extension $\Delta L$ to observed displacement (**Supplementary Note 2**).

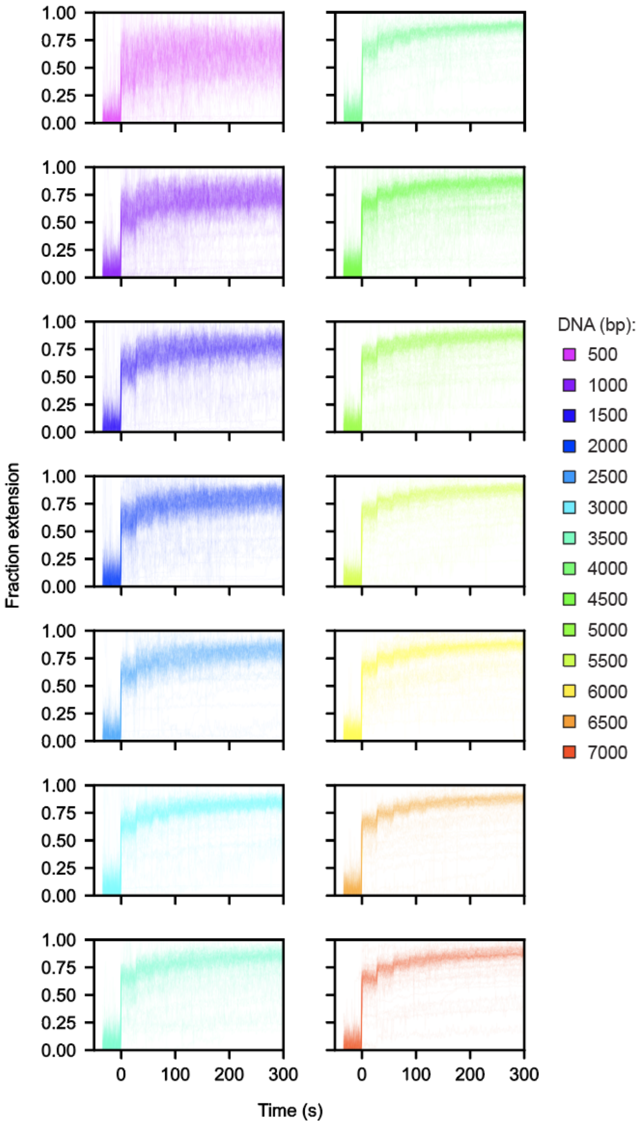


**Figure S10. Individual traces of DNA stretching.** All force-extension curves (calculated fractional molecular extension *vs.* applied pressure) for single-molecule dsDNA tethers of varying lengths. Valves were closed prior to the pressure step ramp and imposed a zero-flow condition. The pressure step ramp started at 0 seconds.

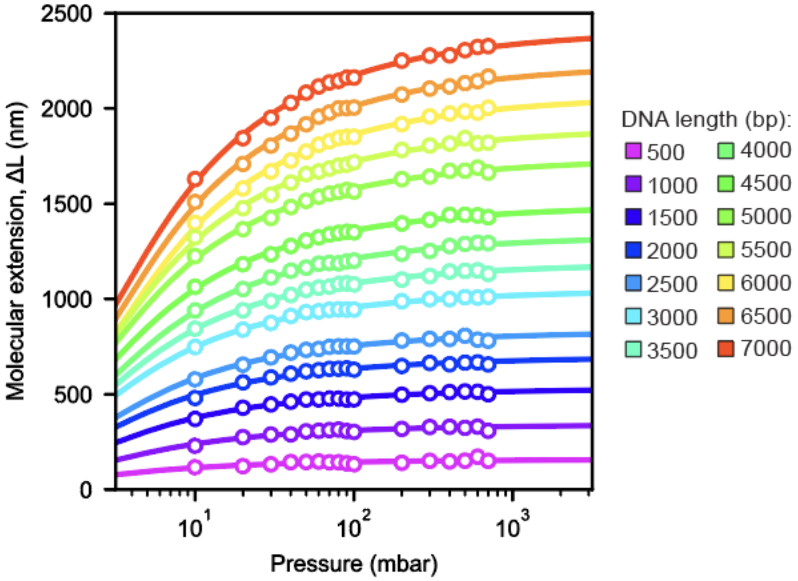


**Figure S11. Log-scale force-extension curve of DNA demonstrating the ability to resolve applied forces that differ by 0.25 pN.** Measured molecular extension as a function of applied pressure for single-molecule tethers of varying lengths. Markers denote calculated molecular extension after iteratively increasing pressure in 10 or 100 mbar steps; lines indicate WLC fits to the data. Across all tether lengths, increasing pressure from 10 to 20 mbar results in easily detectable changes in molecular extension that matched the WLC model fit. Pressure-tension calibration yielded $m = 25$ pN bar$^{-1}$ (**Fig. 3G**, **Supplementary note 1**); a resolvable pressure step of 10 mbar is equal to 0.25 pN.

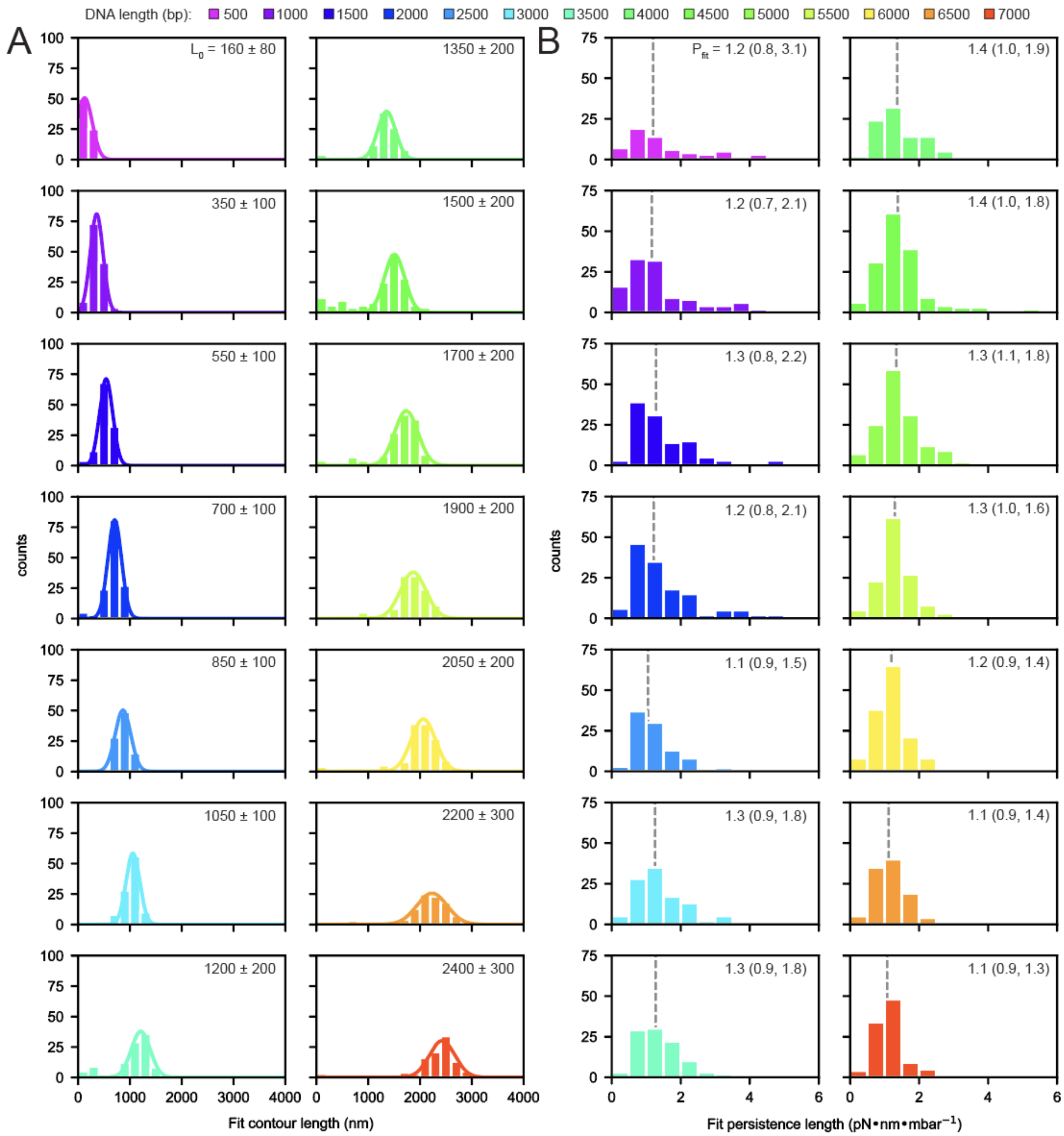


**Figure S12. WLC parameter fit distributions. A.** Distributions of fitted contour lengths for single-molecule dsDNA tethers of increasing lengths; line indicates median fit values and annotations denote fitted mean contour length and standard deviation. **B.** Distributions of the fitted persistence lengths and their log-normal fits for single-molecule dsDNA tethers of increasing lengths; line indicates median and annotations denote median fit persistence length and the interquartile range in parentheses. Fitted persistence lengths calculated the tension-to-pressure relationship (see **Fig. 3X, Supplementary Note 1).**

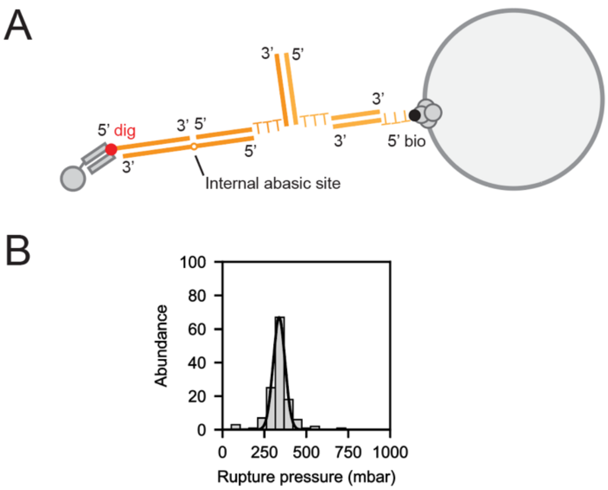


**Figure S13. Unzipping DNA duplex as an orthogonal flow-tension calibration. A.** Attachment geometry for irreversible unzipping of DNA duplexes. Schematic showing a bead attached via a representative dsDNA duplex used for iterative hybridization and unzipping (not drawn to scale). A dsDNA fragment displaying one half of the duplex to be unzipped is anchored to the surface via a digoxigenin/Fab linkage and a dsDNA displaying the other half of the duplex is anchored to a bead via a biotin/streptavidin linkage. **B.** Unzipping pressure measurements of a 15-bp DNA duplex attached to a 7-kbp tether and 1-micron bead. A Gaussian fit yielded a most probable rupture pressure of 340 $\pm$ 30 mbar.

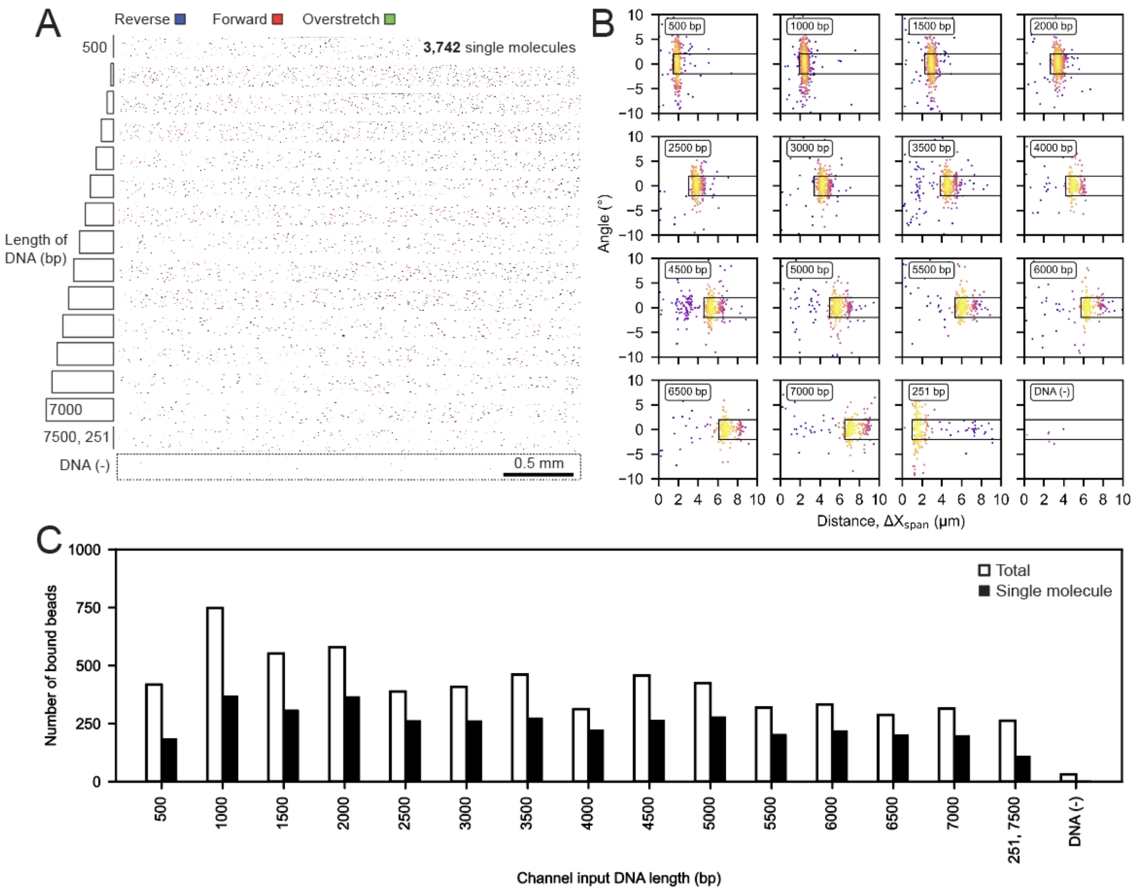


**Figure S14. Quantifying singly- and multiply-tethered beads per channel during overstretching experiments (3-μm diameter beads). A.** Inverted fluorescence microscopy images showing beads immobilized by dsDNA tethers of increasing lengths across 16 channels. **B.** Density plots of measured traversal angle *vs.* traversal distance for all beads detected in each channel; boxes indicate beads classified as tethered by a single molecule. **B.** Total (white bars) and single-molecule-tethered (black bars) beads for channels containing dsDNA constructs of the indicated length.

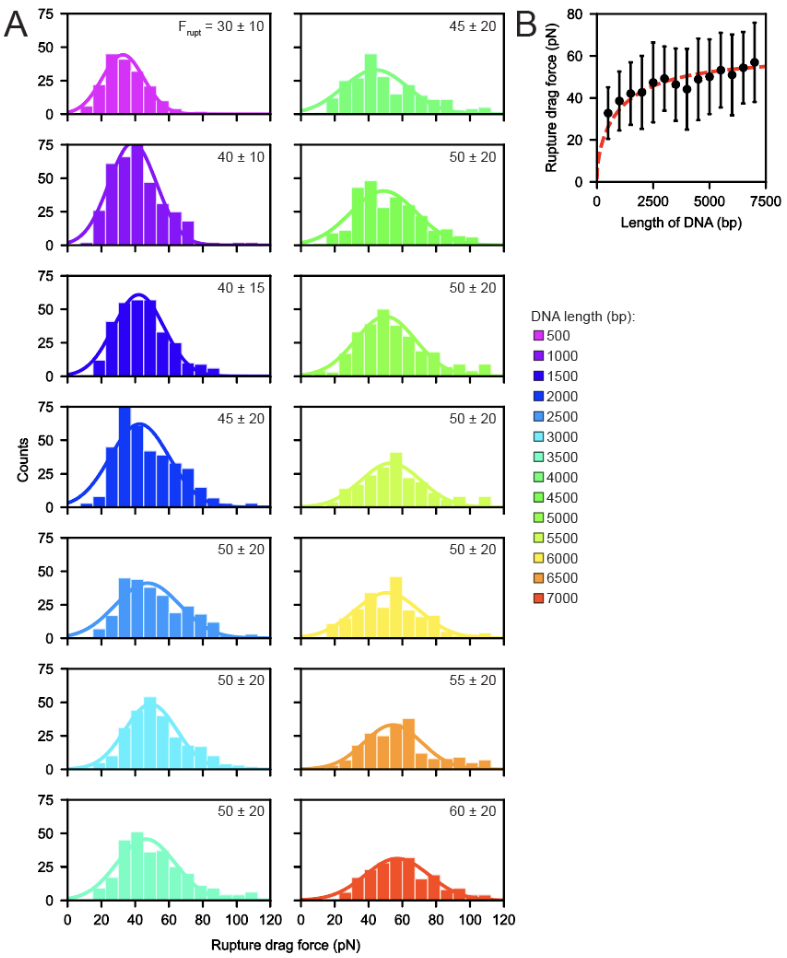


**Figure S15. Rupture force histograms. A.** Distributions of drag forces at the time of bead rupture. Line indicates a Gaussian fit to determine the most probable rupture force; annotation denotes Gaussian mean and standard deviation. The ramp rate of drag force was ~2.2 pN $s^{-1}$. **B.** Rupture force as a function of dsDNA tether length. Markers and error bars denote fitted Gaussian mean and standard deviation, respectively. Red dashed line indicates expected rupture forces calculated based on a geometric model. Measured rupture events likely result from rupture of the weakest linkage of the tether (likely the Fab:digoxigenin linkage).

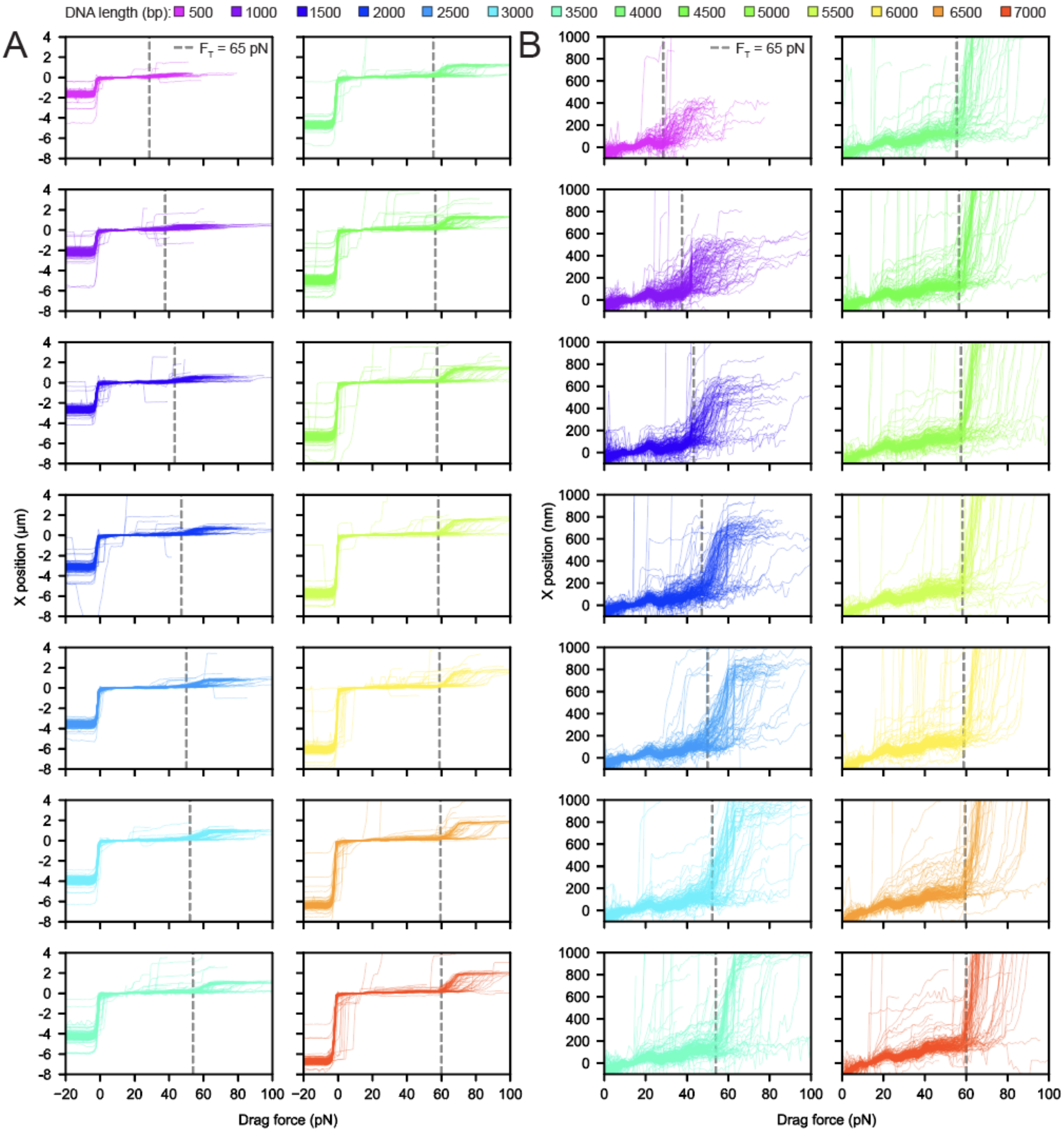


**Figure S16. Flow reversal and zoomed-in views of parallelized overextension. A.** Measured displacement (in microns) as a function of drag force for beads tethered by single dsDNA duplexes of varying lengths. The transition from negative to positive forces occurs upon flow reversal near the start of the assay. **B.** Measured displacement (in nanometers) as a function of drag force just after flow reversal for beads tethered by single dsDNA duplexes of varying lengths; zoomed in view highlights detected overextension events. In A and B, grey dashed lines indicate an expected tension of 65 pN calculated from the geometric model.

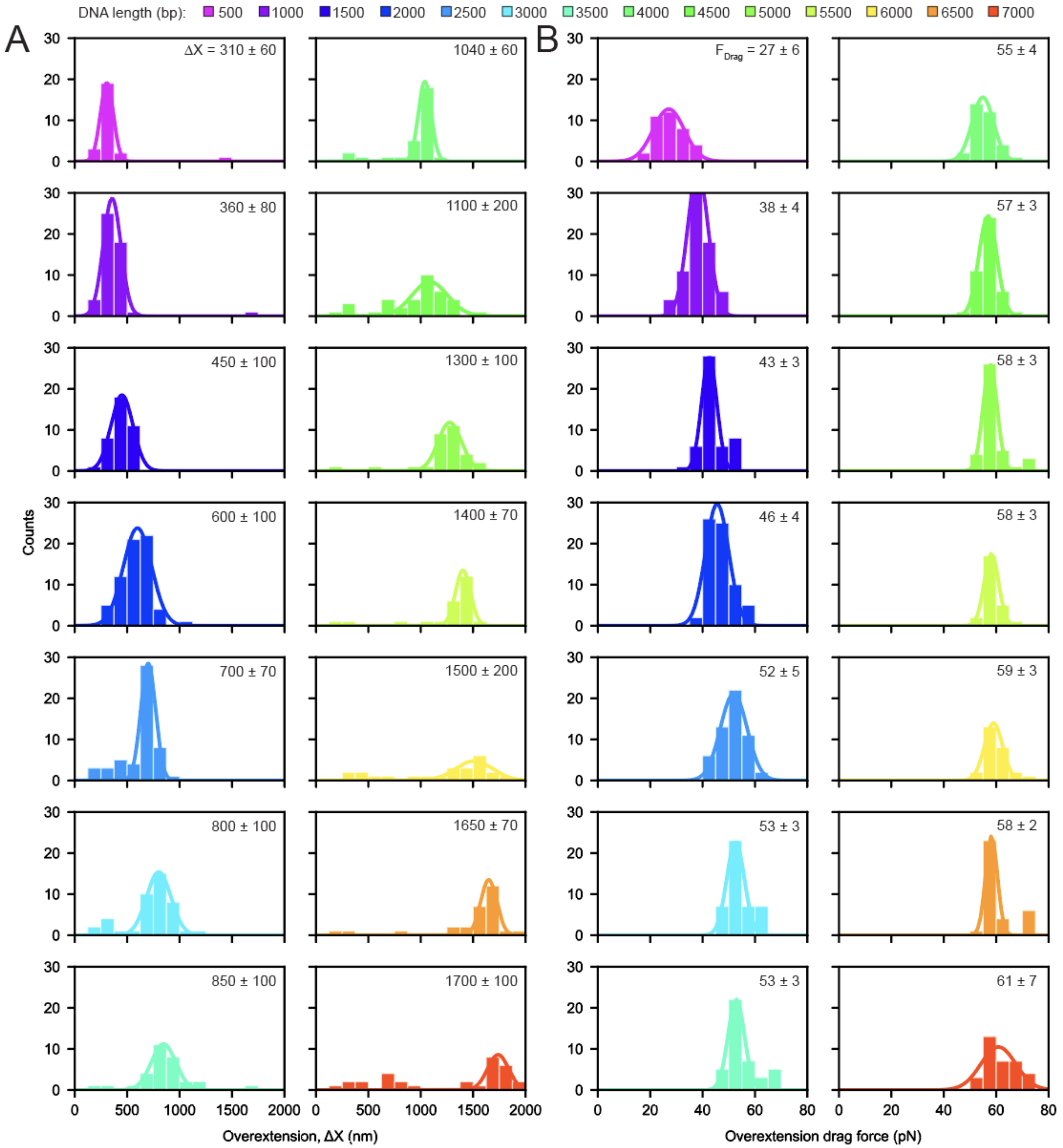


**Figure S17. Overextension distance and force histograms.** Measured distributions of displacements (**A**) and calibrated drag forces (**B**) at overextension transitions for beads tethered by single dsDNA duplexes of varying lengths. Lines indicate Gaussian fit with annotated fit parameters.

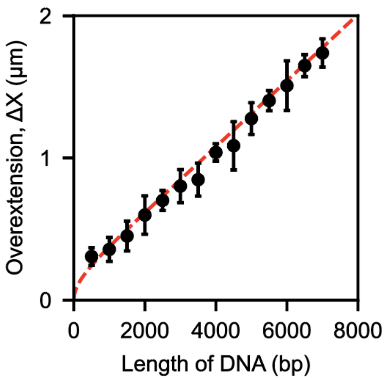


**Figure S18. Overextension distances.** Measured overextension displacement *(ΔX)* as a function of DNA tether length. Error bars denote standard deviations; red dashed line indicates expected results for the geometric model.

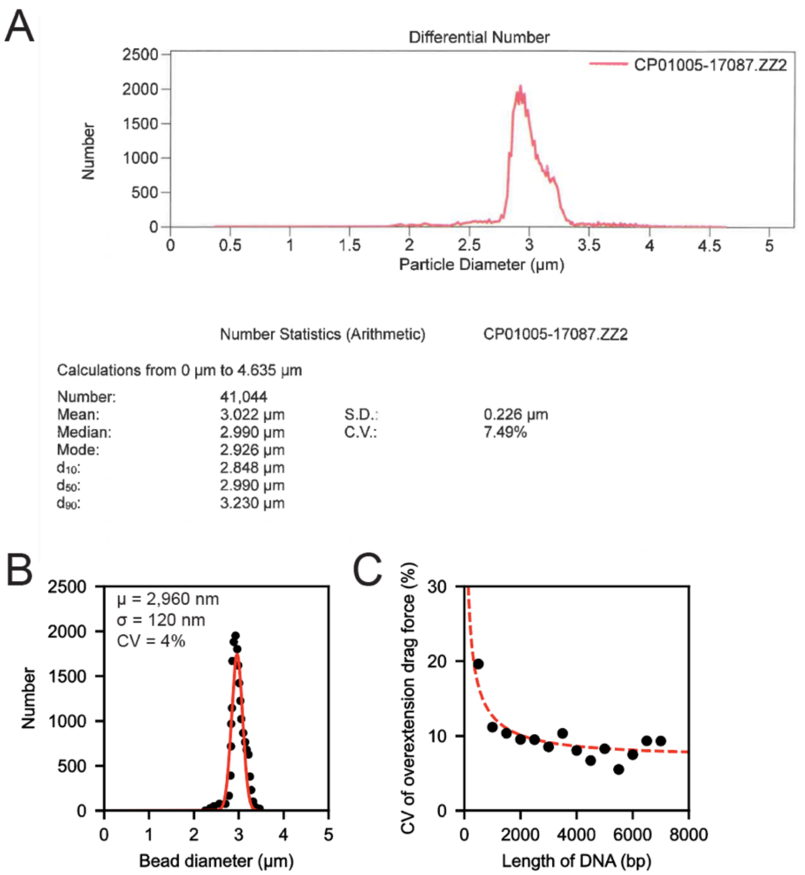


**Figure S19. Variation of overextension drag forces decreases with tether length and asymptotically approaches $2\cdot \delta R\, R^{-1}$. A.** Bead diameter distribution provided by Bangs Labs. **B.** Extracted bead diameter distribution (black markers) from S19A along with a Gaussian fit (red line); annotation indicates bead diameter mean (µ), standard deviation (σ), and coefficient of variation (CV). **C.** Measured coefficient of variation (CV) of the drag force required for overextension as a function of dsDNA tether length (black markers); red line indicates expected CV based on the geometric model (CV = $2\cdot \delta R\, R^{-1} \cos(\alpha)^{-1}$) that considers how drag force is amplified on short tethers by a factor of $\cos(\alpha)^{-1}$. The least squares fit yielded a coefficient of variation in drag force of *7.3%* in the limit of long tethers. The least squares fit for the coefficient of variation of drag force agreed with the measured variance of bead diameter, $2\cdot CV \sim 8\%$.

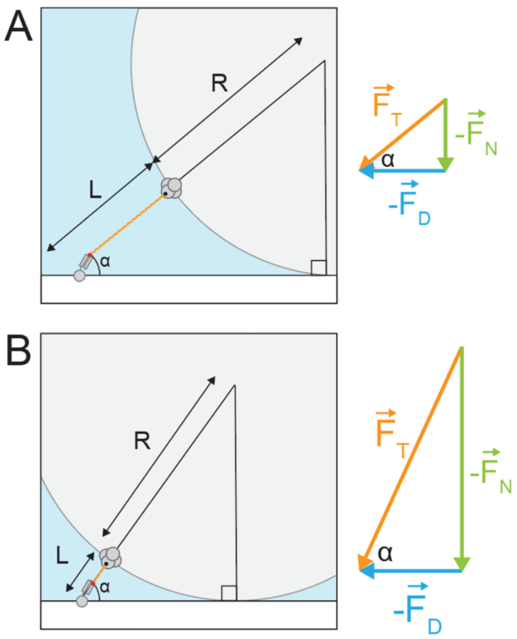


**Figure S20. Tension is a function of drag force and molecular geometry. A.** Schematic showing relevant geometry (left) and force body diagrams (tension force ($F_T$, orange), normal force ($F_N$, green), and drag force ($F_D$, blue); right) for a tether only slightly smaller than the bead radius. **B.** Schematic showing relevant geometry (left) and force body diagram (right) for a tether much smaller than the bead radius. As the tether shortens relative to the bead radius, the magnitudes of the tension and normal force vectors increase but the magnitude of the drag force vector (blue) stays constant. See Supplementary Note 2.

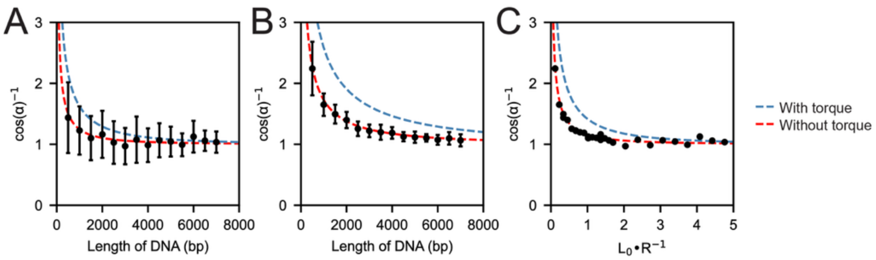


**Figure S21. Geometric models without torque predict the molecular geometries calculated from DNA stretching and overstretching experiments. A.** Measured $cos(\alpha)^{-1}$ scaling factor (black markers) for 1-μm green fluorescent beads tethered by single dsDNA tethers of varying lengths. Error bars denote standard deviation; dashed lines indicate predictions from models with (blue) and without (red) torque. **B.** Same plot as in A for 3-μm beads. **C.** Measured $cos(\alpha)^{-1}$ scaling factor (black markers) for 1-μm and 3-μm beads *vs.* the dimensionless geometric variable $\mathrm{K} = \Delta L \cdot R^{-1}$ (**Supplementary Note 2**).

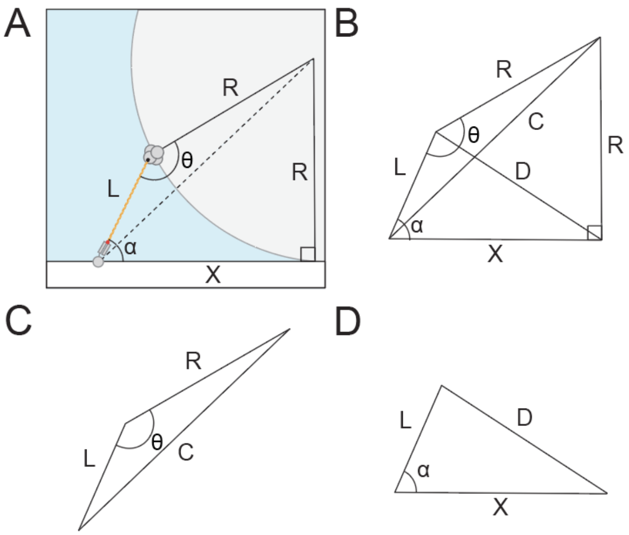


**Figure S22. Influence of hydrodynamic torque on molecular geometry.** See Supplementary Note 2.

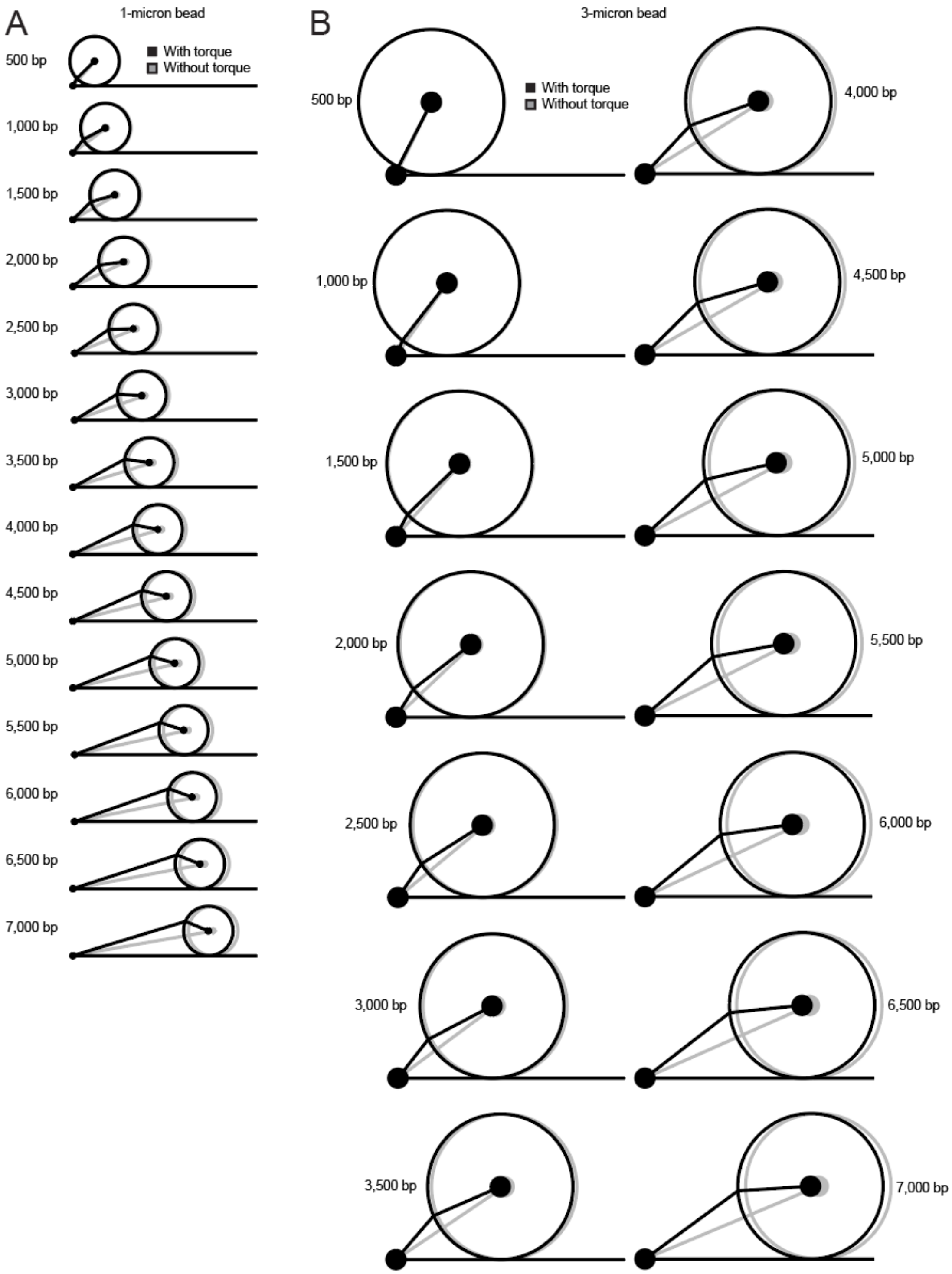


**Figure S23. Expected effect of torque on molecular geometry. A.** Schematic of expected geometries under force for a 1-micron bead in the presence (black) and absence (grey) of torque. When torque is included in the geometric model, the bead rotates and moves upstream relative to the no-torque model (see **Supplementary Note 2**). **B.** The same schematic and calculation for a 3-micron bead. The molecular geometry is parameterized by the dimensionless geometric variable $\mathrm{K} = L \cdot R^{-1}$ such that the geometry in (B) is simply scaled from (A). See Supplementary Note 2.

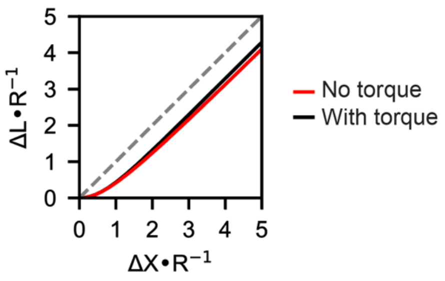


**Figure S24. Predicted relationship between molecular extension and bead displacement for geometric models with and without torque.** Predicted bead radius-normalized molecular extension ($\Delta L$) *vs.* bead displacement ($\Delta X$) for models with (black line) and without (red line) torque; dashed grey line indicates the one-to-one line. See Supplementary Note 2.

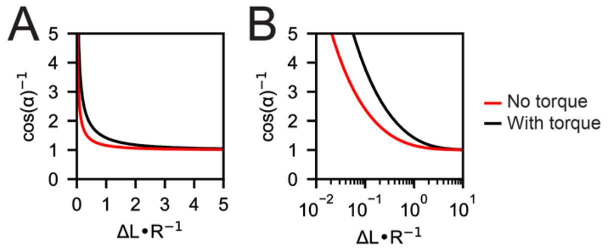


**Figure S25. Predicted scaling factor between drag and tension force (*cos(α)*$^{-1}$) for geometric models with and without torque. A.** Predicted *cos(α)*$^{-1}$ scaling factor between drag and tension forces for no torque (red line) and with torque (black line) models as a function of radius-normalized tether length ($\Delta L$) shown on a linear scale. **B.** Same plot as in (A) shown on a log scale. The models agree for tethers that are ~2-fold longer than the radius of the bead. See Supplementary Note 2.

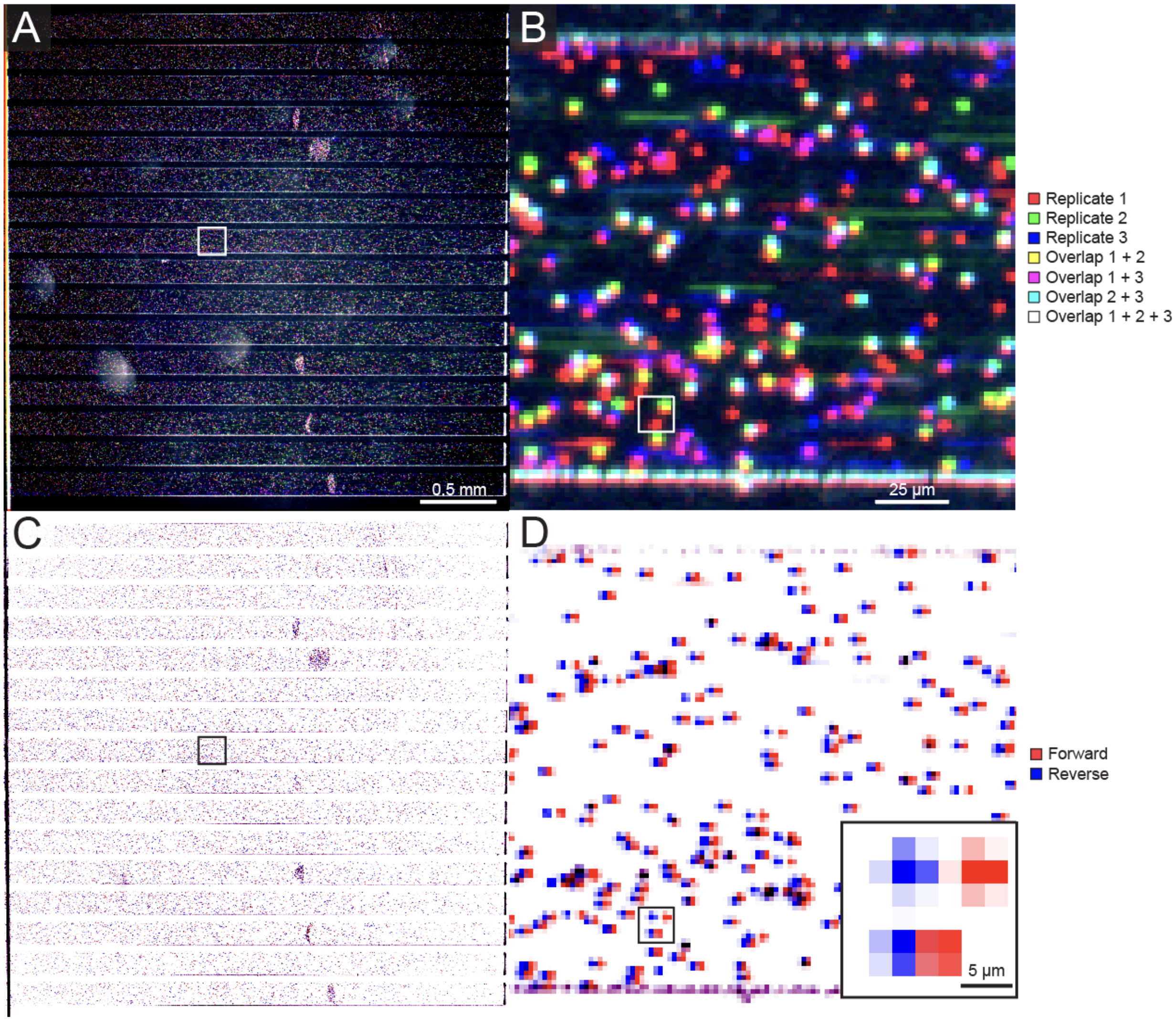


**Figure S26. Overlapping locations of bead binding during iterative binding on the same chip. A.** Aligned fluorescence images of a 16-channel device showing positions of beads after 3 iterative rounds of hybridization in red (replicate #1), green (replicate #2), and blue (replicate #3). White box indicates zoomed in area shown in B. **B.** Zoomed in and aligned fluorescence image after 3 iterative rounds of hybridization. Bead positions that overlapped between hybridization rounds are indicated in yellow, purple, cyan, and white (see legend). **C.** Inverted images showing bead positions during two frames of a flow reversal measurement in Replicate 1. **D.** Zoomed in and aligned inverted image The inset region in (C) is the same as (A) and (B). The bead displacement indicated that the top and bottom beads were attached to a 7-kbp and 3-kbp tether, respectively.

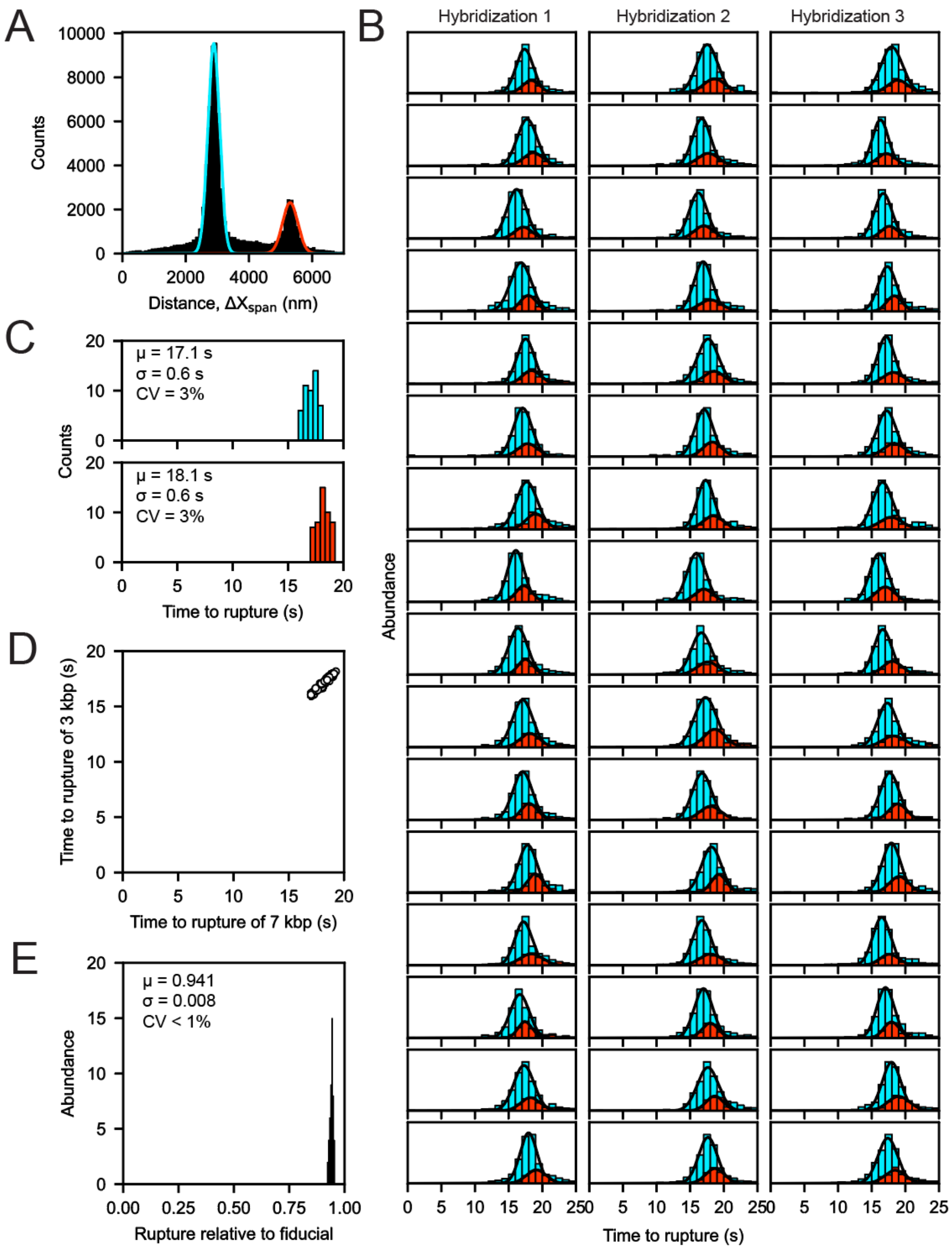


**Figure S27. Mechanical demultiplexing and calibration of fiducials between 3- and 7-kbp tethers on a single device. A.** The distribution of bead displacements showed two populations of tethers corresponding to 3-kbp (blue) or 7-kbp (red) tethers. The same GC47N15 duplex was formed on both tethers. **B.** Three consecutive rupture force measurements on the same chip. Rupture events were categorized by the identity of their tether as determined in (A). **C.** Distribution of mean time to rupture across all channels and rounds of hybridization. The longer 7-kbp tether (red) took longer to rupture on average than the shorter 3-kbp tethers (blue). **D.** Time to rupture of the shorter vs. longer tether beads from the same channel were correlated. **E.** Normalization of the time to rupture of the short tether relative to the long reduced the coefficient of variation and indicated that the short tethers ruptured earlier than the long tethers.

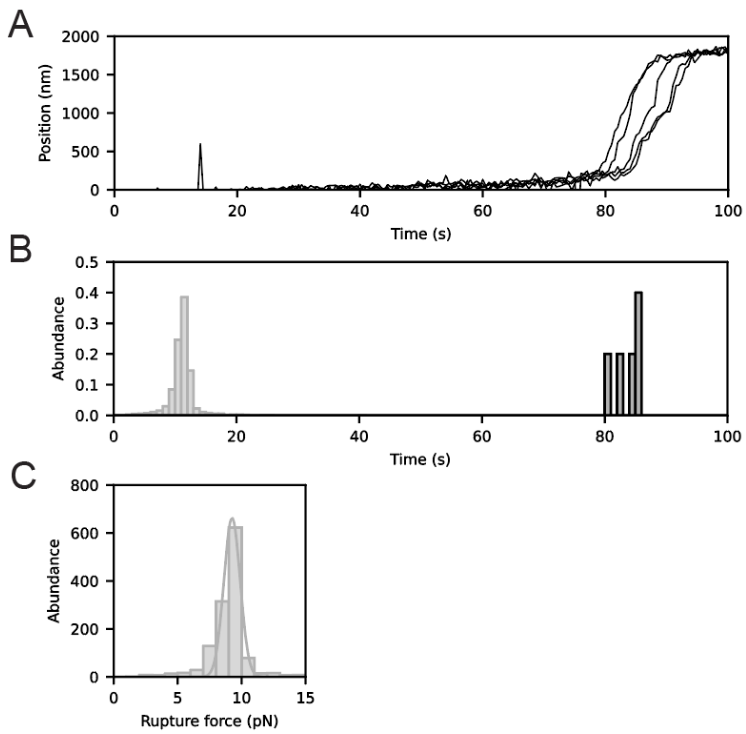


**Figure S28. Calibrated unzip measurement of the 15-bp fiducial duplex. A.** Bead position as a function of time for 1-$\mu$m dynabeads tethered by a biotin- and digoxigenin-modified bifunctional 7-kbp dsDNA tether construct lacking an unzipping duplex fragment. The sharp transition around and just after 80 s corresponds indicates the B-to-S transition, providing an independent measure of when 65 pN of tension was applied to the tethers. **B.** Histograms of the times at which 1-µm Dynabeads tethered by single molecules either ruptured or displayed an expected B-S transition, Light grey bars indicate times of rupture/unzipping events for beads tethered by a 3-kbp dsDNA fragment displaying the GC47N15 ‘unzipping’ construct; dark grey bars indicate times of an observed B-to-S transition for beads tethered by 7-kbp dsDNA. Measurements were made in basic microfluidic flow chamber devices that have lower resistance than the multichannel device, enabling measurement of both unzipping and overextension with 1-micron beads. AutoCAD design files are available as Supplementary Files in a Zenodo online repository. **C.** Histogram of calibrated unzipping force measurements of the GC47N15 duplex relative to the B-to-S transition force. Line indicates a gaussian fit with a mean rupture force of 9.2 $\pm$ 0.6 pN.

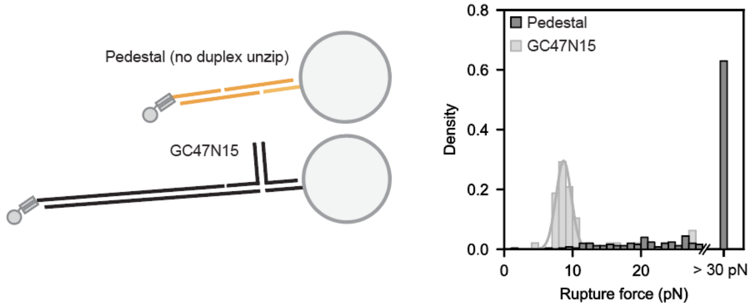


**Figure S29. Control experiments establishing that the strength of the surface attachments exceeded 30 pN. A.** Schematic illustrating a representative dsDNA duplex used for iterative hybridization and unzipping (‘GC47N15’, bottom) and a control duplex used to test the strength of surface attachment linkages (‘pedestal’, top). **B.** Histogram showing the frequency of measured rupture forces for ‘pedestal’ and ‘GC47N15’ constructs. Most single-molecule “pedestal” tethers (161 of 251) did not rupture when ramping to 30 pN at a rate of 0.5 pN $s^{-1}$.

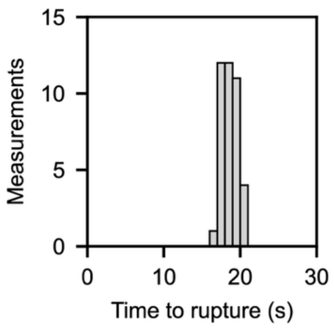


**Figure S30. Reproducibility of time to unzipping for fiducial unzipping construct across four devices.** Histogram showing time to rupture measurements across four devices for the same GC47N15 duplex attached to a 7-kbp tether. The mean time to rupture was 19 $\pm$ 1 seconds (CV = 5%).

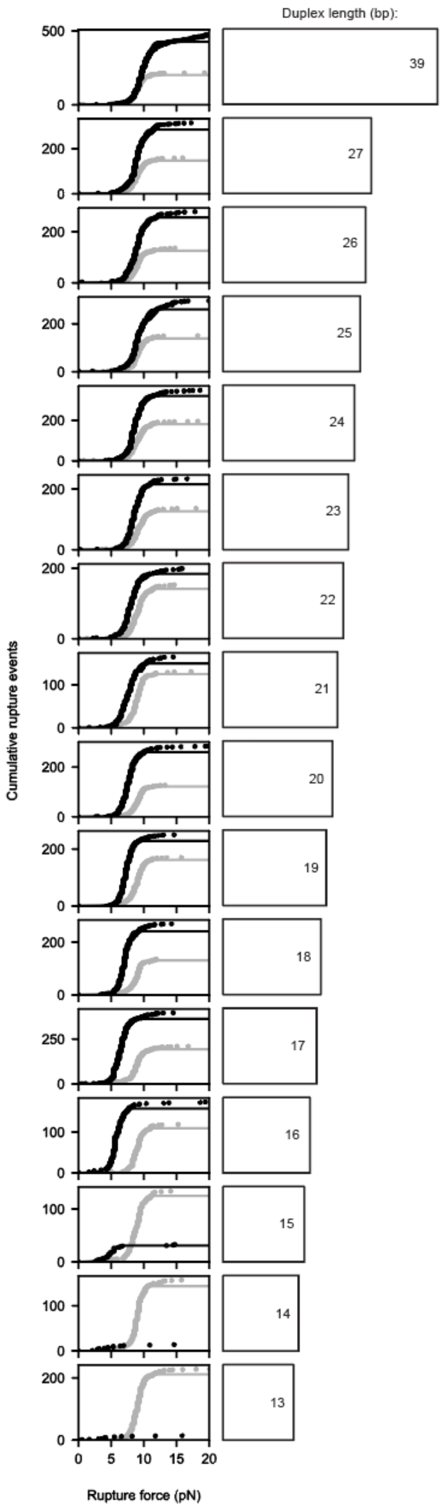


**Figure S31. Cumulative rupture force distributions and fits for 100% AT duplexes.** Cumulative rupture force distributions for variable duplexes (black data points) and ‘fiducial’ duplexes (grey data points). Horizontal bars (right) indicate length of the measured duplex. Bell-Evans model fits for all data are shown in black or grey for variable and fiducial duplexes, respectively.

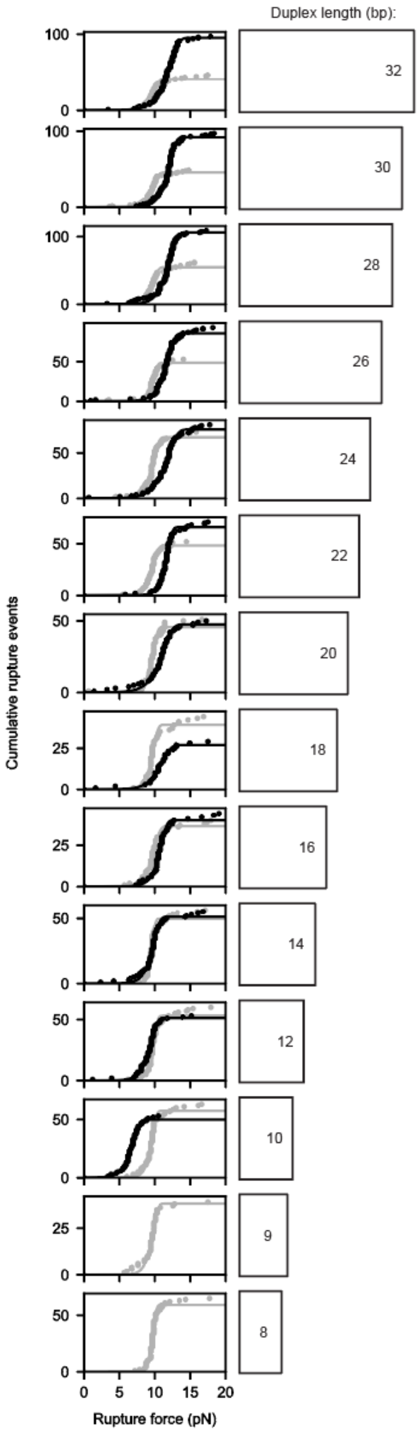


**Figure S32. Cumulative rupture force distributions and fits for 50% GC duplexes.** Cumulative rupture force distributions for variable duplexes (black data points) and 'fiducial' duplexes (grey data points). Horizontal bars (right) indicate length of the measured duplex. Bell-Evans model fits for all data are shown in black or grey for variable and fiducial duplexes, respectively.

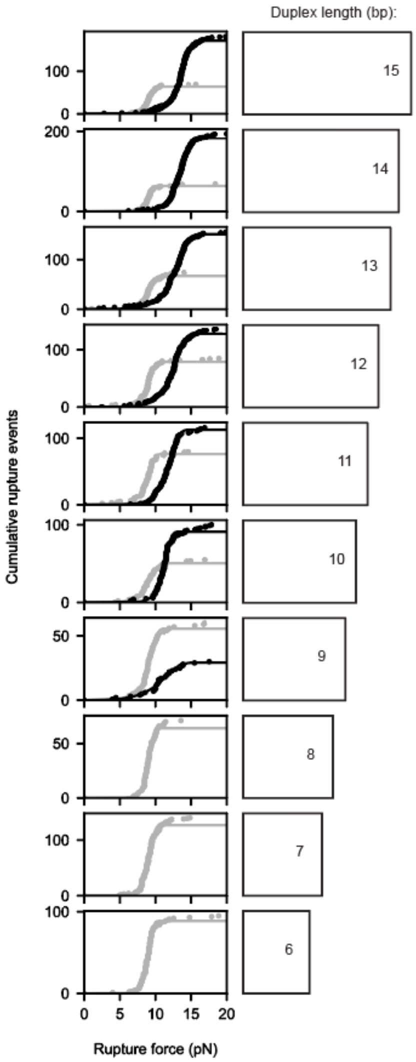


**Figure S33. Cumulative rupture force distributions and fits for 100% GC duplexes.** Cumulative rupture force distributions for variable duplexes (black data points) and 'fiducial' duplexes (grey data points). Horizontal bars (right) indicate length of the measured duplex. Bell-Evans model fits for all data are shown in black or grey for variable and fiducial duplexes, respectively.

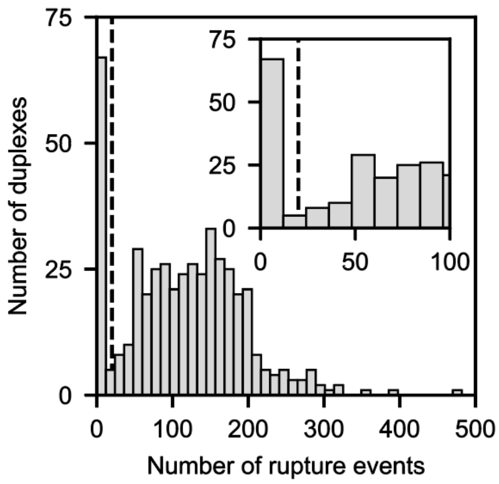


**Figure S34. Single-molecule observation histogram for duplex unzipping measurements.** Histogram showing the number of duplexes with a given number of rupture events across the standard DNA duplex library. Dashed line indicates the single-molecule event threshold (N = 20 single-molecule rupture events) for classifying rupture events as DNA duplex unzipping; duplexes with 0 rupture events corresponded to duplexes that were not sufficiently stable to tether beads to surfaces.

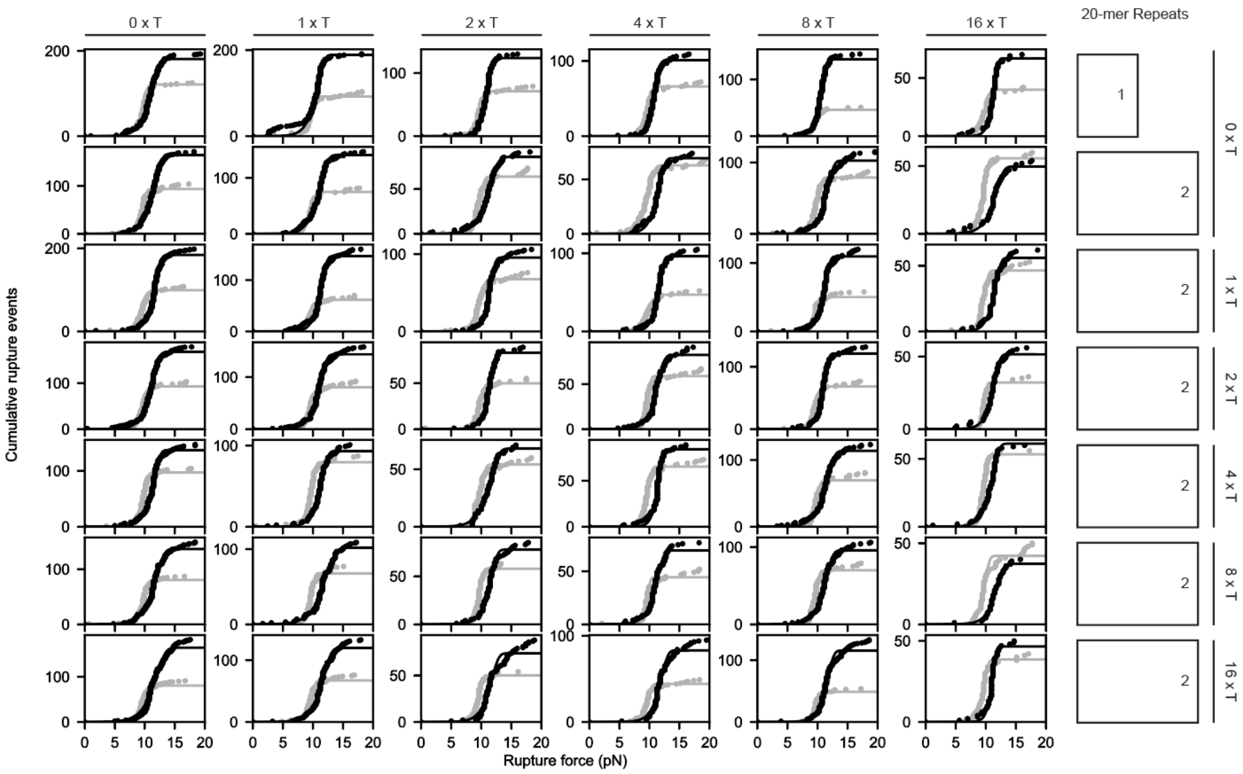


**Figure S35. Cumulative rupture force distributions and fits of 20-mer multivalent duplexes.** Cumulative rupture force distributions for variable duplexes (black data points) and 'fiducial' duplexes (grey data points). Horizontal bars (right) indicate number of duplex repeats. Linker sizes are annotated on the edges of the distributions: right labels indicate length of linker A, and top labels indicate length of linker B. Bell-Evans model fits for all data are shown in black or grey for variable and fiducial duplexes, respectively. The 1 x 20-bp construct corresponds to a single 20-bp unit duplex.

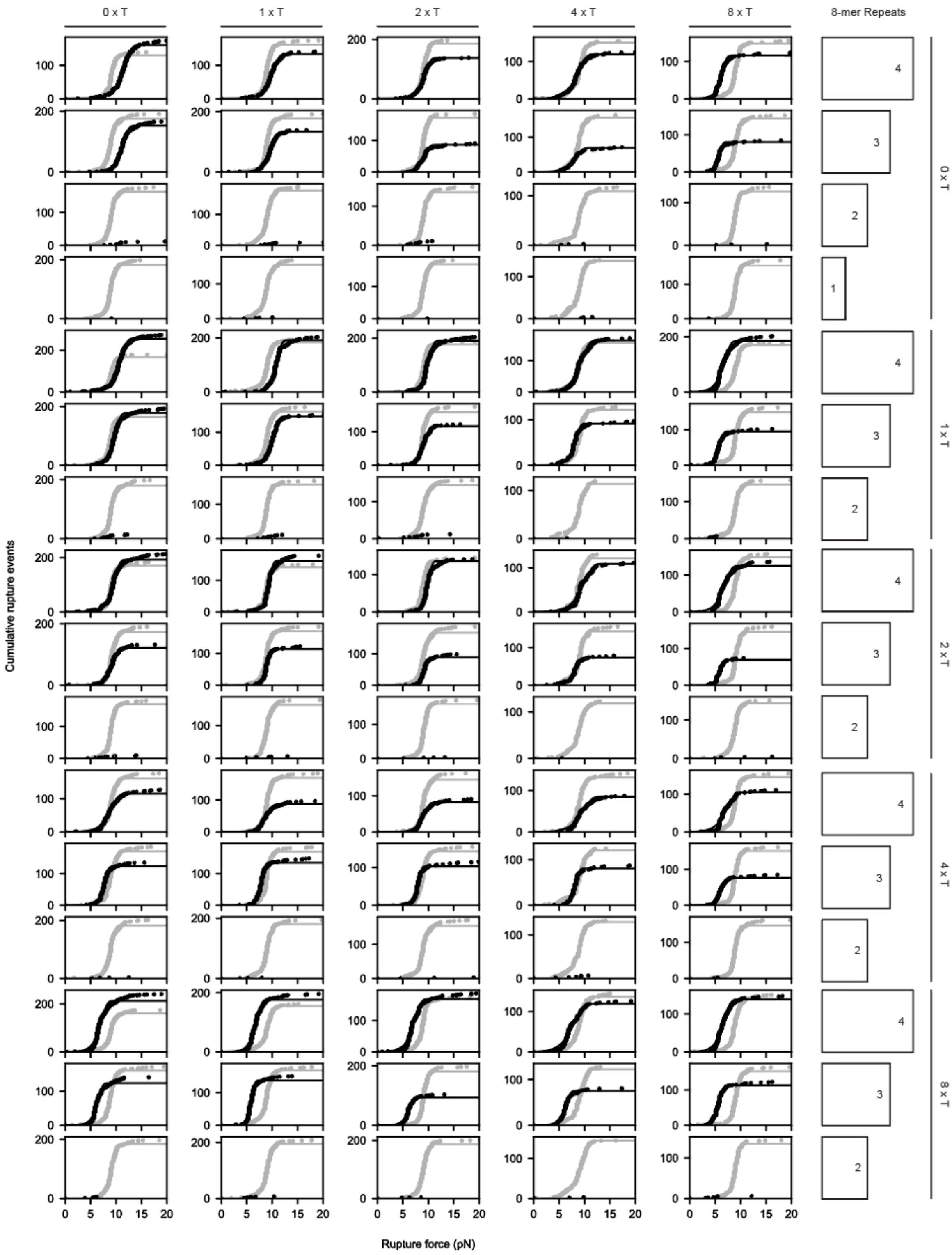


**Figure S36. Cumulative rupture force distributions and fits of 8-mer multivalent duplexes.** Cumulative rupture force distributions for variable duplexes (black data points) and 'fiducial' duplexes (grey data points). Horizontal bars (right) indicate number of duplex repeats. Linker sizes are annotated on the edges of the distributions: right labels indicate length of linker A, and top labels indicate length of linker B. Bell-Evans model fits for all data are shown in black or grey for variable and fiducial duplexes, respectively. The 1 x 8-bp construct corresponds to a single 8-bp unit duplex.

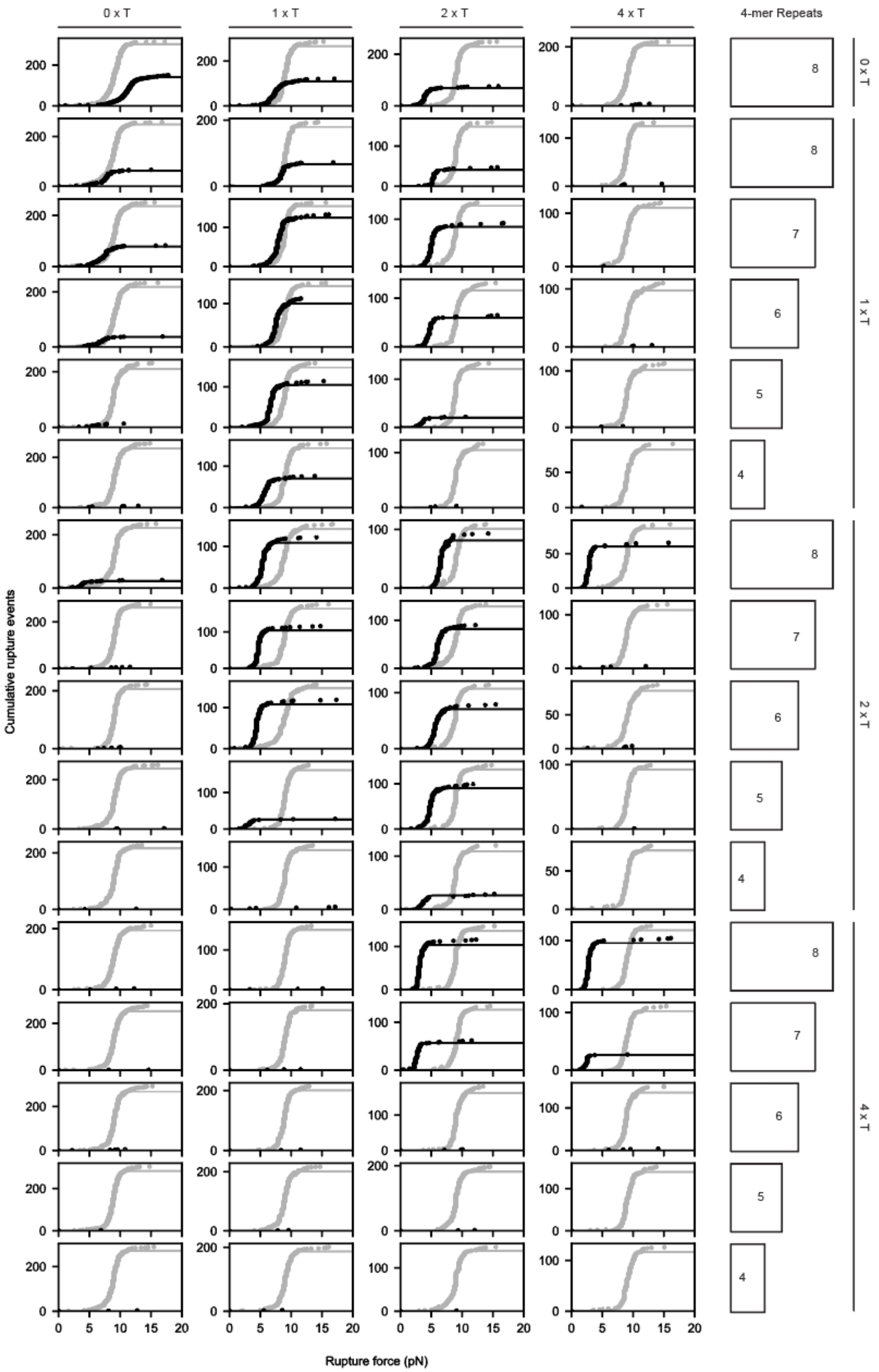


**Figure S37. Cumulative rupture force distributions and fits of 4-mer multivalent duplexes.** Cumulative rupture force distributions for variable duplexes (black data points) and 'fiducial' duplexes (grey data points). Horizontal bars (right) indicate number of duplex repeats. Linker sizes are annotated on the edges of the distributions: right labels indicate length of linker A, and top labels indicate length of linker B. Bell-Evans model fits for all data are shown in black or grey for variable and fiducial duplexes, respectively.

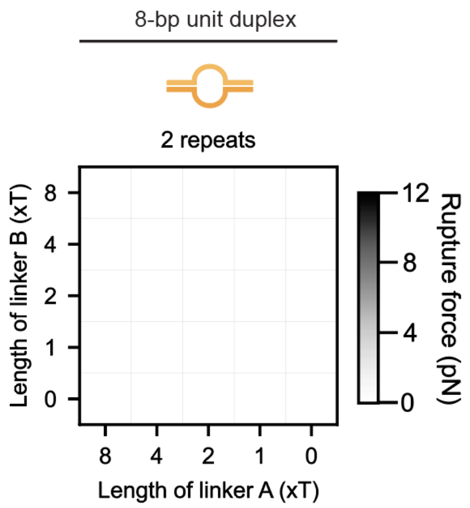


**Figure S38. Rupture force matrix for 2-repeat 8-bp multivalent duplex, which yielded no observed rupture events.** Rupture force matrix for the 2 x 8-bp multivalent duplex. The schematic illustrates the multivalent duplex with the largest, equal-length linkers. Dark grey indicates larger unzipping forces. White indicates no observed rupture events with a rupture force of 0 pN. There were no observed rupture events for the 2 x (8-bp duplex) construct.

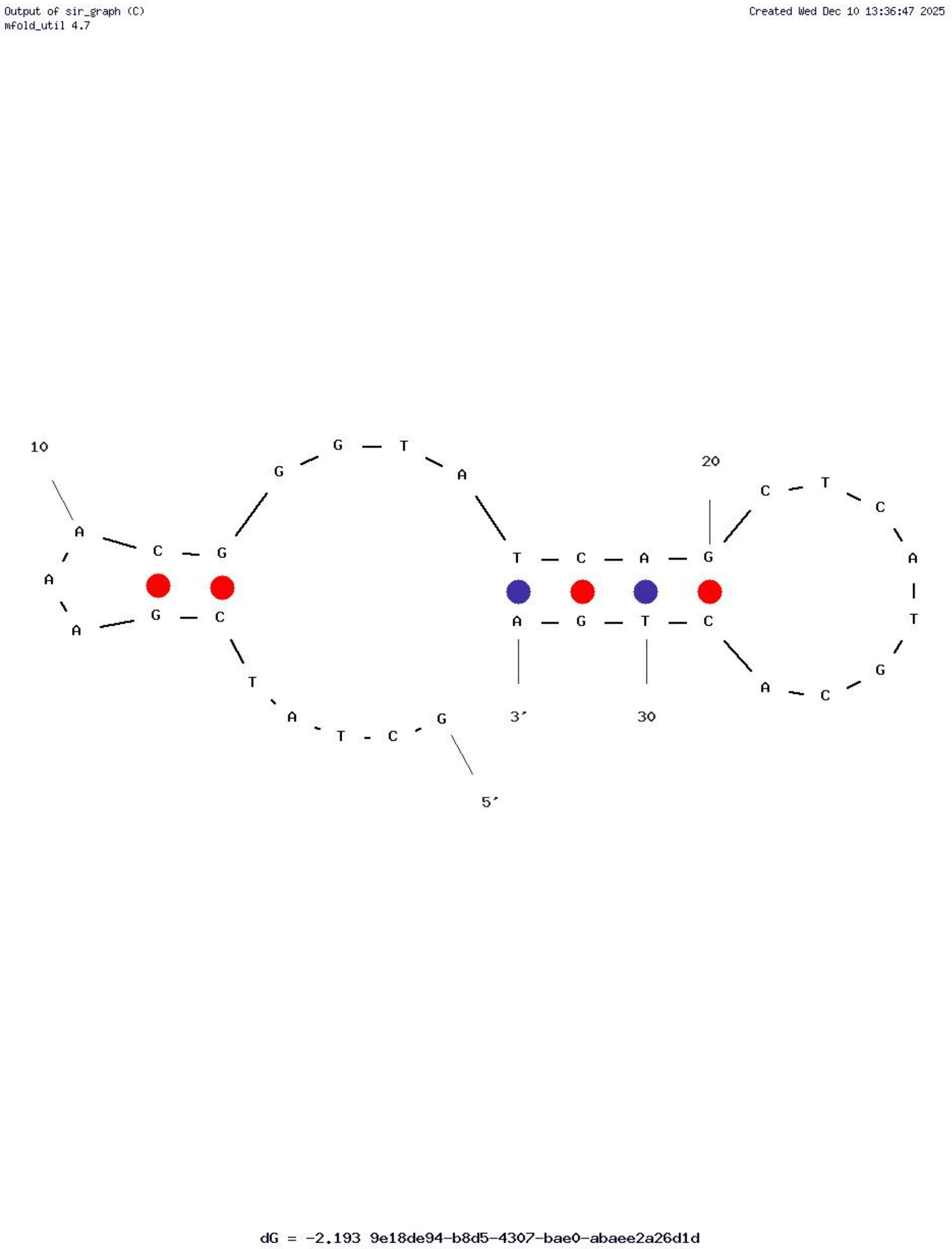


**Figure S39. Predicted secondary structure in the 2 x 8-bp construct.** Secondary structure was predicted with UNAFold at 20ºC in PBS buffer.[13]

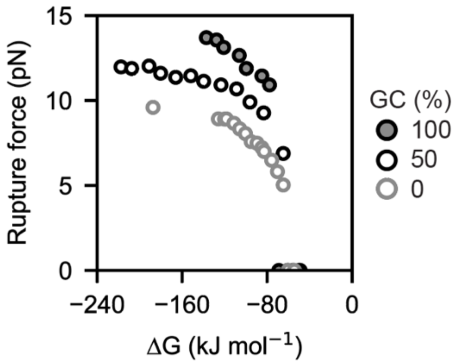


**Figure S40. Unzipping force as a function of free energy of hybridization for short standard duplexes.** Unzipping forces of standard DNA duplexes with 0% (open-face data points with grey borders), 50% (open-face data points with black borders), and 100% (grey data points with black borders) GC content in PBS at 21ºC at a ramp rate of 0.5 pN $s^{-1}$ (**Fig. 5C**). Markers represent mean rupture forces; free energies of hybridization at 21ºC were calculated with DuplexFold.[14]

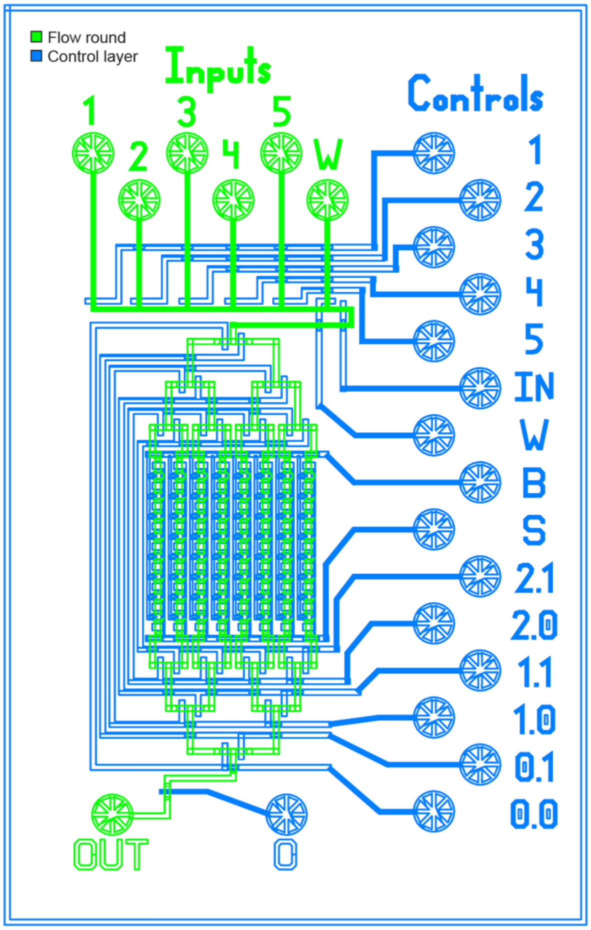


**Figure S41. Schematic of the Minichip device.** Schematic shows different layers of photoresist required to fabricate molding masters for the flow (AZ50 XT, green) and control (SU-8 2025) molds. AutoCAD design files are available as Supplementary Files in a Zenodo online repository.

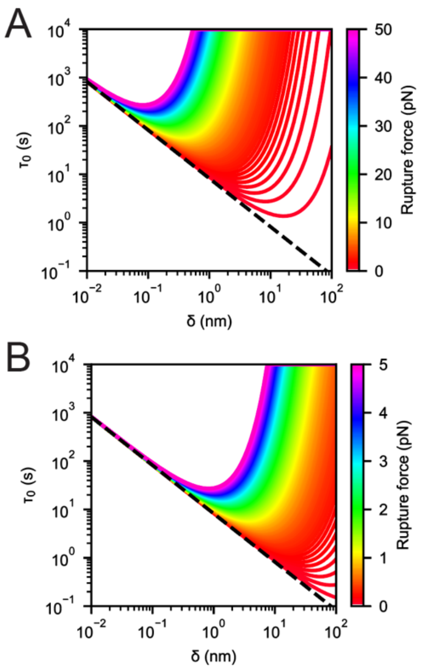


**Figure S42. Analytical solutions of the Bell-Evans model.** Contour map indicating combinations of zero-force bond lifetimes and distances to the transition state for the rupture pathway that yield the same rupture force with a ramp rate of 0.5 pN $s^{-1}$. The black dotted line is the physical limit for a physically defined most probable rupture force, corresponding to $\tau_0 = \frac{k_B T}{\dot{F}\delta}$, where the characteristic time scales of thermal and force-induced dissociation are similar. For $\tau_0 < \frac{k_B T}{\dot{F}\delta}$, the most probable rupture force is predicted to be less than 0 pN. The range of parameters span from 0 to 50 pN in (**A**) and to 5 pN in (**B**).

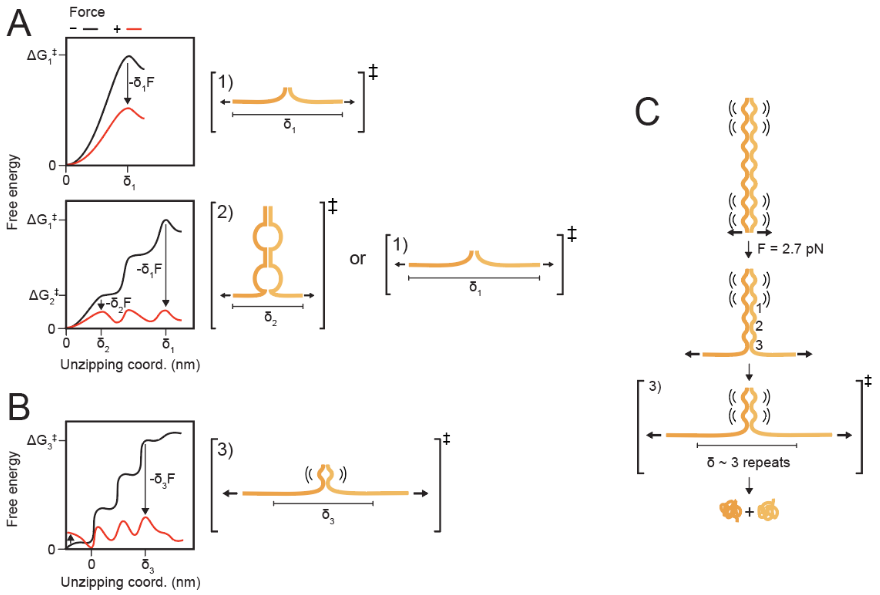


**Figure S43. Free energy diagrams for transition state models. A.** Transition state models of unzipping for standard and multivalent DNA duplexes. The free energy barrier of unzipping is $\Delta G^{\ddagger}$, and the distance to the transition state of unzipping is $\delta$. The free energy barrier decreases with force, F, applied in the direction of the reaction coordinate, by $-F \cdot \delta$. Transition state (1) is a fully unzipped duplex, and transition state (2) is one unzipped unit duplex of the multivalent duplex. **B.** An alternative transition state model of the multivalent duplex with an intermediate number of unzipped unit duplexes in the transition state. In this model, the flanking unit duplexes are less likely to be bound and therefore have a smaller contribution to the free energy of the bound state relative to the interior duplexes. **C.** A schematic reaction of the 8 x (4-bp + 4T) multivalent duplex. At zero force, the duplex is bound, but the flanking unit duplexes are less stable. At the most probable rupture force, the flanking unit duplex experiencing unzipping force is unbound prior to irreversible unzipping. In the transition state of unzipping, 3 to 4 interior duplexes unzip, resulting in irreversible rupture.

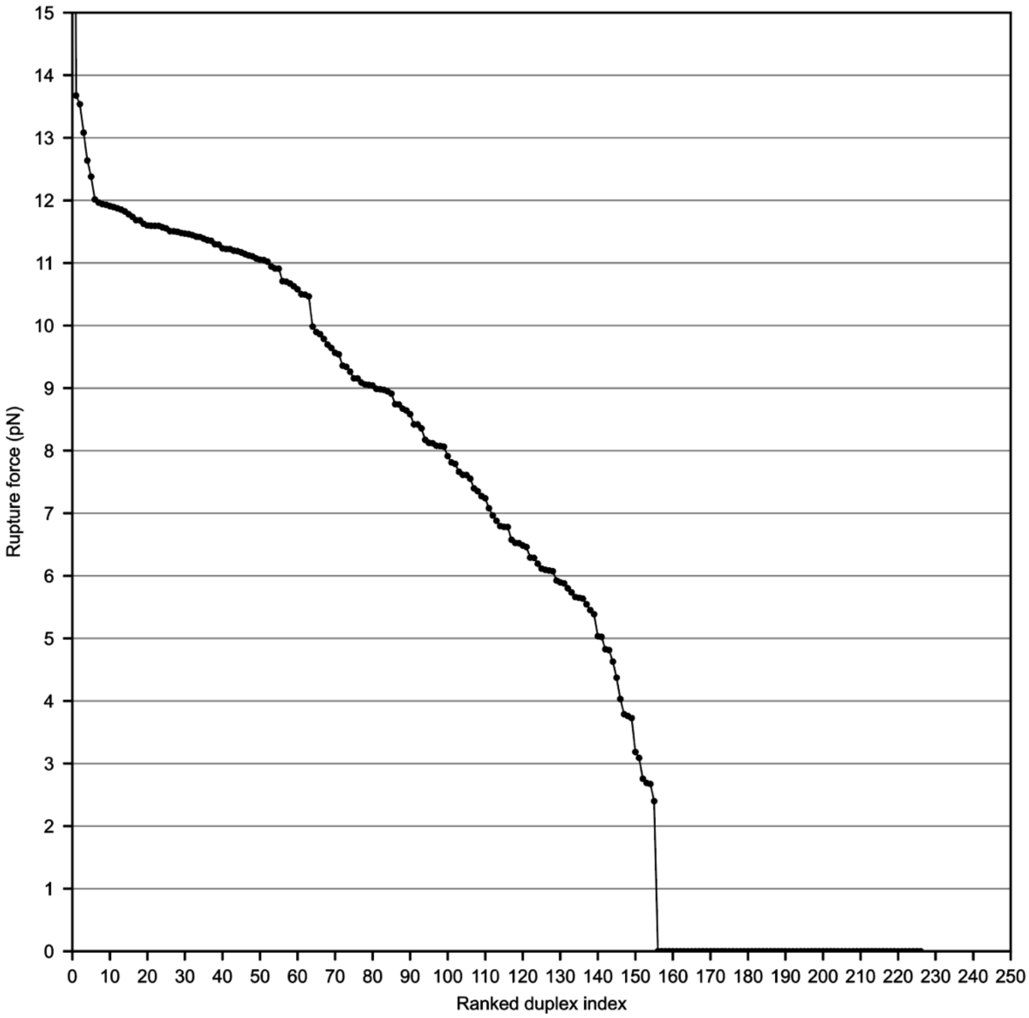


**Figure S44. Ranked unzipping forces from the first and second DNA duplex libraries.** Unzipping forces shown for all measured DNA duplexes. Rupture force measurements were made in PBS at room temperature at a force ramp rate of 0.5 pN $s^{-1}$. Duplexes with a rupture force of 0 pN were likely thermally unstable and had fewer than 20 observations (**Supplementary note 4**, **Figure S34**).

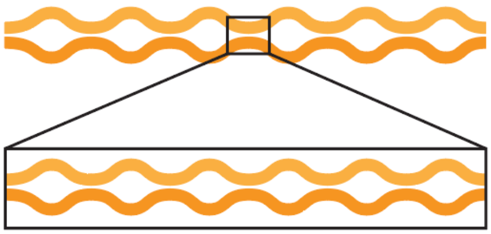

**Figure S45. Fractal multivalent duplex design.** The fractal design is a multivalent duplex in which the unit duplex itself is a multivalent duplex.

| Technology | Conditions per experiment | FOV ($mm^2$) | Resolution (nm) | Rel. error in force (δR/R) | Tracked beads | Single molecules |
|---|---|---|---|---|---|---|
| $SM^3FS$ | 16 | 11 | 20 | 2 | 30,000 | 18,000 |
| Microfluidic force spectroscopy | 1 | 15 | 20 | 2 | 50,000 | 10,000 |
| Centrifuge force spectroscopy | 1 | 0.04 | 2 | 3 | >3,000 | 3,000 |
| Acoustic force spectroscopy | 1 | 0.9 | 5 | 3 | 2,000 | 150 |
| High-throughput magnetic tweezers | 1 | 0.1 | 1 | 3 | 1,000 | 500 |
| Optical tweezers | 1 | - | 0.1 | 0 to 3 | 1 | 1 |

**Table S1. Parameters of bead-based force spectroscopy assays.** Field of view (FOV), spatial resolution, relative error in force ($\delta F_D \; F_D^{-1}$) versus error in bead radius, and measurement throughput for bead-based force spectroscopy assays including microfluidic force spectroscopy,[15,16] centrifuge force spectroscopy,[17,18] acoustic force spectroscopy,[19] high-throughput magnetic tweezers,[20–22] and optical tweezers.[23,24]
